# Vitamin K-dependent γ-carboxylation regulates calcium flux and adaptation to metabolic stress in β-cells

**DOI:** 10.1101/2022.05.13.491370

**Authors:** Julie Lacombe, Kevin Guo, Jessica Bonneau, Denis Faubert, Florian Gioanni, Alexis Vivoli, Sarah M. Muir, Soraya Hezzaz, Vincent Poitout, Mathieu Ferron

## Abstract

Vitamin K is a micronutrient necessary for the γ-carboxylation of glutamic acids. This post-translational modification occurs in the endoplasmic reticulum (ER) and affects secreted proteins. Clinical studies have recently implicated vitamin K in the pathophysiology of diabetes, but the underlying molecular mechanism remains unknown. Here, we show that mouse β-cells lacking γ-carboxylation fail to adapt their insulin secretion in the context of age-related insulin resistance or diet-induced β-cell stress. In human islets, γ-carboxylase expression positively correlates with improved insulin secretion in response to glucose. We identified Endoplasmic Reticulum Gla Protein (ERGP) as a novel γ-carboxylated ER-resident calcium-binding protein expressed in β-cells. Mechanistically, γ-carboxylation of ERGP protects cells against calcium overfilling by diminishing STIM1 and Orai1 interaction and restraining store-operated calcium entry. These results reveal a critical role for vitamin K-dependent γ-carboxylation in the regulation of calcium flux in β-cells and in their capacity to adapt to metabolic stress.

## INTRODUCTION

Type 2 diabetes (T2D) is a metabolic disorder characterized by insulin resistance, hyperglycemia and hyperinsulinemia (Hudish et al., 2019). Traditionally, T2D has been viewed as a disease initiated by peripheral insulin resistance ultimately resulting in pancreatic β-cell exhaustion. However, recent studies suggest that uncontrolled and excessive insulin secretion by β-cells could be the driving force that elicits peripheral insulin resistance and metabolic complications in T2D (Mehran et al., 2012; Mittendorfer et al., 2022). Human and animal studies indicate that several factors can influence T2D susceptibility, including age, genetic variants, and diet. Many of these factors are thought to directly impact β-cell function (Solis-Herrera et al., 2000). Increased consumption of highly processed, calorie-rich but nutrient-poor food is one possible contributor to the current T2D pandemic (Srour et al., 2020). Paradoxically, in Western countries, excess calorie intake is frequently associated with deficiencies in a number of micronutrients, including trace elements such as zinc (Chabosseau and Rutter, 2016), and several vitamins (Kaidar-Person et al., 2008). Other studies have linked micronutrient deficiencies to an increased risk of diabetes (Hoffman et al., 2021; Via, 2012). Yet, the role of micronutrients in β-cell function remains poorly understood.

Vitamin K (VK), a fat-soluble vitamin, functions as a co-factor during the γ-carboxylation reaction that converts glutamic acid (Glu) residues to γ-carboxyglutamic acid (Gla) residues in proteins transiting through the endoplasmic reticulum (ER). Two ER-resident enzymes are involved in this reaction, which together form the VK cycle: γ-glutamyl carboxylase (GGCX), and vitamin K oxidoreductase (VKORC1) (Lacombe and Ferron, 2018). GGCX requires reduced VK (VKH_2_) as an essential cofactor, which upon carboxylation of proteins, is oxidized to VK epoxide (VKO) and then reconverted to VKH_2_ by VKORC1. The presence of this post-translational modification in proteins results in higher affinity for calcium ions (Ca^2+^). Altogether, in vertebrates, less then fifteen γ-carboxylated proteins have been identified so far, all of them being secreted proteins. Gamma-carboxylation is essential in the liver for the activity of several coagulation factors (e.g., prothrombin, factor IX, etc.), and in arteries and cartilage to modulate the activity of Matrix Gla Protein (MGP), which prevents extra-osseous tissue mineralization (Furie et al., 1999; Murshed et al., 2004). Gamma-carboxylation also negatively regulates the function of osteocalcin, a bone-derived hormone with pleiotropic actions (Ferron et al., 2015; Lee et al., 2007). Whether γ-carboxylation occurs on ER-resident proteins and regulates cellular functions in a cell-autonomous manner is currently unknown.

Clinical and genetic data suggest that VK insufficiency or reduced VK intake are associated with an increased risk of developing metabolic syndrome or T2D (Pan and Jackson, 2009). Two longitudinal studies found a positive association between low dietary VK intake and risk of developing T2D (Beulens et al., 2010; Ibarrola-Jurado et al., 2012). It was also observed that 40% of morbidly obese patients are characterized by VK insufficiency and that low serum VK correlates positively with the presence of T2D in these subjects (Dihingia et al., 2018; Ewang-Emukowhate et al., 2015; Zwakenberg et al., 2019). Finally, VK supplementation in patients with T2D significantly decreased their fasting glucose and HbA1c blood concentrations (Karamzad et al., 2020; Rahimi Sakak et al., 2021). These clinical studies suggest a link between VK insufficiency and the risk of developing diabetes. However, they also raise important questions regarding the mechanism by which VK protects against T2D. Does VK directly affect β-cell function? If so, what are the cellular and molecular mechanisms involved, and which γ-carboxylated protein mediates the protective effect of VK?

In the current study, we aimed at answering these questions using a combination of unique genetic, cellular, and biochemical tools we developed to study γ-carboxylation. We show that GGCX and VKORC1, the two enzymes of the VK cycle, are expressed and active in mouse and human pancreatic islets and β-cells. Using loss-of-function models, we found that the inactivation of *Ggcx* impairs β-cell function in young mice exposed to a short bout of high-fat diet (HFD), and compromises β-cell survival in older animals fed a regular diet. Finally, we identify <u>E</u>ndoplasmic <u>R</u>eticulum <u>G</u>la <u>P</u>rotein (ERGP), as a novel γ-carboxylated ER-resident calcium-binding protein that prevents calcium overfilling by regulating store-operated calcium entry (SOCE) in β-cells. These data demonstrate that γ-carboxylation plays a critical role in the capacity of β-cells to adapt to physiological stress and describe a new cellular function for this post-translational modification.

## RESULTS

### Vitamin K-dependent carboxylation occurs in islets and β-cells

To identify tissue(s) involved in the beneficial effect of VK on glucose metabolism and T2D, we first examined GGCX and VKORC1 protein levels in a panel of human tissues using the ProteomicsDB resource (Schmidt et al., 2018). This analysis revealed that pancreatic islets were ranked fourth and first for GGCX and VKORC1 protein expression respectively (Fig. 1A), in agreement with our own data showing that *Ggcx* and *Vkorc1* genes are highly expressed in mouse islets (Fig. S1A-B). To more precisely dissect *Ggcx* and *Vkorc1* expression within mouse islets, we used fluorescence-activated cell sorting (FACS) to isolate β-cells based on the expression of the fluorescent reporter protein tdTomato (Tom) conditionally expressed in the insulin-positive cells of *Ins1^Cre/+^*; *Rosa26^CAG-lox-stop-lox-tdTomato^* mice. Using this strategy, we obtained a pure β-cell population (Tom+), as demonstrated by the expression of the insulin genes *Ins1* and *Ins2,* and the absence of the other endocrine cell type markers *Gcg*, *Ppy* and *Sst*, which were highly expressed in the Tom-population (Fig. 1B). Further quantitative PCR (qPCR) analyses revealed that *Ggcx* and *Vkorc1* are expressed in β-cells and other islet endocrine cells (Fig. 1C). *Tabula Muris* single cell transcriptomics data (Consortium, 2018) confirmed *Ggcx* and *Vkorc1* expression in the endocrine pancreas, whereas very few pancreatic exocrine cells express these genes (Fig. S1C-D). These results agree with another set of publicly available mouse islet transcriptomic data (DiGruccio et al., 2016). In addition, we found that GGCX and VKORC1 proteins are expressed at similar and higher levels respectively in purified β-cells as compared to whole islets (Fig. 1D).

**Figure 1:**
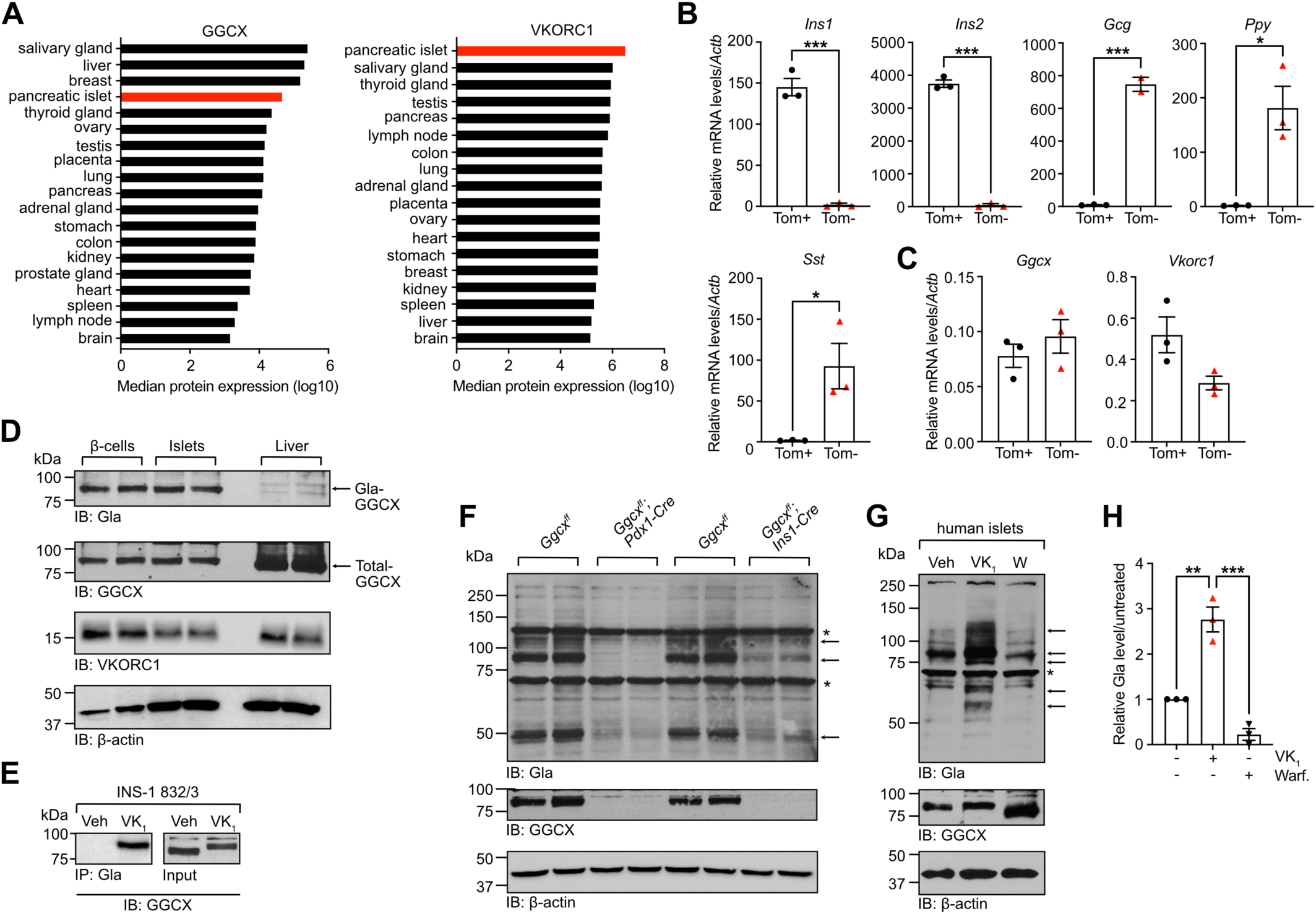
Vitamin K-dependent γ-carboxylation machinery is active in islets and β-cells. **(A)** GGCX and VKORC1 protein abundance in various human tissues expressed as normalized intensity based absolute quantification (iBAQ; www.proteomicsdb.org). **(B-C)** Dispersed islet cells from *Ins1^Cre/+^*; *Rosa26^CAG-lox-stop-lox-tdTomato^*mice were sorted by flow cytometry based on tdTomato expression (Tom+ versus Tom-) and gene expression analysed by quantitative PCR and normalized to *Actb* (n=3). **(D)** Protein expression in β-cells, islets and livers from *Ins1^Cre/+^*; *Rosa26^CAG-lox-stop-lox-tdTomato^*mice was measured by western blot using anti-Gla, anti-GGCX and anti-VKORC1 antibodies. β-Actin was used as a loading control. **(E)** INS-1 832/3 cells were cultured in presence of vitamin K_1_ (VK_1_; 22μM) or vehicle for 3 days and GGCX γ-carboxylation was assessed by anti-Gla immunoprecipitation followed by western blot analysis with anti-GGCX antibodies. **(F)** Islets from *Ggcx^ff^, Pdx1-Cre* and *Ggcx^ff^; Ins1-Cre* mice and their respective *Ggcx^ff^*littermates were harvested and γ-carboxylation and GGCX expression were analysed by western blot using anti-Gla and anti-GGCX antibodies. β-actin was used as a loading control. Arrows indicate γ-carboxylated proteins, while asterisks indicate non-specific binding. **(G-H)** Human islets from non-diabetic cadaveric donors were cultured in presence of VK_1_ (22μM), warfarin (50μM) or vehicle for 3 days and both γ-carboxylation and GGCX expression were analysed by western blot using anti-Gla and anti-GGCX antibodies. β-actin was used as a loading control. **(G)** Representative western blot experiment with islets from donor R266. **(H)** Gamma-carboxylation was quantified using arbitrary densitometry units of Gla signals over β-actin signals. Data from VK_1_ and warfarin treated samples were normalized over vehicle treatment (n=3). Results represent the mean ± SEM; Unpaired, 2-tailed Student’s *t* test was used in (B-C); Ordinary one-way ANOVA with Bonferroni’s post-tests was used in (H); \*\*\**P* < 0.001; \*\**P* < 0.01; \**P* < 0.05.

Previous studies have established that GGCX carboxylates itself in a VK-dependent manner in vitro and in vivo in perinatal liver (Berkner and Pudota, 1998; Lacombe et al., 2018). Using γ-carboxylated GGCX (Gla-GGCX) as a readout of a functional VK cycle, we demonstrated that carboxylation does take place in pancreatic islets and specifically in β-cells (Fig.1D). GGCX expression was also detected in the rat insulinoma cell line INS-1 832/3 and its γ-carboxylation was induced following treatment with phylloquinone (vitamin K_1_; VK_1_), which is absent from cell culture media and fetal bovine serum (Haque et al., 2014) (Fig.1E). To determine the extent of γ-carboxylation in vivo in islets and β-cells, we isolated islets from *Ggcx^ff^; Pdx1-Cre* and *Ggcx^ff^; Ins1-Cre* mice in which *Ggcx* has been inactivated specifically in the pancreas or in β-cells respectively (Fig. 1F and Fig. S1E-F). Western blot analyses with a previously characterized α-Gla specific antibodies (Lacombe et al., 2018) revealed the presence of carboxylated proteins in islets and β-cells as demonstrated by the reduced α-Gla immunoreactivity in *Ggcx^ff^; Pdx1-Cre* and *Ggcx^ff^; Ins1-Cre* islets (Fig.1F). Finally, GGCX expression was also detected in human islets and culturing them with VK_1_ significantly increased protein γ-carboxylation.

Conversely, warfarin, an inhibitor of VK oxidoreductase activity (Shen et al., 2017), reduced it (Fig. 1G-H, Table S1). We also observed that in the presence of warfarin, GGCX migrates faster on SDS-PAGE because of its incomplete γ-carboxylation, in agreement with previous studies (Berkner and Pudota, 1998; Lacombe et al., 2018). Together, these data support the conclusion that VKORC1, GGCX and γ-carboxylated proteins are present in islets and β-cells.

### Loss of γ-carboxylation induces a diabetic signature in islets

As a first step to determine the role of VK-dependent carboxylation in islets and β-cells, we analyzed the expression profile of *Ggcx^ff^; Pdx1-Cre* islets by RNA-sequencing (RNAseq). In comparison to control *Ggcx^ff^* islets, we found that 319 genes were differentially expressed in *Ggcx^ff^; Pdx1-Cre* islets (adjusted *P* value ≤ 0.05; Table S2). We divided this set of genes into two groups based on whether their expression was increased (114 genes) or decreased (205 genes) following *Ggcx* inactivation in islets, then completed a series of bioinformatics analyses on each group. Gene set enrichment analyses with Gene Ontology revealed that many biological processes implicated in the response to ER stress were significantly enriched within the group of genes downregulated in *Ggcx^ff^; Pdx1-Cre* islets, such as *ER-nucleus signaling pathway*, *positive regulation of response to ER stress*, *regulation of response to ER stress*, *I-kappaB kinase/NF-kappaB signaling* and *regulation of apoptotic signaling pathway* (Fig.S2A). Similarly, when we interrogated the KEGG pathway database, we found that this group of genes was enriched for pathways such as *apoptosis*, *protein processing in ER* and *NF-kappa B signaling pathway* (Fig.S2B). Using the UniProt annotated keywords database, we observed, in the group of genes that were upregulated in *Ggcx^ff^; Pdx1-Cre* islets, a very strong enrichment for protein keywords related to the secretory pathway (e.g., *glycoprotein, signal, secreted, disulfide bond* and *extracellular matrix*; Fig. S2C). Based on these observations, we hypothesized that the capacity of islet cells to respond and adapt to ER stress might be dysregulated in the absence of γ-carboxylation, potentially leading to impaired β-cell function. In support of this notion, we found that a network of genes previously implicated in the β-cell response to ER stress, including *Ddit3* (CHOP), *Atf4*, *Eif2ak3* (PERK)*, Herpud1*, *Trib3*, *Pdia4*, *Ppp1r1a* and *Atp2a2* (SERCA2) (Johnson et al., 2014; Sharma et al., 2021), was down-regulated in the absence of γ-carboxylation (Fig. S2D).

We next determined to which extent the transcriptome of *Ggcx^ff^; Pdx1-Cre* islets intersects with the gene expression profile of islets from pre-diabetic (adult C57BL/6 mice on HFD for 8 weeks) or diabetic (8-weeks old *Lepr^db/db^*and 7-weeks old *Ire1α ^ff^; Ins2-Cre^ERT/+^*) mouse models (Lee et al., 2020; Motterle et al., 2017; Wang et al., 2012). We found that 75% of the up- and 45% of the down-regulated genes in *Ggcx^ff^; Pdx1-Cre* islets were similarly dysregulated in at least one of these mouse models (Fig. S2E). About one-third of these genes encode proteins found within the secretory pathway (Fig.S2F). The overlap between the transcriptome of *Ggcx^ff^; Pdx1-Cre* islets and each mouse model was statistically significant for all comparisons and the highest significances were found for the comparison with islets from the diabetic mouse models (Table S3). Overall, these data suggest that loss of function of *Ggcx* in pancreatic endocrine cells induces a diabetic gene signature in these cells, presumably by altering their capacity to respond to ER stress.

### GGCX is necessary for the maintenance of an adequate β-cell mass in adult mice

To determine the role of VK-dependent carboxylation in islet function in vivo, we next analyzed the metabolic consequences of a pancreas-specific inactivation of *Ggcx* (*Ggcx^ff^; Pdx1-Cre* mice). The *Pdx1-Cre* driver was selected because it resulted in efficient deletion in pancreatic islets (Ferdaoussi et al., 2015), without expressing the human growth hormone (hGH), which was found to be present in several β-cell-specific Cre transgenes (*RIP-Cre*, *MIP-Cre^ERT^*, etc.) and to affect β-cell function and proliferation (Brouwers et al., 2014; Oropeza et al., 2015). *Ggcx* mRNA level was reduced by >90% and GGCX protein was undetectable in the pancreatic islets of these mice (Fig. S1E and Fig. 1F). In agreement with efficient inactivation of GGCX, protein γ-carboxylation was abrogated in *Ggcx^ff^; Pdx1-Cre* islets (Fig. 1F). When compared to control littermates these mice did not display any differences in energy expenditure parameters (energy expenditure, O_2_ consumption, CO_2_ release), physical activity, food intake, pancreas weight and body weight (Fig. S3A-E). In addition, inactivation of *Ggcx* occurred only in the pancreas of the *Ggcx^ff^; Pdx1-Cre* mice and not in any other tissue tested, including the hypothalamus and other parts of the brain (Fig. S3F).

Glucose tolerance test (GTT) revealed that an absence of γ-carboxylation in islets does not affect glucose handling in 16-week-old mice (Fig. 2A). However, at 24 weeks of age, *Ggcx^ff^; Pdx1-Cre* mice showed significantly elevated fasting blood glucose and decreased glucose tolerance (Fig. 2B). This defect could be traced to reduced circulating insulin following injection of glucose compared to control (Fig. 2C). Insulin sensitivity, assessed by an insulin tolerance test, appeared to be unaffected (Fig. 2D). Pancreas immunohistochemistry revealed that β-cell area and mass were reduced in *Ggcx^ff^; Pdx1-Cre* mice at 32 weeks but not at 12 weeks of age (Fig. 2E). Accordingly, total insulin content was diminished in the pancreas of 24-28 weeks old *Ggcx^ff^; Pdx1-Cre* mice (Fig. 2F). Beta-cell area and β-cell mass were also significantly lower in mice lacking the two vitamin K oxidoreductases, *Vkorc1* and *Vkorc1l1*, in the pancreas only (Fig. S3G-J), confirming the implication of the VK-cycle in the observed phenotype.

**Figure 2:**
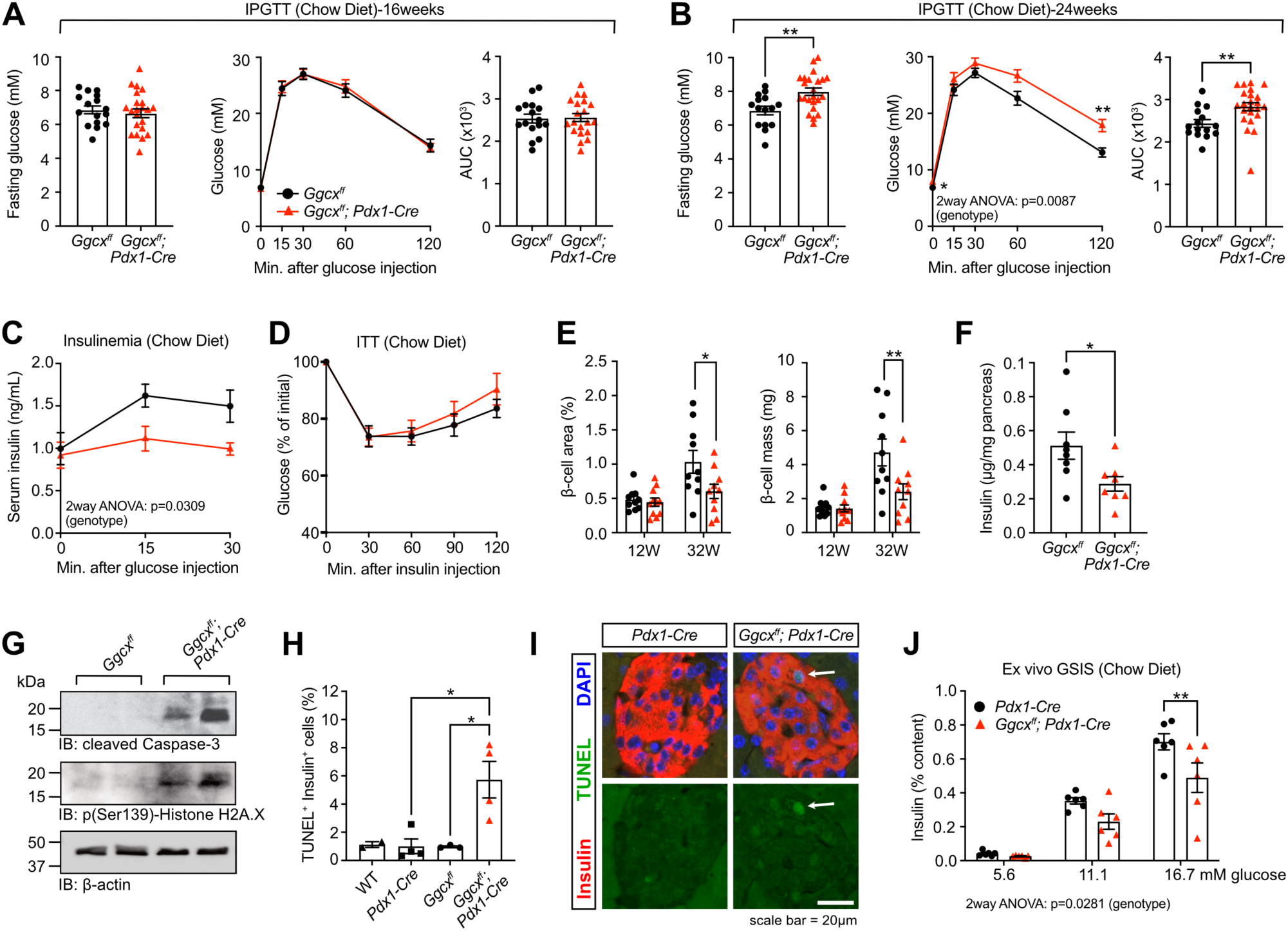
GGCX is necessary for the maintenance of an adequate β-cell mass in adult mice. **(A-B)** Intraperitoneal glucose tolerance test (IPGTT) with *Ggcx^ff^; Pdx1-Cre* and *Ggcx^ff^* male mice was performed following an over-night fast and intra-peritoneal injection of a bolus of glucose (2g/kg of body weight). Blood glucose was analyzed at fasting and at 15-, 30-, 60- and 120-minutes post-injection in **(A)** 16-weeks-old (n=16-21) and **(B)** 24-weeks-old (n=15-23) mice. **(C)** The insulinemic response to glucose was measured after an over-night fast and intra-peritoneal injection of a bolus of glucose (3g/kg of body weight) in 24-weeks-old male mice (n=10). Serum was collected at fasting and 15- and 30-minutes post-injection and insulin concentration measured by ELISA. **(D)** Insulin tolerance test (ITT) was performed following a 5h fast and intra-peritoneal injection of insulin (1U/kg) in 24-weeks-old male mice (n=9-10). **(E)** Histomorphometric analysis on pancreas section following insulin staining and hematoxylin counterstaining from 12- and 32-weeks-old mice (n=10-11). **(F)** Pancreas from 24- to 28-weeks old *Ggcx^ff^; Pdx1-Cre* and *Ggcx^ff^* male mice were homogenized and insulin content measured by ELISA (n=8). **(G)** The presence of cleaved-caspase-3 and p(Ser139)-Histone H2A.X in *Ggcx^ff^; Pdx1-Cre* and *Ggcx^ff^* islets was analysed by western blotting. β-Actin was used as a loading control. **(H)** β-cell specific apoptosis was detected by TUNEL and insulin co-staining on pancreas sections from 32-weeks old mice. **(I)** Representative picture showing TUNEL^+^ Insulin^+^ β- cells in pancreas section from *Ggcx^ff^; Pdx1-Cre* mice (indicated by the white arrow). **(J)** Ex vivo glucose stimulated insulin secretion (GSIS) test on isolated islets from *Pdx1-Cre* and *Ggcx^ff^; Pdx1-Cre* mice. Secreted insulin was normalized over islet insulin content (n=6, with 10 islets each). Results represent the mean ± SEM; Unpaired, 2-tailed Student’s *t* test was used in (A-B) for fasting glucose and AUC and in (F); Two-way ANOVA with Bonferroni’s post-tests was used in (A-B) for IPGTTs and in (C-E) and (J); Ordinary one-way ANOVA with Bonferroni’s post-tests was used in (H); \*\*\**P* < 0.001; \*\**P* < 0.01; \**P* < 0.05.

By western blot using antibodies against cleaved-caspase-3 and phospho(Ser139)-Histone H2A.X, we detected apoptosis and DNA damage in >32-weeks old *Ggcx^ff^; Pdx1-Cre* islets, but not in *Ggcx^ff^*controls (Fig. 2G). Beta-cell specific apoptosis was independently confirmed using TUNEL and insulin co-staining on pancreas sections (Fig. 2H-I). To rule out the possibility that the *Pdx1-Cre* transgene itself was responsible for the phenotype observed in *Ggcx^ff^; Pdx1-Cre* mice, β-cell apoptosis, β-cell mass and pancreas insulin content were analyzed in *Pdx1-Cre* mice. None of these parameters were affected by the presence of the Cre recombinase (Fig. 2H and Fig. S3K-L).

The role of γ-carboxylation in insulin secretion was next assessed directly by performing ex vivo glucose stimulated insulin secretion (GSIS) test on GGCX-deficient (*Ggcx^ff^; Pdx1-Cre*) and control (*Pdx1-Cre*) islets incubated with increasing concentrations of glucose (5.6mM, 11.1mM, and 16.7mM). These static ex vivo GSIS experiments revealed a significant reduction in the capacity of the *Ggcx*-deficient islets to secrete insulin in response to glucose (Fig. 2J). These results were obtained using pools of similarly sized islets and were normalized to insulin content, suggesting that γ-carboxylation is not only necessary to maintain a proper β-cell mass, but is also required for an adequate insulin secretion in response to glucose in β-cells.

### Pancreas or β-cell specific deletion of *Ggcx* compromises insulin secretion in response to high fat diet

Since the phenotype of the *Ggcx^ff^; Pdx1-Cre* mice appears to be age-dependent, we hypothesized that GGCX activity would be predominantly required when β-cells need to adapt to stress such as age-related insulin resistance. To more directly test GGCX involvement in acute β-cell stress response, 10-week-old *Ggcx^ff^; Pdx1-Cre* mice, which had not developed metabolic and β-cell mass phenotypes yet (Fig. 2A and E), were fed a high-fat diet (HFD; 60% kcal from fat) or a matched control low-fat diet (10% kcal from fat) for 7 days. Previous studies have established that one week of HFD feeding in wildtype mice was sufficient to induce β-cell ER stress, glucose intolerance and hyperinsulinemia without significantly affecting peripheral insulin sensitivity (Sharma et al., 2015; Stamateris et al., 2013). qPCR analysis of ER-stress markers (*spliced Xbp1*, *Ddit3, Ppp1r15a*, *Syvn1, Hspa5* and *Edem1*) on isolated islets confirmed that this short bout of HFD induces ER-stress in both control and *Ggcx^ff^; Pdx1-Cre* islets (Fig. S4A-F). Interestingly, *Ddit3* (CHOP), *Ppp1r15a* (GADD34) and *Hspa5* (Grp78/BiP) were significantly more elevated following short HFD feeding in the *Ggcx^ff^; Pdx1-Cre* islets, while *sXbp1* was lower, suggesting a dysregulated unfolded protein response (UPR) in the absence of γ-carboxylation. This 7-day HFD feeding was also enough to increase body weight in mice, regardless of the presence of *Ggcx* in their islets (Fig. S4G). However, in contrast to control animals, mice deprived of *Ggcx* expression in islets were not able to maintain their fed blood glucose level following HFD (Fig. 3A). The insulinemic response to glucose in absolute value or expressed as a stimulation index (SI: blood insulin concentration at 15 minutes or 30 minutes over T0) was not affected in the absence of *Ggcx* in mice fed the control diet (Fig. 3B-C), in agreement with the fact that glucose handling was not altered in these mice at 12 weeks of age when fed a regular chow diet (Fig. 2A). In contrast, following 7 days on HFD, *Ggcx^ff^; Pdx1-Cre* mice showed a strong suppression in SI, which was significantly lower than the SI of *Pdx1-Cre* control mice (Fig. 3D-E). Of note, this impaired SI was associated with elevated fasting insulin (Fig. 3F), while fasting glucose was not reduced and glucose tolerance was moderately impaired (Fig. S4H).

**Figure 3:**
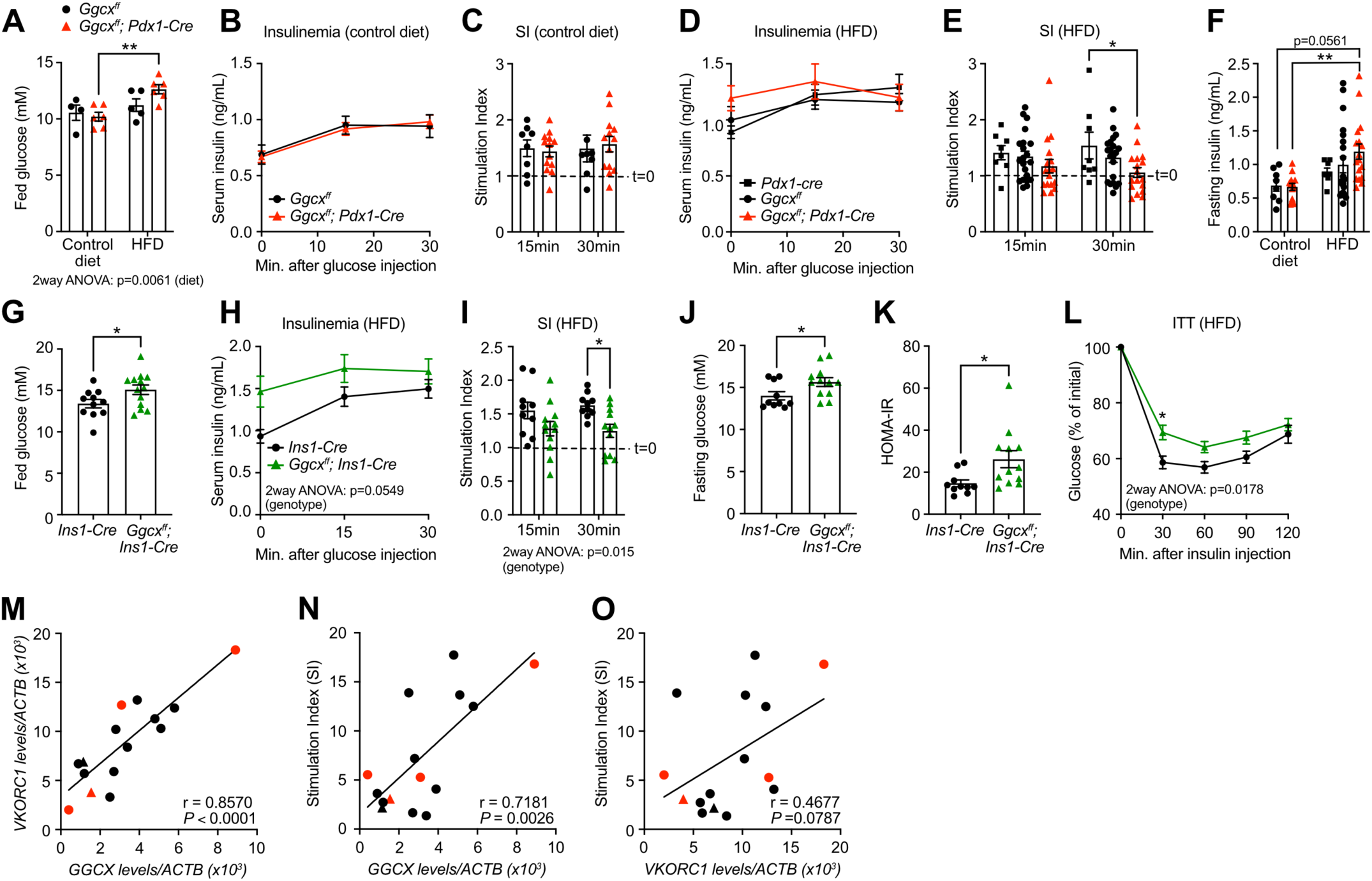
Pancreas or β-cell specific deletion of *Ggcx* compromises insulin secretion in response to high fat diet. **(A-L)** Ten-weeks old male mice of different genotypes were fed with a high-fat (60%) or a control low-fat (10%) diet for 7 days and metabolic analysis performed. **(A)** Fed blood glucose level for *Ggcx^ff^; Pdx1-Cre* and *Ggcx^ff^* male mice (n=4-6). **(B-C)** For the control low-fat diet fed *Ggcx^ff^; Pdx1-Cre* and *Ggcx^ff^* mice, the insulinemic response to glucose was measured after a 5-hour fast and intra-peritoneal injection of a bolus of glucose (2g/kg of body weight) (n=8-13). Data are represented in **(B)** absolute value and **(C)** as stimulation index (blood insulin concentration at 15 or 30 minutes over T0). The dashed line represents a stimulation index of 1 at fasting. **(D-F)** For the high-fat diet fed *Ggcx^ff^; Pdx1-Cre*, *Ggcx^ff^* and *Pdx1-Cre* mice, the insulinemic response to glucose is represented in **(D)** absolute value or **(E)** as stimulation index, and **(F)** fasting insulin levels are shown (n=8-21). **(G-L)** Metabolic analysis of *Ggcx^ff^; Ins1-Cre* and *Ins1-Cre* mice following 7 days HFD feeding. **(G)** Fed blood glucose, **(H-I)** insulinemic response to glucose, **(J)** fasting blood glucose, **(K)** HOMA-IR and **(L)** ITT (5h fasting; Insulin 0.75U/kg) were measured (n=10-13). **(M-O)** Correlation in 15 human islet donor samples between **(M)** *Ggcx* and *Vkorc1* gene expression levels, and between **(N)** *Ggcx* or **(O)** *Vkorc* and each sample’s stimulation index (insulin secretion at 10mM over 1mM glucose). Data were normalized using *Actb* and association were analyzed using Pearson’s correlation. Black circles represent non-diabetic male donors, red circles diabetic male donors, black triangle non-diabetic female donor and red triangle diabetic female donor. Results represent the mean ± SEM; Two-way ANOVA with Bonferroni’s post-tests was used in (A-E), (H-I) and (L); Ordinary one-way ANOVA with Bonferroni’s post-tests was used in (F); Unpaired, two-tailed Student’s *t* test was used in (G), (J) and (K); Pearson’s correlation was used in (M-O); \*\**P* < 0.01; \**P* < 0.05.

To determine if GGCX affects β-cell function in a cell-autonomous manner, we next analyzed *Ggcx^ff^; Ins1-Cre* mice. At 10 weeks of age, *Ggcx^ff^; Ins1-Cre* mice maintained on a regular chow diet had normal glucose tolerance, fasting glucose and fasting insulin (Fig. S4I-K). When *Ggcx^ff^; Ins1-Cre* mice were fed a HFD for 7 days, no difference in body weight was noted (Fig. S4L), but their fed glucose level was significantly increased (Fig. 3G) and their SI was reduced (Fig. 3H-I) in comparison to *Ins1-Cre* control mice fed the same HFD. Remarkably, in the same animals, fasting glucose was significantly increased (Fig. 3J), although fasting serum insulin was also increased (Fig. 3H). Glucose tolerance assessed through intraperitoneal or oral glucose load was unaffected (Fig. S4M-N), suggesting a compensatory increase in insulin-independent glucose disposal (Alquier and Poitout, 2018). Fasting hyperglycemia associated with hyperinsulinemia suggests peripheral insulin resistance in these animals. Accordingly, the homeostatic model assessment for insulin resistance (HOMA-IR) was increased in *Ggcx^ff^; Ins1-Cre* mice on HFD (Fig. 3K), and insulin sensitivity assessed by an insulin tolerance test (ITT) was reduced in the same animals (Fig. 3L). Together, these observations indicate that the absence of γ-carboxylation directly impacts β-cells’ capacity to adapt their insulin secretion in the face of metabolic stress, resulting in increased fasting insulin, loss of glucose-stimulated insulin secretion and reduced peripheral insulin sensitivity.

To relate these findings to humans, we analyzed *GGCX* and *VKORC1* gene expression in human islets from 15 non-diabetic and diabetic donors and observed that the level of these two enzymes vary widely between donors, but nevertheless strongly correlate with one another (Fig. 3M and Table S1). This observation suggests that for certain individuals, the γ-carboxylation machinery in their β-cells might be more active compared to others. Further analysis revealed that *GGCX* expression levels positively and significantly correlate with the capacity of islets to secrete insulin in response to glucose (r=0.7181; *P*=0.0026; Fig. 3N). The levels of *VKORC1* also positively correlate with the same parameter although it did not reach statistical significance (r=0.4677; *P*=0.0787; Fig. 3O). These results imply that γ-carboxylation could also impact glucose-stimulated insulin secretion in human β-cells.

### Vitamin K attenuates apoptosis induced by ER calcium depletion

The results presented so far suggest that GGCX is required for β-cells to adapt to metabolic stress in vivo. To determine if VK and γ-carboxylation can protect β-cells from the acute effects of ER stress, INS-1 832/3 β-cells were cultured for 24h in media containing 25mM glucose in the presence or absence of thapsigargin, an inhibitor of the sarco/endoplasmic reticulum Ca^2+^-ATPase (SERCA), and a pharmacological inducer of ER stress (Sharma et al., 2015). Doses of thapsigargin (10-40nM) inducing moderate to elevated cell death in INS-1 cell cultures were selected based on previously published data (Weldemariam et al., 2022). Induction of Grp78/BiP and phospho(Ser724)-IRE following thapsigargin treatment confirmed the presence of ER stress and UPR (Fig. S5A-B). In addition, western blots using antibodies against cleaved-caspase-3 and phospho(Ser139)-Histone H2A.X showed that thapsigargin dose-dependently induced apoptosis and DNA damage in β-cells, while pre-treatment with VK_1_ reduced the deleterious effects of 10 and 20nM of thapsigargin (Fig. 4A). Interestingly, in the presence of thapsigargin, GGCX protein level was increased by more than two-fold and its γ-carboxylation was also increased when VK_1_ was included in the media (Fig. 4B). These results suggest that γ-carboxylation is activated in response to ER stress to protect β-cells from apoptosis.

**Figure 4:**
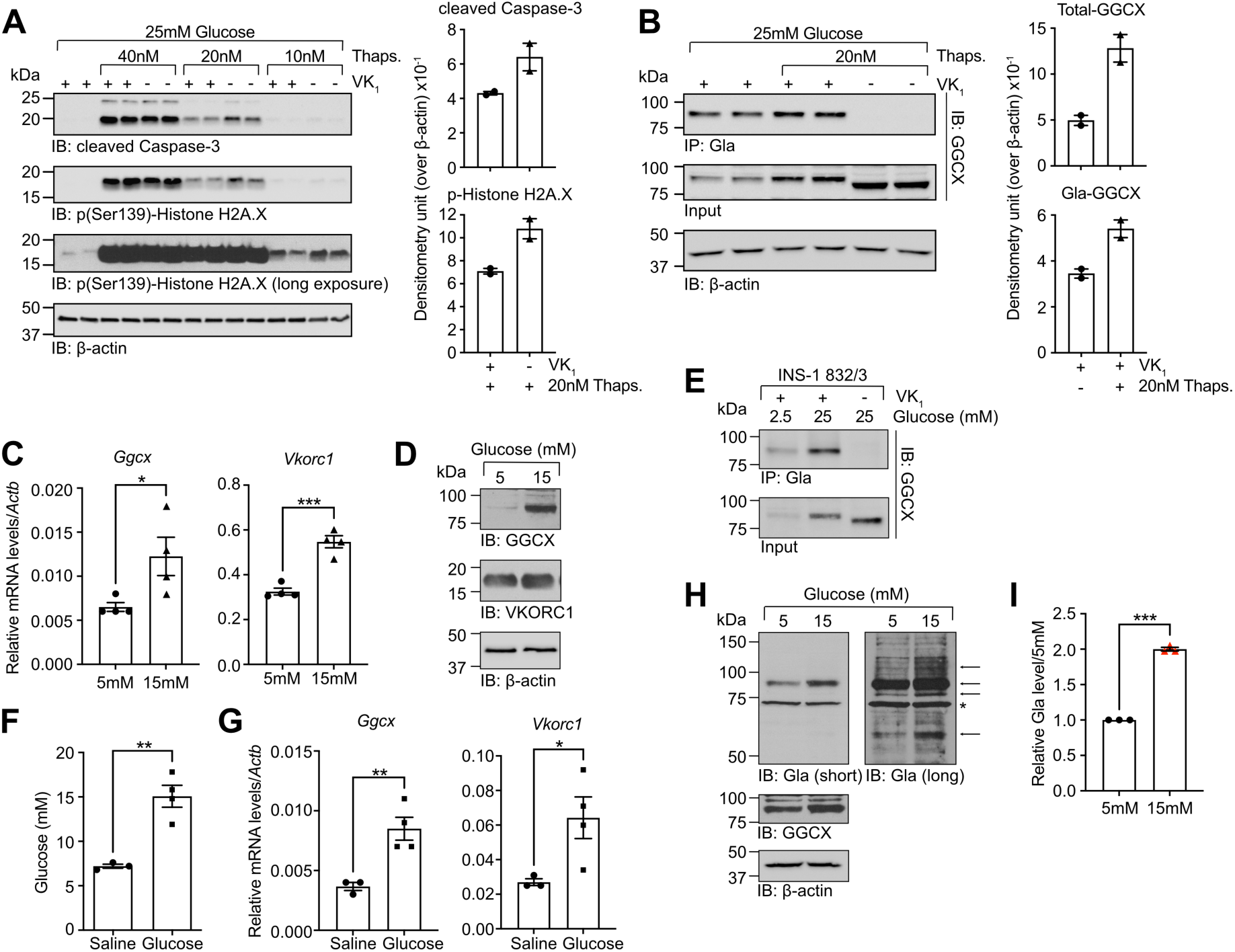
Gamma-carboxylation protects β-cells from ER stress induced cell death and is regulated by glucose. **(A-B)** INS-1 832/3 cells were cultured with VK_1_ (22μM) or vehicle for 48 hours before being cultured for 24 hours in media containing 25mM glucose and thapsigargin (0, 10, 20, 40nM). Western blot was performed to analyze **(A)** cellular fitness using cleaved-caspase-3 and p(Ser139)-Histone H2A.X antibodies. β-actin was used as a loading control. Quantification was performed using arbitrary densitometry units of cleaved-caspase-3 and p(Ser139)-Histone H2A.X signals obtained in sampled treated with 20nM thapsigargin over β-actin signals (right panels). **(B)** GGCX γ-carboxylation was assessed by anti-Gla immunoprecipitation followed by western blot using anti-GGCX antibodies. Quantification was performed using arbitrary densitometry units of GGCX and Gla-GGCX signals over β-actin signals (right panels). **(C-D)** Islets from C57BL/6J mice were cultured for 3 days in media containing either 5 or 15mM glucose. *Ggcx* and *Vkorc1* expression were analyzed by **(C)** qPCR (n=3-4) and **(D)** western blot. **(E)** INS-1 832/3 cells were cultured for 3 days in media containing 2.5 or 25mM glucose in presence of vitamin K (VK_1_; 22μM) or vehicle, and GGCX γ-carboxylation was assessed by anti-Gla immunoprecipitation followed by western blot using anti-GGCX antibodies. **(F-G)** Two-months old Wistar rats were infused during 4 days with saline or glucose and **(F)** average blood glucose for the last 3 days of infusion is shown for each mouse. **(G)** Gene expression was analyzed by qPCR (n=3-4) and data were normalized to *Actb*. **(H-I)** Human islets from non-diabetic cadaveric donors were cultured in presence of VK_1_ (22μM) in media containing either 5 or 15mM glucose for 3 days. Gamma-carboxylation and GGCX expression were analysed by western blot using anti-Gla and anti-GGCX antibodies. β-actin was used as a loading control. **(H)** Representative western blot experiment with islets from donor R288. **(I)** Gamma-carboxylation was quantified using arbitrary densitometry units of Gla signals over β-actin signals. Data from 15mM glucose treated samples were normalized over 5mM glucose treatment (n=3). Results represent the mean ± SEM; Unpaired, two-tailed Student’s *t* test was used in (C), (F), (G) and (I); \*\*\**P* < 0.001; \*\**P* < 0.01; \**P* < 0.05.

### Glucose regulates vitamin K-dependent carboxylation in β-cells

We next sought to determine if glucose itself, a physiological inducer of ER stress and UPR in β-cells (Sharma et al., 2015), could regulate the expression of the VK cycle enzymes and γ-carboxylation. We cultured wild type C57BL/6J mouse islets for 3 days in media containing either 5 or 15mM glucose and assessed the impact on GGCX and VKORC1 expression and VK cycle function. *Ggcx* and *Vkorc1* expression was increased both at the mRNA and protein level in response to 15mM glucose (Fig. 4C-D). GGCX expression and γ-carboxylation were also induced when the rat β-cell line INS-1 832/3 was cultured in the presence of high glucose (25 mM) concentrations (Fig. 4E). To confirm that glucose regulates the VK cycle in vivo, 2-month-old Wistar rats were infused with glucose for 3 days, had their islets isolated and gene expression was analyzed by qPCR. Glycemia reached ∼15 mM in glucose-infused rats (Fig. 4F), and this was sufficient to significantly increase *Ggcx* and *Vkorc1* expression in their islets (Fig. 4G) compared to rats infused with saline solution which maintained their blood glucose at ∼7 mM. Finally, GGCX expression and global γ-carboxylation were also increased in non-diabetic human islets cultured for 3 days in media containing 15mM glucose compared to islets cultured with 5mM glucose (Fig. 4H-I, Table S1). These data indicate that glucose positively regulates the level and activity of the VK cycle in rodent and human β-cells.

### ERGP is a vitamin K-dependent γ-carboxylated protein expressed in β-cells

To elucidate the molecular mechanism by which VK-dependent γ-carboxylation regulates β-cell survival and insulin secretion, we next sought to identify γ-carboxylated protein(s) present in β-cells. In our mouse islet RNAseq dataset, the genes encoding known γ-carboxylated proteins, including the clotting factors II, VII, IX and X, matrix Gla protein and osteocalcin, were all expressed at very low levels (Fig. S6A). Together with the detection in islets of multiple γ-carboxylated proteins by western blot (Fig. 1F), these observations suggest that β-cells express previously uncharacterized Gla proteins.

In the context of another project, we immunoprecipitated γ-carboxylated proteins from 5-day old wildtype (WT) mouse liver extracts using our pan-specific α-Gla antibody and identified by mass spectrometry aspartyl/asparaginyl β-hydroxylase (ASPH) as a novel putative intracellular Gla protein (Table S4). The *Asph* locus contains two promoters (P1 and P2), undergoes extensive alternative splicing and encodes for three previously characterized ER-resident proteins: aspartyl/asparaginyl β-hydroxylase (ASPH), junctate and junctin (Dinchuk et al., 2000; Feriotto et al., 2005) (Fig. S6B). ASPH is expressed in several tissues under the control of P1, while junctate and junctin expressions are controlled from P2 and appears to be restricted to cardiac and skeletal muscles (Feriotto et al., 2006). ASPH is a type II transmembrane protein of ∼80 kDa containing three luminal domains: an EF-hand calcium-binding domain, a negatively charged Glu rich domain (GRD) containing 39 Glu residues, and an alpha-ketoglutarate dependent hydroxylase domain (Fig. S6B). Junctate and junctin are ER-resident proteins corresponding to shorter variants of ASPH. Junctate and junctin share a different N-terminus cytosolic domain compared to ASPH, and junctin is characterized by a basic luminal domain instead of the acidic GRD domain found in junctate and ASPH (Fig. S6B). Islet RNAseq data indicate that overall *Asph* expression level in islets is comparable to *Ggcx* and higher than any of the other genes encoding known γ-carboxylated proteins (Fig. S6A). Consistent with previous studies showing that junctate and junctin are specifically expressed in skeletal muscle and heart from promoter P2, our own data show this promoter is inactive in mouse islets, confirming that junctate and junctin isoforms are not expressed in these cells (Fig. S6B-C). However, we detected a high level of expression for the last unique exon of junctate in islets, indicating that a truncated version of ASPH might be expressed in these cells. This shorter isoform was amplified by PCR on mouse islet cDNA using a pair of oligonucleotides located respectively in the first exon of ASPH (downstream of P1) and in the last exon of junctate. Sequencing of the PCR product identified this transcript as *Asph* variant 4. The protein encoded by this transcript is identical to the first 308 amino acids (a.a.) of ASPH, but lacks the last 433 a.a. including the hydroxylase domain (Fig. S6B). Since the N-terminus of this protein is distinct from the one of junctate and this previously uncharacterized ER-resident protein appears to be γ-carboxylated (Table S4 and see below), we decided to name it <u>E</u>ndoplasmic <u>R</u>eticulum <u>G</u>la <u>P</u>rotein (ERGP). The mRNAs encoding ASPH and ERGP are both expressed in pancreatic islets, with ERGP mRNA being at least three times more expressed than ASPH mRNA (Fig. S6C). Publicly available islet transcriptomic data further indicate that the mRNA of ASPH and ERGP, but not of junctate and junctin, are expressed in β-cells (Fig. S6B) (DiGruccio et al., 2016). Based on these observations we decided to determine if ASPH and ERGP are the γ-carboxylated proteins detected in islets and β-cells.

ASPH (A) and ERGP (E) share a Glu rich domain (GRD) that could be prone to γ-carboxylation. To detect these two proteins, we generated and affinity-purified rabbit polyclonal antibodies against the GRD domain; hereafter called α-A/E-GRD. Western blot analysis showed that these antibodies cannot detect ASPH or ERGP deletion mutants that do not possess the GRD (Fig. S6D-E). Importantly, the addition of either VK_1_ or warfarin did not change the immunoreactivity of α-A/E-GRD antibodies against full-length ASPH and ERGP, suggesting that γ-carboxylation does not impact α-A/E-GRD binding (Fig. S6E). Anti-Gla immunoprecipitation (IP) followed by α-A/E-GRD western blot or α- A/E-GRD IP followed by α-Gla western blot confirmed that both ASPH and ERGP are γ-carboxylated in one-week-old WT liver but not in the liver of *Vkorc1*^-/-^ mice lacking γ-carboxylation at this age (Fig. S6F-G) (Lacombe et al., 2018). Using the same approach, we could show that ASPH and ERGP are expressed and γ-carboxylated in adult control mouse islets, and that their γ-carboxylation was greatly reduced in islets isolated from *Vkorc1*^-/-^;*APOE-Vkorc111* mice, which have lower VK oxidoreductase activity and γ-carboxylation in all tissues except the liver (Fig. 5A and Fig. S6H) (Lacombe et al., 2018). Interestingly, these analyses showed that ERGP expression and γ-carboxylation were respectively ∼7 and 8-fold higher than ASPH in control islets (Fig. 5B). The same IP analysis performed on *Ggcx^ff^; Ins1-Cre* islets lacking γ-carboxylase activity exclusively in β-cells, revealed that ERGP, but not ASPH, is strongly γ-carboxylated in β-cells (Fig. 5C). Immunofluorescence on wildtype mouse islets confirmed the expression of ERGP in islet endocrine cells, including β-cells (Fig. 5D). Finally, α-Gla IP followed by LC-MS/MS analyses on INS-1 832/3 cells cultured in 25mM glucose and treated or not with 20nM of thapsigargin indicated that rat ERGP is also carboxylated in a VK-dependent manner in stressed β-cells (Table S5 and Fig. S6I). Together these results establish that ERGP, and to a lesser extent ASPH, are novel γ-carboxylated proteins expressed in mouse β-cells.

**Figure 5:**
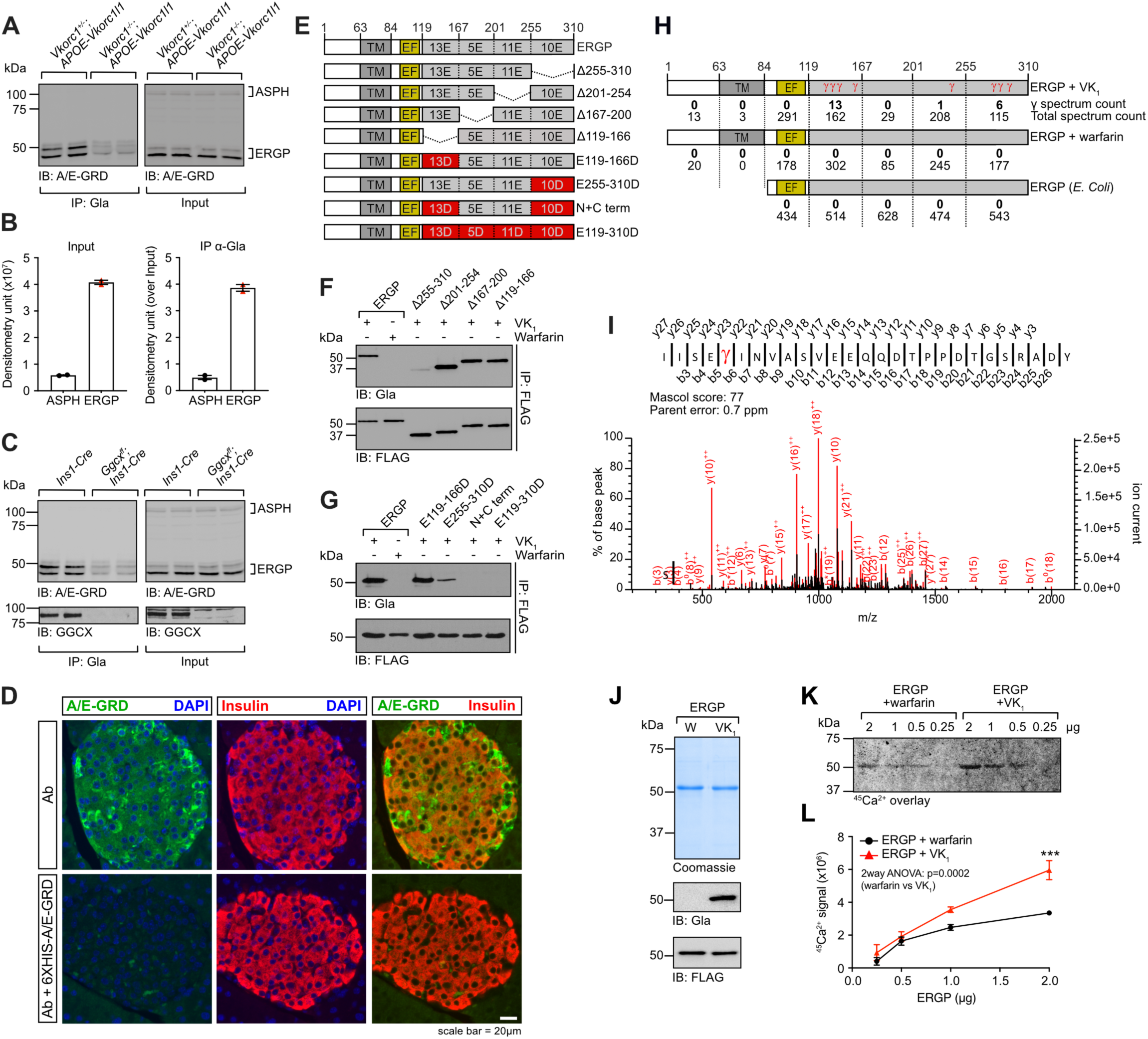
ERGP is a γ-carboxylated protein expressed in β-cells. **(A)** ASPH and ERGP γ-carboxylation was assessed in *Vkorc1^+/-^; APOE-Vkorc1l1* and *Vkorc1^-/-^; APOE-Vkorc1l1* mouse islets by immunoprecipitation with anti-Gla antibody followed by western blot with the anti-A/E-GRD antibody. **(B)** Quantification of expression and γ-carboxylation were measured using arbitrary densitometry units of ASPH and ERGP signals. **(C)** ASPH, ERGP and GGCX γ-carboxylation were assessed in *Ggcx^ff^; Ins1-Cre* and *Ins1-Cre* mouse islets by immunoprecipitation with anti-Gla antibody followed by western blot with anti-A/E-GRD or anti-GGCX antibodies. **(D)** Immunofluorescence on pancreas sections from wildtype mice using anti-A/E-GRD and anti-insulin antibodies. DAPI was used to stain nuclei. Scale bar: 20μm. The specificity of the anti-A/E-GRD antibody was demonstrated by competition of its binding with recombinant 6XHIS-A/E-GRD protein. **(E)** Schematic representation of full length ERGP, GRD deletions and glutamic acid to aspartic acid residue mutations. **(F-G)** HEK293 cells transfected with the indicated constructs were cultured with VK_1_ (22μM) or warfarin (50μM) as specified. FLAG-tagged proteins were immunoprecipitated with anti-FLAG agarose beads followed by western blot with anti-Gla or anti-FLAG antibodies. **(H)** Schematic representation of ERGP depicting the total number of spectrum counts and the total number of γ-carboxylated spectrum counts detected when ERGP is expressed in HEK293 cells in presence of VK_1_ or warfarin, or expressed in *E. coli*. **(I)** Representative LC-MS/MS spectrum showing a γ-carboxylated residue in the peptide ranging from residue 289 to 316 in purified ERGP-3XFLAG expressed in HEK293 grown in presence of VK_1_. **(J)** One μg of ERGP-3XFLAG purified from HEK293 cells cultured with VK_1_ (γ-carboxylated) or warfarin (uncarboxylated) was stained with Coomassie and γ-carboxylation monitored by western blot using anti-Gla antibodies. Anti-FLAG was used as a loading control. **(K)** Representative calcium overlay assay. Membrane-immobilized γ-carboxylated and non-carboxylated ERGP-3XFLAG were incubated with ^45^Ca^2+^ and radioactivity detected using a storage phosphorimager screen. **(L)** Calcium binding was quantified using arbitrary densitometry units (n=3). Results represent the mean ± SEM; Two-way ANOVA with Bonferroni’s post-tests was used in (L); \*\*\**P* < 0.001.

### ERGP is γ-carboxylated on several glutamic acid residues located in its Glu-rich domain

To identify the domain(s) and specific glutamic acid residues subjected to γ-carboxylation in ASPH and ERGP, we first expressed full-length ASPH-3XFLAG, or mutants lacking either the cytosolic, the EF-hand or the Glu-rich domain in HEK293 cells, a cell line which supports VK-dependent carboxylation (Lacombe et al., 2018). Full-length ASPH γ-carboxylation was detected in HEK293 cells cultured in the presence of VK_1_, but not in presence of warfarin (Fig. S6J). Although neither deletion of the cytosolic nor the EF-Hand domains significantly affected ASPH γ-carboxylation, deletion of the GRD completely abrogated its γ-carboxylation (Fig. S6J). The amino acid sequence of the GRD is poorly conserved across mammalian species. However, the enrichment of glutamic acid residues has been retained throughout evolution, suggesting a fundamental biological function for ASPH/ERGP GRD γ-carboxylation (Fig. S6K). Internal deletions within the GRD of ERGP indicated that most of the γ-carboxylation sites are in the region encompassing residues 255 to 310 and/or that the C-terminal domain contains a critical sequence for the recognition by GGCX (Fig. 5E-F). To rule out the possibility that this deletion reduces γ-carboxylation by affecting ERGP conformation and recognition by GGCX, we mutated glutamic acid residues throughout the GRD into aspartic acid residues, which cannot be γ-carboxylated by GGCX (Fig. 5E). Using this series of mutant proteins, we found that γ-carboxylated residues are mainly located in the N- and C-terminal regions of the GRD (Fig. 5G and Fig. S6L). In agreement with these findings, LC-MS/MS analysis detected the presence of γ-carboxylated residues at the N- and C-terminus of the GRD (Fig. 5H-I and Table S6). Confirming the specificity of this LC-MS/MS approach, no Gla containing peptides were identified when ERGP was purified from HEK293 cells treated with warfarin or from E. coli which lack γ-carboxylation machinery (Fig. 5H).

Since γ-carboxylation increases the affinity of proteins for Ca^2+^ (Furie et al., 1999), we next tested whether this post-translational modification could modulate the calcium-binding capacity of ERGP. Carboxylated and uncarboxylated ERGP-3XFLAG were expressed and purified from HEK293 cells cultured in the presence of VK_1_ or warfarin respectively (Fig. 5J). As revealed by ^45^Ca^2+^ overlay experiments, γ-carboxylated ERGP binds significantly more calcium than its uncarboxylated counterpart when identical amounts of protein were used in the assay (Fig. 5K-L), suggesting that the presence of Gla residues in the GRD increases ERGP capacity to bind Ca^2+^.

### Carboxylated ERGP regulates store-operated calcium entry (SOCE)

Because our data suggest that ERGP is the predominant γ-carboxylated protein present in islets and β-cells, we decided to further investigate the cellular role of this specific isoform. The cellular function of ERGP is unknown and to our knowledge has never been investigated. One study in T cells suggested that junctate acts as a calcium-sensing ER protein regulating the formation of Stromal interaction molecule 1 (STIM1) and Orai1 protein complexes (Srikanth et al., 2012). STIM1 is an ER transmembrane protein that acts as a calcium sensor which, upon ER calcium mobilization, oligomerizes and is transported to plasma membrane (PM) proximal ER where it heteromerizes with the plasma membrane calcium channel Orai1 (Lunz et al., 2019). The STIM1-Orai1 complexes are critical to activate store-operated calcium entry (SOCE), a cellular response whereby extracellular calcium enters the cytosol following ER calcium depletion to replete cellular calcium. SOCE is also a prominent calcium influx mechanism by which several cell types maintain cytosolic and ER calcium levels at rest (Palty et al., 2012; Samtleben et al., 2015). SOCE has been implicated in the regulation of insulin secretion by β-cells (Sabourin et al., 2015), with loss of STIM1 leading to reduced insulin secretion and increased ER stress (Kono et al., 2018). These observations prompted us to investigate whether ERGP and γ-carboxylation would affect cellular calcium flux and SOCE.

To eliminate potential confounding effects caused by endogenous human ASPH or ERGP, we used CRISPR/Cas-9 genome editing to knockout all *ASPH* encoded isoforms in HEK293 cells (*ASPH^-/-^* HEK293) (Fig. S7A-B). Store-operated calcium entry (SOCE) machinery was recapitulated in these cells by expressing STIM1-Myc and Orai1-HA in the presence or absence of ERGP-3XFLAG. Cells were then cultured with or without VK_1_ to modulate ERGP carboxylation (Fig. S7C). Carboxylated GGCX was detected in VK_1_ treated cells regardless of ERGP expression, confirming efficient γ-carboxylation in these cells. Ratiometric cytosolic calcium measurement by live-cell imaging with Fluo-4 and Fura Red Ca^2+^ indicators was next used to assess SOCE, which was triggered first by depleting ER calcium with thapsigargin in Ca^2+^ free buffer followed by calcium addback to the buffer (Fig. 6A). Using these experimental settings, we observed that cells expressing γ-carboxylated ERGP are characterized by a reduction in their ER calcium release as well as reduced SOCE (Fig. 6B-D). In addition, there was a significant reduction in basal cytosolic calcium level in the same cells (Fig. 6B and 6E), suggesting that ERGP, only when γ-carboxylated, restrains cytosolic calcium in cells.

**Figure 6:**
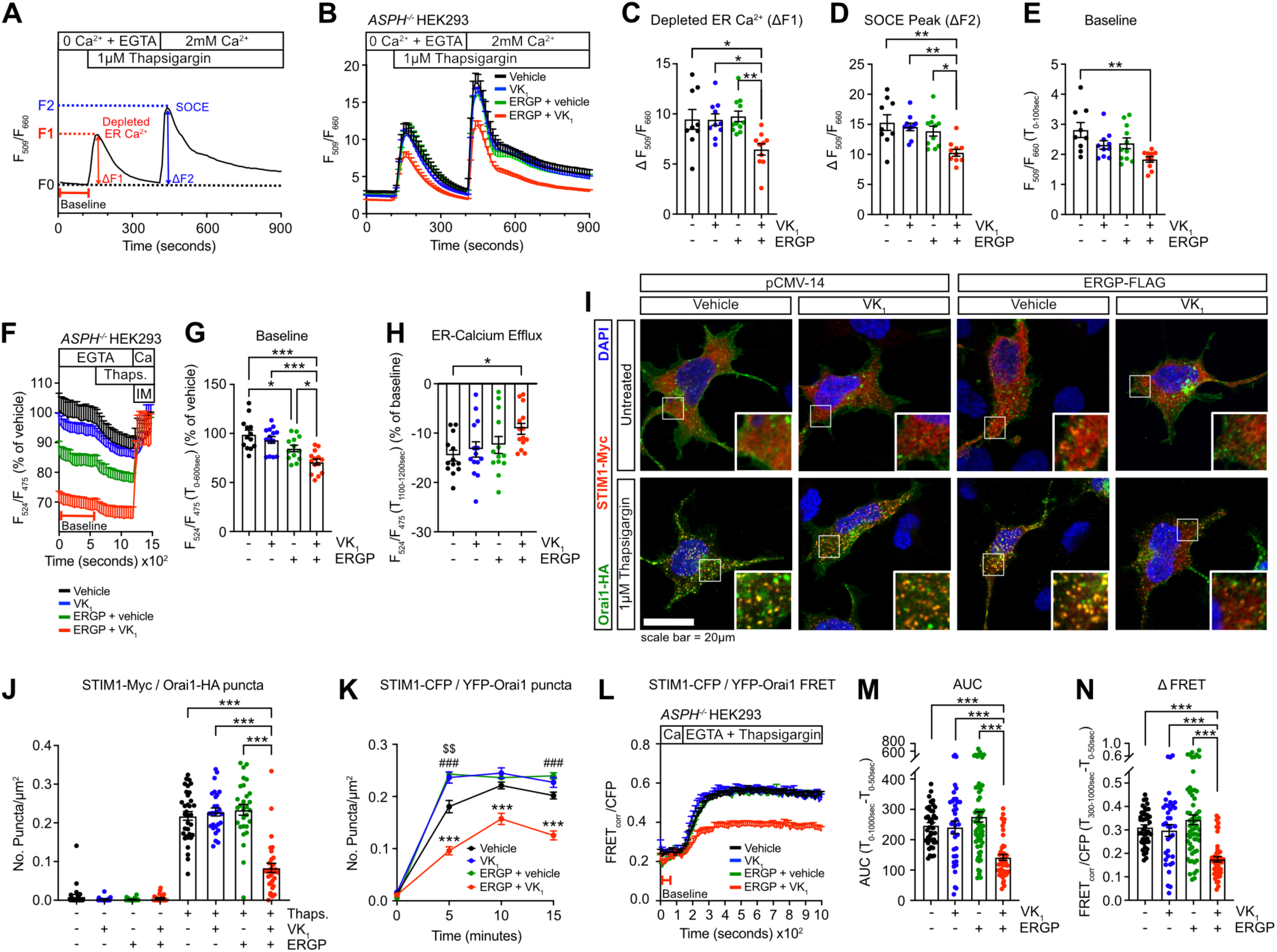
ERGP γ-carboxylation regulates calcium flux by modulating STIM1/Orai1 interaction. **(A)** Representation of the strategy used to measure and quantify store-operated calcium entry (SOCE) by live-cell imaging. Fluo-4 and Fura Red loaded cells were incubated in calcium-free buffer containing EGTA at baseline. Endoplasmic reticulum (ER)-calcium store depletion was triggered with 1μM thapsigargin and SOCE activated by the addition of calcium in the buffer at the indicated times. Calcium imaging traces are represented as the ratio between Fluo-4 and Fura Red emission intensity (F_509_/F_660_). **(B)** Calcium imaging traces represent the average F_509_/F_660_ ratio for each condition (n=9-11 experimental replicates per condition, with data from 54-130 cells averaged for each replicate). **(C-D)** Quantification of **(C)** depleted ER calcium and **(D)** SOCE peak are represented as ΔF1 and ΔF2 respectively, as described in (A). **(E**) Baseline cytosolic calcium defined as the average F_509_/F_660_ ratio during the first 100 seconds of recording. **(F-H)** Measurements of ER calcium levels in *ASPH^-/-^*HEK293 cells transfected with the D4ER ratiometric calcium probe (n=11-14 experimental replicates per condition, with data from 61-118 cells averaged for each replicate). **(F)** Representative experiment showing ER calcium level measurements by live-cell imaging in calcium-free buffer containing EGTA followed by ER-calcium store depletion with 1μM thapsigargin and maximal ER calcium replenishment with 3μM ionomycin and 2mM calcium. D4ER calcium imaging traces are expressed as a ratio between the Citrine FRET emission and CFP emission (F_524_/F_475_) for each condition. Data are normalized to the mean of the first recorded 100 seconds of the vehicle condition and are represented as percentage (%) of vehicle. **(G)** Baseline ER calcium defined as the average F_524_/F_475_ ratio during the 600 seconds of recording before thapsigargin addition. Data are represented as % of vehicle. **(H)** ER calcium efflux following thapsigargin corresponding to average F_524_/F_475_ ratio recorded during the last 100 seconds of the thapsigargin treatment expressed as a % of the average of the F_524_/F_475_ ratio recorded at baseline. **(I)** Representative confocal immunofluorescence images of *ASPH^-/-^* HEK293 transfected as in (B) and treated with 1μM thapsigargin or vehicle for 15 minutes. Cells were fixed and labeled with anti-Myc (STIM1) and anti-HA (Orai1) antibodies. DAPI was used to stain nuclei. Scale bar: 20μm. **(J)** Quantification of co-localized STIM1-myc and Orai1-HA puncta from experiment in (I). Data are represented as number of puncta/μm^2^ (n=27-34 cells per condition, from 3 independent experiments). **(K)** CFP and YFP co-expressing puncta in *ASPH^-/-^* HEK293 cells transfected with STIM1-CFP and Orai1-YFP plasmids were quantified before and following 5-, 10-, and 15-minutes treatment with 1μM thapsigargin and 3mM EGTA. Data are represented as number of puncta/μm^2^ (n=37-61 cells). **(L-N)** STIM1-CFP and YFP-Orai1 SOCE complex formation was measured by FRET. STIM1-CFP/YFP-Orai1 complex formation was triggered by the addition of 3mM EGTA and 1μM thapsigargin 120 seconds after starting recording (n=37-61 cells). **(L)** FRET signal traces averaged from individual cells represented as corrected FRET intensity over CFP intensity (FRET_corr_/CFP). **(M)** Area under the curve (AUC) of FRET_corr_/CFP traces in (L) from baseline to thapsigargin FRET signal from individual cells for each condition. **(N)** Average change in FRET_corr_/CFP from baseline (average T_0-50sec_) to peak after thapsigargin and calcium depletion (average T_300-1000sec_). Results represent the mean ± SEM; Ordinary one-way ANOVA with Bonferroni’s post-tests was used in (C-E), (G-H), (J) and (M-N); Two-way ANOVA with Bonferroni’s post-tests was used in (K); \*\*\**P* < 0.001; \*\**P* < 0.01; \**P* < 0.05. In (K), * represents comparison between and ERGP+VK_1_ and every other condition, # represents comparison between vehicle and ERGP, and $ represents comparison between vehicle and VK_1_.

The impact of ERGP and of its carboxylation on ER calcium levels was next assessed directly using the D4ER ratiometric ER-targeted Förster resonance energy transfer (FRET) based calcium sensor (Greotti et al., 2016). These analyses confirmed that in the presence of γ-carboxylated ERGP, HEK293 cells have a significantly lower basal ER free calcium level and reduced ER calcium efflux following thapsigargin treatment (Fig. 6F-H). Importantly, apparent ER calcium levels were restored to the baseline levels found in control cells by treatment with the calcium ionophore ionomycin in the presence of 2mM extracellular calcium, indicating accurate and efficient D4ER FRET measurement in all conditions (Fig. 6F and Fig. S7D). The negative impact of γ-carboxylated ERGP on ER and cytosolic calcium in steady state is consistent with the notion that SOCE is essential to maintain resting calcium levels in the ER and cytosol (Samtleben et al., 2015).

Activation of SOCE in cells is characterized by the formation of ER-PM junction puncta containing both STIM1 and Orai1, as observed following a 15 minutes thapsigargin treatment (Fig. 6I, left panels). In these conditions, VK_1_ or ERGP alone did not affect the formation of STIM1-Orai1 puncta, but the presence of γ-carboxylated ERGP significantly reduced the formation of these protein complexes (Fig. 6I-J). A FRET assay using YFP-Orai1 and STIM1-CFP was used as an orthogonal validation of the impact of γ-carboxylated ERGP on the formation of these complexes in living cells. Western blot analysis confirmed similar expression level of the STIM1-CFP and YFP-Orai1 constructs in *ASPH^-/-^* HEK293 cells in all conditions (Fig. S7E) and live-cell imaging experiments confirmed that γ-carboxylated ERGP restrains the formation of puncta co-expressing STIM1-CFP and YFP-Orai1 following 5-, 10- and 15-minutes thapsigargin treatment (Fig. 6K). Accordingly, these experiments revealed that the average corrected FRET signal following thapsigargin addition was significantly reduced in presence of γ-carboxylated ERGP (Fig. 6L-N). These results suggest that when γ-carboxylated, ERGP limits the formation of STIM1-Orai1 complexes, thus restraining SOCE.

### Increased SOCE and basal cytosolic calcium in *Ggcx*-deficient β-cells cause impaired fasting hyperinsulinemia

In islets lacking GGCX exclusively in β-cells, ERGP is not γ-carboxylated (Fig. 5C). Therefore, we next used *Ggcx^ff^; Ins1-Cre* islets as a genetic model of decarboxylated ERGP in β-cells to determine how γ-carboxylation of this protein affects calcium homeostasis in these cells. Cytosolic calcium flux was analyzed in partially dissociated islet cells from *Ggcx^ff^; Ins1-Cre* and *Ins1-Cre* (control) mice by ratiometric live-cell imaging as above. SOCE was monitored with the protocol used for the HEK293 cells (Fig. 6A) except that diazoxide, an opener of the ATP sensitive K^+^ channel, and verapamil, a voltage gated Ca^2+^ channel (VGCC) blocker, were included in the buffered solution for the duration of imaging to prevent membrane depolarization and calcium influx through VGCC (Kono et al., 2018). We first noticed that *Ggcx^ff^; Ins1-Cre* β-cells lacking γ-carboxylated ERGP were characterized by higher cytosolic calcium levels at baseline (Fig. 7A-B). Moreover, SOCE expressed as absolute value or relative to baseline (ΔF) was increased in the same cells (Fig. 7A and C). ER calcium release was measured following treatment with a higher dose of thapsigargin (10μM) and was found to be significantly higher in the *Ggcx^ff^; Ins1-Cre* β-cells (Fig. S7F). Interestingly, we also observed that the number of STIM1 containing puncta at steady state (i.e., 5mM glucose) was more elevated in these cells in comparison to control islets, suggesting that γ-carboxylated ERGP restrain SOCE to maintain proper resting ER calcium levels in β-cells (Fig. S7G-H).

**Figure 7:**
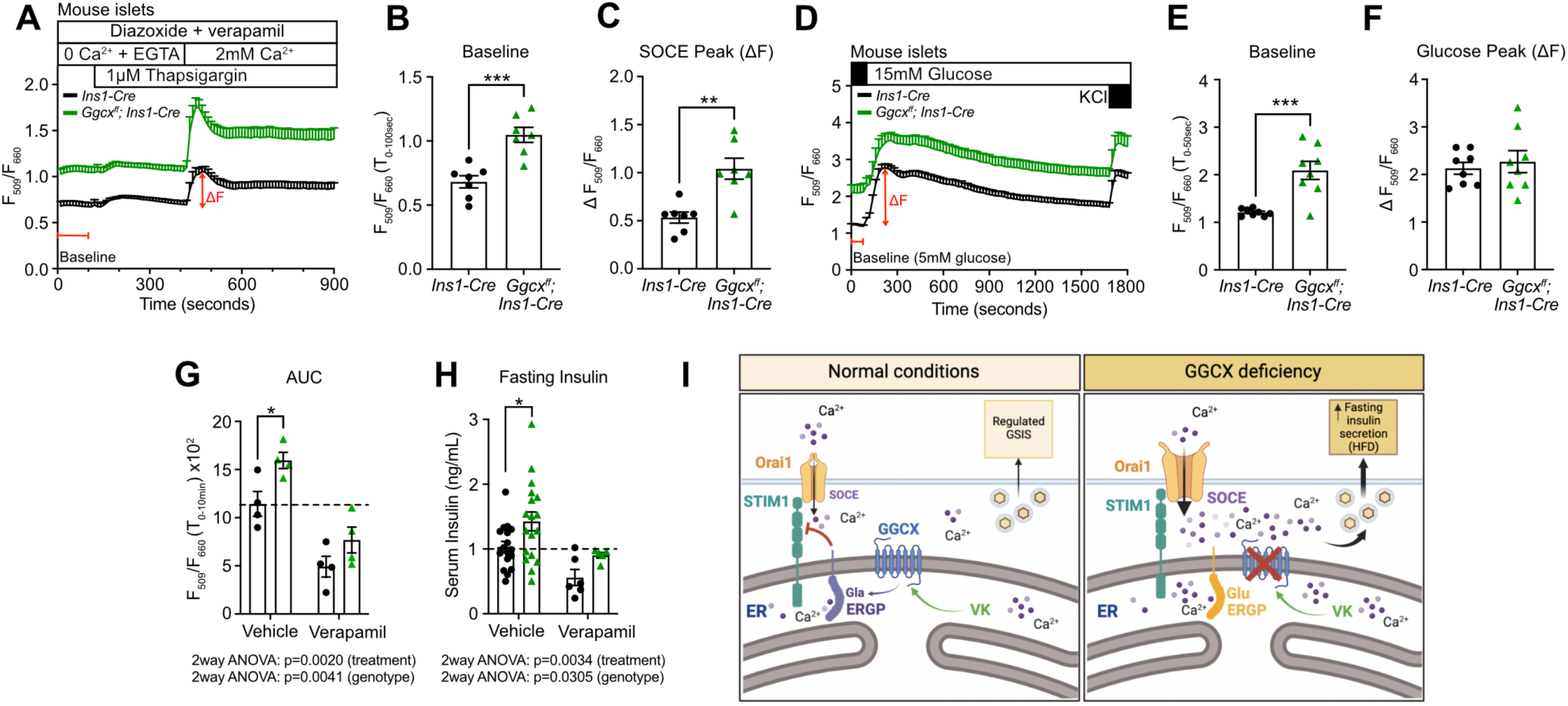
Increased SOCE and basal cytosolic calcium in *Ggcx*-deficient β-cells cause impaired fasting hyperinsulinemia. **A)** SOCE was measured by live-cell calcium imaging in semi-dispersed islets from *Ggcx^ff^; Ins1-Cre* and *Ins1-Cre* mice in buffer containing 5mM glucose, 10μM verapamil and 200μM diazoxide. Calcium traces represent the average ratio between Fluo-4 and Fura Red emission intensity (F_509_/F_660_) for each genotype (n=7 experimental replicates per genotype, with data from 82-195 individual islet cell clusters averaged for each replicate). **(B-C)** Quantification of **(B)** baseline cytosolic calcium level defined as the average F_509_/F_660_ ratio during the first 100 seconds of recording and **(C)** SOCE peak (ΔF) are represented. **(D)** Cytosolic calcium level was measured at 5 and 15mM glucose, and following addition of 30mM KCl at the indicated times. Calcium traces for each condition are represented as the ratio between Fluo-4 and Fura Red emission intensity (F_509_/F_660_) (n=8 experimental replicates per genotype, with data from 139-240 islet cell clusters averaged for each replicate). **(E)** Baseline cytosolic calcium level during the first 50 seconds of recording and **(F)** glucose-stimulated calcium peak (ΔF) quantifications are represented. **(G)** Quantification of calcium imaging experiments of semi-dispersed islets from *Ggcx^ff^; Ins1-Cre* and *Ins1-Cre* mice with Fluo-4 and Fura Red at 11mM glucose. Cells were imaged for 10 minutes, before 50μM verapamil was added for an additional 10 minutes of imaging. Data are represented as the area under the curve of the Fluo-4 and Fura Red ratio before and after verapamil addition (n=4 experimental replicates per genotypes, with data from 90-199 islet cell clusters averaged for each replicate). **(H)** *Ggcx^ff^; Ins1-Cre* and *Ins1-Cre* mice were fed a HFD for 7 days and 1mg/mL verapamil was added to their drinking water for 5 additional days. Fasting serum insulin level was measured by ELISA before and after verapamil treatment (n=5-19 mice per genotype). **(I)** Model proposed for the role of γ-carboxylation in β-cells. In normal conditions, γ-carboxylated ERGP (Gla) modulates SOCE by regulating STIM1 and Orai1 puncta formation to maintain calcium homeostasis in β-cells. In absence of GGCX, uncarboxylated ERGP (Glu) is less efficient at refraining SOCE, which increases cytosolic and ER calcium levels in β-cells. The combination of excess of nutrient (short HFD) and the rise of cytosolic calcium could lead to elevated fasting serum insulin levels. Results represent the mean ± SEM; Unpaired, 2-tailed Student’s *t* test was used in (B-C) and (E-F); Two-way ANOVA with Bonferroni’s post-tests was used in (G-H); \*\*\**P* < 0.001; \*\**P* < 0.01; \**P* < 0.05.

Calcium flux in response to glucose was next assessed. These experiments conducted in a buffered solution containing 2mM Ca^2+^ confirmed higher basal cytosolic calcium levels in *Ggcx^ff^; Ins1-Cre* islets in comparison to *Ins1-Cre* islets (Fig. 7D-E). Cytosolic calcium levels after stimulation with 15mM glucose or KCl were unchanged in the absence of carboxylated ERGP when normalized over the baseline, although they remained elevated in absolute value (Fig. 7D and 7F). Altogether, these data in primary β-cells suggest that ERGP γ-carboxylation is necessary to maintain calcium homeostasis in β-cells, mainly by suppressing SOCE. In line with the data obtained with HEK293 cells, the increased SOCE present in *Ggcx^ff^; Ins1-Cre* islet cells appears to result in increased basal cytosolic and ER calcium levels.

Chronically elevated cytosolic calcium in β-cells has been linked to dysregulated insulin secretion and fasting hyperinsulinemia (Roe et al., 1994; Yong et al., 2021). We therefore next aimed to test if pharmacologically reducing cytosolic calcium in *Ggcx^ff^; Ins1-Cre* β-cells could restore normal circulating fasting insulin levels in vivo. For this purpose, we used verapamil, a VDCC blocker and a well-characterized FDA-approved drug. Verapamil treatment of *Ggcx^ff^; Ins1-Cre* β-cells cultured with 11.1mM glucose ex vivo normalized their cytosolic calcium within a few minutes (Fig. 7G). In addition, verapamil provided in the drinking water for 5 days to mice fed a short HFD was sufficient to normalize the fasting insulin of the *Ggcx^ff^; Ins1-Cre* mice (Fig. 7H). These results indicate that increased cytosolic calcium level in β-cells is likely responsible for the fasting hyperinsulinemia observed in vivo in the *Ggcx^ff^; Ins1-Cre* mice maintained on HFD (Fig. 7I).

## DISCUSSION

### Link between the VK-cycle and stress in β-cells

VK-dependent carboxylation was previously shown to play a critical role in the liver, where it is essential for the activation of a series of coagulation factors and in arteries, cartilage, and bone where it controls the activity of two small Gla proteins: MGP and osteocalcin. Here, using conditional inactivation of *Ggcx* in mice and unique α-Gla antibodies, we establish for the first time that the VK cycle is active in pancreatic islets and, more specifically, in β-cells. Supporting a link between γ-carboxylation and β-cell adaptation to stress, we found that a large set of genes was similarly dysregulated in *Ggcx-*deficient islets and in islets isolated from pre-diabetic and diabetic mice. In addition, we found that glucose regulates γ-carboxylation activity and that VK_1_ treatment can protect β-cells from the deleterious effects of high glucose and ER-stress. Together with the observation that mice lacking *Ggcx* in pancreas or in β-cells only failed to adapt their insulin secretion in response to a short HFD feeding, our data suggest that the VK cycle could be part of a mechanism implicated in β-cell survival and function in conditions of adaptation to metabolic stress. Therefore, we could speculate that lower vitamin K intake or diminished γ-carboxylation activity could be a risk factor for the development of diabetes when combined with nutrient excess. In line with our findings, two recent large observational studies in humans indicate that treatment with warfarin compared to direct anticoagulant not targeting carboxylation does significantly increase the risk of new-onset diabetes in male and female patients with atrial fibrillation (Cheung et al., 2021; Liu et al., 2022).

### ERGP γ-carboxylation as a regulator of β-cell calcium homeostasis

Our results identified ERGP as a previously unrecognized γ-carboxylated protein present in islets and β-cells. These data indicate that γ-carboxylated ERGP reduces the formation of STIM1 and Orai1 puncta following ER calcium store depletion, thereby partially inhibiting SOCE. In addition, in the same cells, baseline cytosolic and ER calcium level were reduced. Importantly, decarboxylated ERGP did not have any effect on puncta formation or SOCE in this heterologous cell system lacking endogenous ERGP. Conversely, decarboxylation of ERGP in *Ggcx^ff^; Ins1-Cre* β-cells was associated with increased cytosolic and ER calcium levels at baseline and following SOCE. The impacts of SOCE inhibition by γ-carboxylated ERGP on ER and cytosolic calcium at steady state is consistent with the notion that SOCE is essential to maintain resting calcium levels in the ER and cytosol, as previously reported by others (Mamenko et al., 2016; Palty et al., 2012; Samtleben et al., 2015; Schulte and Blum, 2022). We cannot exclude that γ-carboxylated ERGP directly controls ER calcium levels, but we believe that it is unlikely given that decreased ER calcium level will be expected to activate SOCE (Park et al., 2009). Altogether the data imply that increased SOCE is the primary defect in *Ggcx^ff^; Ins1-Cre* β-cells.

Junctate, whose luminal domain is identical to ERGP but has a different N-terminal cytosolic sequence, has been reported to also interact with and regulates the sarco-endoplasmic reticulum Ca^2+^-ATPase 2 (SERCA2) pump and the inositol 1,4,5 trisphosphate receptors (IP3R), two other ER membrane proteins controlling calcium flux between the ER and the cytosol (Kwon and Kim, 2009; Treves et al., 2004). Whether γ-carboxylated ERGP also regulates calcium homeostasis in β-cells through a similar mechanism is currently unknown. It is important to note however, that in these earlier publications junctate was studied in its decarboxylated form, since VK was not included in the culture media. It should be also noted that since junctate and ERGP have distinct N-terminal cytosolic domains, these two proteins may affect cellular calcium homeostasis through different mechanisms. Finally, since junctate mRNA was not detected in β-cells, we did not further investigate the γ-carboxylation and function of this protein in this cell type.

STIM1 and SOCE have previously been shown to positively regulate insulin secretion and to reduce ER stress in β-cells (Kono et al., 2018; Sabourin et al., 2015). Other studies have shown that alteration in either ER or cytosolic free calcium levels can induce ER stress and β-cell death (Sabatini et al., 2019). Moreover, the expression level and activity of STIM1 and SERCA2 were found to be reduced in human or mouse T2D islets, and to correlate with altered cytosolic calcium in response to glucose (Kono et al., 2018; Liang et al., 2014). In line with these findings, it was recently reported that tunicamycin-induced ER stress decreases ER calcium levels and increases SOCE in β-cells resulting in increased basal insulin secretion (Zhang et al., 2020). Overall, this body of literature suggests that the acute activation of calcium flux between ER and cytosol is critical for several β-cell cellular processes, including survival and insulin secretion, but that chronic stimulation of calcium signaling pathway can induce ER stress, β-cell dysfunction and death (Sabatini et al., 2019). Our data show that the Ca^2+^ binding capacity of ERGP increases when γ-carboxylated and that this modification restrains SOCE suggesting that γ-carboxylation could modulate ERGP ER calcium sensing capacity. This process appears to be necessary to avoid calcium overfilling and to maintain appropriate cytosolic and ER calcium levels in β-cells. Under conditions of chronically elevated intracellular calcium levels, reducing Ca^2+^ entry would prove beneficial to prevent β-cell dysfunction and diabetes progression. In humans, insufficient VK intake may therefore contribute to β-cell dysfunction in conditions of β-cell stress by reducing ERGP γ-carboxylation, thereby increasing the risk of T2D.

### Fasting hyperinsulinemia as a driver of insulin resistance and T2D

Noteworthily, the mice lacking GGCX in the pancreas or only in β-cells and fed a HFD for 7 days are characterized not only by reduced glucose-stimulated insulin secretion and increased blood glucose, but also by increased fasting serum insulin and reduced insulin sensitivity. These observations are consistent with the notion that in conditions of nutrient excess, chronic elevation of intracellular calcium and ER stress in β-cells can lead to uncontrolled hyperinsulinemia which could ultimately result in peripheral insulin resistance (Yong et al., 2021). There is a growing number of studies in rodents and humans suggesting that prolonged fasting insulin hypersecretion precedes and promotes insulin resistance and could be the initiating event of T2D (Hudish et al., 2019; Mittendorfer et al., 2022). Conversely, reducing insulin secretion can prevent insulin resistance, obesity, and fatty liver disease (Mehran et al., 2012; Yong et al., 2021). The metabolic phenotype of the *Ggcx^ff^; Ins1-Cre* mice following a short period of HFD suggests that γ-carboxylated ERGP may be required both to prevent uncontrolled insulin secretion by β-cells in the context of nutrient excess and to preserve normal glucose-stimulated insulin secretion.

### ERGP as a novel VK-dependent protein

Since the discovery more than 45 years ago that a group of clotting factors was γ-carboxylated on specific glutamic acid residues in a VK-dependent manner (Stenflo et al., 1974), a total of only 15 unique Gla proteins have been identified in mammals. They all share a relatively well-conserved “Gla domain” characterized by the presence of 3 to 12 γ-carboxylated glutamic acid residues and two cysteines forming a disulfide bridge. Here, we identify ERGP and ASPH as two previously unrecognized Gla proteins in liver and β-cells. The presence of Gla residues in the GRD of ERGP was confirmed with two independent approaches: detection with previously reported Gla-specific antibodies and LC-MS/MS. These observations were also validated in genetic models lacking either *Ggcx* or *Vkorc1*. The GRD domain of junctate, which is shared with ERGP, was previously described as a calcium-binding domain (Treves et al., 2000) and our data indicate that the presence of Gla residues in the GRD of ERGP further increases its capacity to bind calcium in vitro.

The GRD domain encoded by the *Asph* locus possesses several unique features compared to the classical Gla proteins. First, ERGP and ASPH are ER-resident proteins, while the other known Gla proteins are either secreted or plasma membrane proteins. Second, the ASPH/ERGP GRD, with more than 190 amino acids (a.a.), is larger than the classical Gla domains which are on average less than 50 a.a. long. In addition, the sequence of the GRD is not similar in any way to the other Gla proteins, except for the presence of multiple Glu residues. Third, in contrast to the classical Gla domain, the ASPH/ERGP GRD does not contain a disulfide bridge and the Gla residues are distributed in the N- and C-terminal regions instead of being clustered in the center. Fourth, ERGP and ASPH do not contain a sequence matching the GGCX substrate recognition sequence found in the other Gla proteins. Together with the observation that GGCX itself is also γ-carboxylated and lacks such a substrate recognition sequence (Hallgren et al., 2013), it suggests that GGCX can recognize substrates through at least two different mechanisms in mammalian cells.

Several non-vertebrate metazoans, including insects and mollusks, possess in their genome genes encoding for GGCX and VKORC1 homologues (Bandyopadhyay et al., 2002). Yet, homologues for the known vertebrate Gla proteins are not present in these organisms, suggesting that the ancestral function(s) and wider biological roles of γ-carboxylation still need to be defined. Genome database searches allowed us to identify ASPH/ERGP homologues containing a GRD in several of these non-vertebrate metazoans possessing a GGCX homologue. These observations suggest that regulation of ER-cytosolic calcium homeostasis through ERGP γ-carboxylation could be an evolutionarily conserved mechanism, which antedates divergence of mollusks, arthropods, and vertebrates.

In conclusion, we identify here VK-dependent γ-carboxylation as an important post-translational modification present in β-cells which regulates the capacity of these cells to adapt to stress. We also identify two new mammalian VK-dependent proteins, ERGP and ASPH, and provide evidence that γ-carboxylation regulates β-cell calcium homeostasis through ERGP. Together, our findings extend the cellular and physiological function of VK-dependent γ-carboxylation and reveal how this pathway may interact with the development of diabetes.

## ACKNOWLEDGEMENTS

We thank C. Julien and D. Pham for mouse genotyping and cell maintenance, Dr. K. Suh for providing the B6.Cg-*Gt(ROSA)26Sor^tm14(CAG-tdTomato)Hze^*/J mice, Dr. D. Pendin and Dr. T. Pozzan for providing the D4ER probe, Dr. V. Calderon for RNA-Seq analysis and Dr. G. Karsenty for his critical reading of the manuscript. We also thank the staff of IRCM Proteomics, Microscopy, Molecular Biology and Histology Core Facilities for their technical support. Human islets for research were provided by the Alberta Diabetes Institute IsletCore at the University of Alberta in Edmonton (http://www.bcell.org/adi-isletcore.html) with the assistance of the Human Organ Procurement and Exchange (HOPE) program, Trillium Gift of Life Network (TGLN) and other Canadian organ procurement organizations. When indicated, human pancreatic islets were provided by the NIDDK-funded Integrated Islet Distribution Program (IIDP) (RRID: SCR_014387) at City of Hope, NIH Grant #2UC4DK098085. This work was supported by funding from the Canada Research Chair program (MF), Diabetes Canada (MF), Diabète Québec (MF), the Canadian Institutes of Health Research (MF, PJT-169685 and PJT-175025; VP, MOP-77686), the US National Institutes of Health (VP, R01-DK-58096) and the CMDO Network (MF). KG received scholarships from IRCM and the Natural Sciences and Engineering Research Council of Canada. JL received a fellowship from Diabetes Canada.

## Author contributions

J.L. and M.F. conceived the study, designed the experiments, and initiated the project. J.L., K.G., D.F. and M.F. collected and analyzed data. J.S., F.G., S.M.M., and S.H. collected data. A.V. and V.P. prepared cDNA from islets mRNA isolated from rats perfused with glucose or saline. M.F. and J.L. wrote the manuscript and all authors commented and contributed to editing the final version. M.F. acts as the guarantor of this work and is responsible for data access. J.L. is listed before K.G. as co–first author because J.L. conceived the study, designed the experiments, and wrote the manuscript.

## METHODS

### Experimental Model and Subject Details Mice

*Ggcx^ff^* mice were generated in our laboratory as described before (Ferron et al., 2015), and maintained on a C57BL/6J genetic background. These mice were bred to the *Pdx1-Cre* (B6.FVB-*Tg(Pdx1-cre)^6Tuv^*/Nci; National Cancer Institute; Stock 01XL5) (Hingorani et al., 2003) or the *Ins1-Cre* (B6(Cg)-*Ins1^tm1.1(cre)Thor^*/J; The Jackson Laboratory; Stock 026801) (Thorens et al., 2015) lines to generate *Ggcx^ff^; Pdx1-Cre* and *Ggcx^ff^; Ins1-Cre* mice. β-cells were labeled with the tdTomato reporter gene by breeding the B6.Cg-*Gt(ROSA)26Sor^tm14(CAG-tdTomato)Hze^*/J (Jackson Laboratory; stock 007914) mice to the *Ins1-Cre* strain. *Vkorc1^ff^* and *Vkorc1l1^ff^*mice were generated in our laboratory as described before (Ferron et al., 2015), bred to the *Pdx1-Cre* line to generate *Vkorc1^ff^; Vkorc1l1^ff^; Pdx1-Cre* mice and maintained on a C57BL/6J background. Other previously described mouse strains used in this study include *Vkorc1^-/-^; APOE-Vkorc1l1* (Lacombe et al., 2018). Male mice were used in all experiments and littermates with the appropriate genotypes always used as controls. Animals were housed at the IRCM in a pathogen-free facility on a 12h light/dark cycle and fed a normal chow diet (Teklad global 19% protein extruded rodent diet; 2919; Envigo), unless otherwise specified. All animal use complied with the guideline of the Canadian Committee for Animal Protection and was approved by IRCM institutional animal care committee.

### Human islets

Cadaveric human islets were obtained from the IsletCore at the Alberta Diabetes Institute from the University of Alberta (Edmonton, Alberta, Canada)(Lyon et al., 2016) and from the Integrated Islet Distribution Program (IIDP) at City of Hope (Duarte, California, USA). Upon arrival, islets were handpicked and processed for experiments. When needed, human islets were cultured in DMEM (5mM glucose, 10% FBS, penicillin/streptomycin) in an incubator at 37°C, 5% CO_2_. Detailed protocols for islet isolation and static glucose-stimulated insulin secretion are available in the protocols.io repository (Lyon et al., 2019) and on the IIDP website (iidp.coh.org). Donor characteristics are described in Table S1. Islet isolation was approved by the Human Research Ethics Board at the University of Alberta (Pro00013094). All donors’ families gave informed consent for the use of pancreatic tissue in research. The IRCM Human Ethics committee approved human islets use.

### Cell lines

Rat insulinoma cell line INS-1 832/3 (Millipore Sigma) was cultured in RPMI-1640 supplemented with 2mM L-Glutamine, 1mM sodium pyruvate, 10mM HEPES, 0.05 mM β-mercaptoethanol, 10% fetal bovine serum (FBS) and penicillin/streptomycin as previously described (Ronnebaum et al., 2008). HEK293 cells (ATCC) were cultured in EMEM supplemented with heat-inactivated 10% FBS and penicillin/streptomycin. Cells were cultured at 37°C with 5% CO_2_.

### Generation of *ASPH^-/-^* HEK 293 cells by CRISPR-Cas9

HEK 293 cells were transfected with single guide RNA (sgRNA; Thermo Fisher) and recombinant *Streptococcus pyogenes* Cas9 protein (SpCas9; Synthego) using Lipofectamine CRISPRMAX Cas9 Transfection reagent (CMAX00001; Thermo Fisher) according to the manufacturer protocol. We selected a sgRNA (Assay ID: CRISPR671774_SGM; Target DNA Sequence: GGACATCTGTAGCTGTCGTT) matching a sequence in the exon 2 of the gene *ASPH* which is shared by all the isoforms encoded by this gene, including ASPH, junctate, junctin and ERGP proteins. Forty-eight hours after the transfection, cells were diluted and seeded in 96-wells plates to establish clonal lines. A total of ninety-six clones were screened by standard Sanger DNA sequencing of the targeted region and the sequence analyzed using the Inference of CRISPR Edits (ICE) Analysis tools of Synthego (https://ice.synthego.com/). Two clones with frameshift-inducing indel on all alleles of *ASPH* were selected and loss of expression of ASPH and ERGP confirmed by western blot experiment and by quantitative PCR (Fig. S7A-B).

## Method Details

### Mouse islet isolation, cell sorting and culture

Mice were anesthetized by intraperitoneal (i.p.) injection of a drug mixture of ketamine hydrochloride and xylazine, sacrificed via cervical dislocation and exsanguinated. The pancreatic duct was perfused with Liberase TL (Roche Applied Science) in Hank’s balanced salt solution (HBSS) containing Ca^2+^/Mg^2+^ and the inflated pancreas was excised and incubated at 37°C for 30 minutes with firm agitation at the 15-minute mark. The digested pancreas was then washed 4 times by decantation with HBSS containing 0.1% BSA and 20mM HEPES pH 7.4, and islets were isolated using Histopaque-1077 density gradient separation (Millipore Sigma). Islets were then transferred to culture media (RPMI, 10% FBS, penicillin/streptomycin) and handpicked under a stereomicroscope (SteREO Discovery.V12; Zeiss).

Islets from *Ins1^Cre/+^*; *Rosa26^CAG-lox-stop-lox-tdTomato^*mice were dissociated by incubating at 37°C for 2 minutes using 0.05% Trypsin-ETDA and pipetting. β-cells (Tom+) and the other islet endocrine cells (Tom-) were sorted out based on tdTomato expression using the FACSAria III cell sorter (Becton Dickinson). Dead cells were excluded based on DAPI staining and only singlets were sorted. Cells were lyzed directly after sorting either in guanidium thiocyanate lysis solution for RNA extraction or in 1% triton lysis buffer for protein extraction.

### INS-1 832/3 and islet treatments

To test the presence of γ-carboxylation, INS-1 832/3 cells and human islets were cultured in their respective culture media containing either VK_1_ (22μM; V3501; MilliporeSigma), warfarin (50μM; SC-204941; Santa Cruz Biotechnology), or vehicle during 3 days before analysis. The dose of 22 μM (10 μg/ml) of VK_1_ was selected based on previous reports showing that maximal γ-carboxylation of known Gla proteins was achieved with VK_1_ concentration of >1-20 μM in cell culture (Ghosh et al., 2021; Tie et al., 2011).

### Metabolic analysis

For mice fed a regular chow diet, metabolic analysis was performed as follows. For intraperitoneal glucose tolerance tests (IPGTT), mice were fasted for 16 hours, and blood glucose levels were measured after fasting and at 15-, 30-, 60- and 120-minutes following i.p. injection of glucose (2g/kg of BW). To measure circulating insulin concentration following a bolus of glucose, tail vein blood was collected after a 16-hour fasting and at 15- and 30-minutes post-injection (i.p.) with glucose (3g/kg of BW). Serum insulin was measured using ELISA (Insulin ELISA mouse; Mercodia). Insulin tolerance test (ITT) was performed after 5 hours of fasting following i.p. injection of insulin (1U/kg; Humulin R, Lilly) and blood glucose was measured after fasting and 30-, 60-, 90- and 120-minutes post-injection.

For short high fat diet (HFD) feeding, 10-weeks old mice were fed either a lard-based diet (60% kcal from fat; TD.06414; Envigo) or an ingredient matched control diet (10% kcal from fat; TD.08806; Envigo) for 7 days, after which body weight, random fed blood glucose and metabolic tests were performed. For circulating insulin measurements, tail vein blood glucose was collected after 5 hours of fasting and at 15- and 30-minutes post-injection (i.p.) with glucose (2g/kg). Serum insulin was measured using ELISA and data represented in absolute value or as stimulation index (insulin concentration at 15 or 30 minutes / insulin concentration at fasting). For IPGTTs, tail vein blood glucose was measured after 5 hours of fasting and at 15-, 30-, 60- and 120-minutes post-injection with glucose (1.5g/kg). For oral glucose tolerance tests (OGTTs), tail vein blood glucose was measured after 5 hours of fasting and at 15-, 30-, 60- and 120-minutes post-gavage with glucose (1.5g/kg). For ITTs, tail vein blood glucose was measured after a 5-hour fasting and at 30-, 60-, 90- and 120-minutes post-injection (i.p.) with insulin (0.75U/kg). Homeostatic model assessment of insulin resistance (HOMA-IR) was calculated as follows: Fasting glucose (mM) x Fasting insulin (μU/mL)/22.5 (Lacombe et al., 2020). When indicated, following 7 days of HFD feeding, verapamil (V4629, Sigma) was added to the drinking water (1mg/mL) of the mice for 5 days while fed an HFD.

O_2_ consumption, CO_2_ release, food intake and physical activity were analyzed using an 8-chamber Promethion Continuous Metabolic System (Sable Systems International) as before (Al Rifai et al., 2017). Briefly, after a 48-hour acclimation period, data were collected for 96 hours. Energy expenditure (kcal/hour) was calculated by indirect calorimetry using the following formula: 60 x (0.003941 x VO_2_ (ml/min) + 0.001106 x VCO_2_ (ml/min)). Physical activity was measured as beam breaks for the x-, y- and z-axis using infrared beams connected to the system.

### Ex vivo glucose-stimulated insulin secretion (GSIS)

Islets from *Ggcx^ff^; Pdx1-Cre* and *Pdx1-Cre* mice were isolated and left in culture media (RPMI, 10% FBS, penicillin/streptomycin) for recovery for 40 hours. Islets were hand-picked and quickly washed in Krebs-Ringer bicarbonate HEPES (KRBH) buffer (114mM NaCl, 2.5mM CaCl_2_, 1.2mM KH_2_PO_4_, 4.7mM KCl, 1.16mM MgSO_4_, 25.5mM NaHCO_3_, 10mM HEPES and 0.1% fatty acid free BSA, pH 7.35) containing 5.6mM glucose, and then pre-incubated two times for 1 hour at 37°C. Groups of 10 similarly sized islets were transferred to microtubes containing 5.6mM glucose KRBH and incubated for 1h at 37°C. Islets were then incubated in KRBH buffer containing 11.1mM glucose, followed by KRBH buffer containing 16.7mM glucose, each incubation for 1 hour at 37°C. After each incubation, supernatant was collected and at the end of the protocol, islets were lyzed in 0.18N HCl, 75% EtOH extraction buffer. Insulin was measured in the different fractions by ELISA (Mercodia). Islets insulin content was used to normalize insulin secretion.

### Pancreas immunohistochemistry, immunofluorescence and insulin content

Pancreases were weighed and fixed in 10% formalin for 24 hours at room temperature, embedded in paraffin and sectioned at 5μm. For immunohistochemistry and immunofluorescence experiments, rehydration was followed by an antigen retrieval step (sub-boiling for 10 minutes in 10mM sodium citrate pH 6.0).

For β-cell mass quantifications, insulin was detected using rabbit anti-insulin antibodies (1:200, sc- 9168; Santa Cruz Biotechnology), Vectastain Elite ABC-peroxidase kit (Vector Laboratories; PK-6101) and NovaRED Substrate Kit (Vector Laboratories; SK-4800) following manufacturer’s instructions. Pancreas tissue was counterstained using Mayer’s hematoxylin and histomorphometric analyses were performed using the OsteoMeasure Analysis System (Osteometrics). β-cell mass was calculated as follows: β-cell area (%) x pancreas weight (mg) / 100.

For immunofluorescence, blocking was performed in PBS containing 5% normal donkey serum and 0.3% Triton for 1 hour at room temperature. Sections were then incubated with antibodies diluted in PBS, 1% BSA and 0.1% Triton, first with goat anti-insulin antibodies (sc-7839; Santa Cruz Biotechnology) and rabbit anti-A/E-GRD antibodies (generated in our laboratory, see below) over-night at 4°C, and second with Alexa-Fluor 594-conjugated donkey anti-goat (705-585-147; Jackson Immunoresearch Laboratories) and Alexa-Fluor 488-conjugated donkey anti-rabbit (711-545-152; Jackson Immunoresearch Laboratories) antibodies for 1 hour at room temperature. Specificity of anti-A/E-GRD antibodies was assessed by competition with recombinant 25μg/mL 6XHIS-A/E-GRD protein during incubation with the primary antibodies. Nuclei were stained with DAPI. Volocity 6.0 quantitation module was used to threshold for and select Insulin^+^ cells and to determine the intensity of the A/E-GRD signal in Insulin^+^ areas.

For apoptosis detection, the Click-iT Plus TUNEL Assay kit (C10617; Invitrogen) was used following manufacturer’s instructions except the proteinase K treatment was replaced by a 10-minute incubation in citrate buffer. Goat anti-insulin antibodies (sc-7839; Santa Cruz Biotechnology) and Alexa-Fluor 594-conjugated donkey anti-goat antibodies (705-585-147; Jackson Immunoresearch Laboratories) were used as described above. Nuclei were stained with DAPI. Insulin^+^ TUNEL^+^ cells were detected using the automated DM5500B fluorescence microscope (Leica) with a Retiga EXi (QImaging) and 40X objective. Volocity 6.0 quantitation module was used to threshold for and count Insulin^+^ cells with TUNEL^+^ nuclei.

For pancreatic insulin content measures, each pancreas was weighed and homogenized in an acid-ethanol buffer (1.5% HCl; 70% EtOH) after overnight fasting and 2 hours of refeeding. Samples were neutralized using 1M Tris-HCl pH 7.5 (1:1) and insulin measured by ELISA (Mercodia). Insulin content was normalized to the pancreas weight.

### RNA isolation and qPCR

For mouse and human islets gene expression analysis, 20-40 handpicked islets per mouse or donor were lyzed in guanidium thiocyanate lysis solution, and tRNA (20ug) was added before total RNA was isolated as described (Chomczynski and Sacchi, 2006). Samples were then treated with DNaseI (18068015; Invitrogen), and mRNA reversed transcribed using M-MLV reverse transcriptase (28025013; Invitrogen) and random hexamers and oligo dT primers. Relative gene expression was quantified using PowerUp SYBR Green Master Mix (A25741; Applied Biosystems) and ViiA7 Real-Time PCR System (Applied Biosystems).

### DNA constructs and transfections

Mouse ASPH-3XFLAG and ERGP-3XFLAG plasmids were generated by PCR amplification using pENTR223.1-ASPH (Clone ID: BC166658; Transomic Technologies) as a template and cloning in the HindIII and BamHI restriction sites of the p3XFLAG-CMV-14 expression vector (MilliporeSigma). ASPH-3XFLAG and ERGP-3XFLAG deletion mutants were generated by PCR using Q5 High Fidelity DNA polymerase (M0491; NEB) and primers extending in opposite directions and flanking the region to be deleted. ERGP DNA fragments containing glutamic acid to aspartic acid point mutations were synthesized (Genscript) and cloned into ERGP-3XFLAG plasmid via an internal StuI site and the 3’ BamHI site. Mouse Orai-HA was generated in two steps. First, the 3’ section of Orai1 ORF was cloned from pCMV-SPORT6-Orai1 (BC023149; Transomic Technologies) in the EcoRI and XbaI restriction sites of a pcDNA3.1-Myc-His B expression vector with an HA tag. The missing 5’ section of Orai1 ORF was cloned from mouse osteoblast cDNA in the EcoRI and Orai1 internal ApaI restriction sites. Mouse pcDNA3.1-Stim1-Myc plasmid was obtained from Addgene (17732). Using the above-mentioned constructs as template, STIM1-CFP and YFP-Orai1 were inserted into the pcDNA3.1-Myc-His B vector using Gibson assembly. pcDNA-D4ER plasmid was previously described and obtained from Dr. Diana Pendin (Greotti et al., 2016).

HEK293 cells were transfected with “tagless” pCDNA3.1-GGCX and pCDNA3.1-VKORC1 to ensure maximal γ-carboxylation, and with the indicated plasmids using Lipofectamine 2000 transfection reagent (11668019; Invitrogen) following manufacturer’s instructions. Six hours post-transfection, media was changed and VK_1_ (22μM) or warfarin (50μM) was added when specified. Generation of a clonal cell line stably expressing ERGP-3XFLAG was generated via transfection of HEK293 cells with the pERGP-3XFLAG-CMV-14 plasmid previously linearized by digestion with ScaI. Cells with integration of the plasmid were selected using G418 antibiotics and isolated colonies were expanded. ERGP expression was assessed by western blot using anti-FLAG antibodies and immunofluorescence confirming clonality.

### Calcium overlay

HEK293 cells stably expressing mouse ERGP-3XFLAG, and transfected with GGCX and VKORC1, were cultured in the presence of either VK_1_ (22μM) or warfarin (10μM). Purified γ-carboxylated and uncarboxylated ERGP-3XFLAG proteins were resolved by SDS-PAGE, transferred to a nylon membrane and cross-linked with 0.5% glutaraldehyde. The membrane was quenched with 50mM glycine and washed 3 times with binding buffer containing 60mM KCl, 5mM MgCl_2_, 10mM imidazole-HCl, pH 6.8, and then incubated with radiolabelled binding buffer containing 8.8μM ^45^CaCl_2_ (PerkinElmer) for 1 hour, washed with distilled H_2_O and dried. Radioactivity was captured by a storage phosphor screen and detected by a laser scanner imaging system (Typhoon FLA 9500; Cytiva).

### Generation of rabbit polyclonal anti-ASPH/ERGP-GRD (anti-A/E-GRD) antibodies

Rabbit ERGP ER luminal domain anti-serum was generated by immunizing rabbits with a 6XHIS tagged protein containing amino acid 85-310 of ERGP (MediMabs). Antibodies specific to the glutamic acid rich domain (GRD) were affinity purified using a GST-tagged protein corresponding to the GRD. The specificity of the antibody towards the GRD of ERGP and ASPH was tested by western blot.

### Immunoprecipitation and western blot

Cells or tissues were homogenized in lysis buffer containing 20mM Tris-HCl (pH 7.5), 150mM NaCl, 1mM EDTA (pH 8.0), 1mM EGTA, 2.5mM NaPyrophosphate, 1mM β-glycerophosphate, 10mM NaF, 1% Triton, 1mM phenylmethylsulfonyl fluoride (PMSF) and protease inhibitors (4693132001; Roche Diagnostics). For anti-Gla immunoprecipitation, 200μg of protein extracts was incubated with 10μg of rabbit anti-Gla antibodies overnight with rotation at 4°C followed by 3 hours incubation with Protein A-Agarose beads (11719408001; Roche Diagnostics) and washed 4 times with lysis buffer.

Immunoprecipitated proteins were heated at 70°C in Laemmli buffer for 10 minutes before resolving on a 7.5% polyacrylamide Tris-Glycine gels. Proteins were detected using standard western blot procedures with rabbit anti-GGCX (16209-1-AP; ProteinTech) or rabbit anti-A/H antibodies generated in our laboratory (see above). For anti-A/E-GRD IP, the same procedure was followed. FLAG-tagged proteins were immunoprecipitated from 100μg of protein extracts using anti-FLAG agarose beads (A2220; MilliporeSigma) incubated for 2 hours with rotation at 4°C. Densitometry analyses were performed with the Image Lab software (version 5.0; Bio-Rad Laboratories).

Other antibodies used for western blot in this study include rabbit anti-VKORC1 generated in our laboratory and previously reported (Ferron et al., 2015), rabbit anti-cleaved Caspase-3 (9661; Cell Signaling), rabbit anti-phospho(Ser139)-Histone H2A.X (9718; Cell Signaling), rabbit anti-phospho(S724)-IRE1 (ab124945, Abcam), rabbit anti-GRP78/BIP (11587-1-AP, Proteintech), rabbit anti-GFP (50430-2-AP, Proteintech), mouse anti-β-Actin (A1978; MilliporeSigma), rabbit anti-GAPDH (5174; Cell Signaling), mouse anti-Myc (2276; Cell Signaling), rabbit anti-HA (C29F4; Cell Signaling), mouse anti-FLAG (F1804; MilliporeSigma) and rabbit anti-FLAG (14793; Cell Signaling). To detect VKORC1, cleaved Caspase-3 and p(Ser139)-Histone H2A.X, proteins were resolved on 10% polyacrylamide Tris-Tricine gels.

### Calcium live-cell imaging in *ASPH^-/-^* HEK293 and islets

*ASPH^-/-^* HEK 293 cells were transfected with 0.5μg GGCX, 0.5μg VKORC1, 0.44μg STIM1-Myc, 0.22μg Orai1-HA, 0.33μg ERGP-3XFLAG or p3XFLAG-CMV-14 empty vector as indicated in 35mm culture dishes and the next day cells were plated on poly-L-lysine (P1274; MilliporeSigma) coated glass coverslips (18mm diameter, #1.5 thickness; 72290-08; Electron Microscopy Sciences). Coverslips were coated with 0.1mg/mL poly-L lysine in sterile ddH_2_O at room temperature for 1 hour, before being washed three times with sterile ddH_2_O and left to dry for 2 hours. *ASPH^-/-^* HEK 293 cells were loaded with 5μM Fluo-4 AM (F14201; Invitrogen) and 2.5μM Fura Red AM (F3021; Invitrogen) in a HEPES-buffered saline solution (HBSS; 120mM NaCl, 5mM KCl, 0.8mM MgSO_4_, 2mM CaCl_2_, 10mM Glucose, 20mM HEPES, pH 7.4) containing 0.02% pluronic F-127 (P6866; Invitrogen) at 37°C for 30 minutes. Cells were then washed twice with HBSS and incubated for an additional 30 minutes at 37°C to allow complete de-esterification of intracellular AM esters. Calcium imaging was performed in Ca^2+^ free HBSS containing 1mM EGTA. Baseline measurements (F0) were recorded for 120 seconds before ER calcium stores were depleted by adding thapsigargin (final 1μM) and SOCE was triggered by adding CaCl_2_ (final [Ca^2+^] 2mM) 420 seconds after starting recording. For measurements using the D4ER calcium probe, *ASPH^-/-^* HEK293 cells were transfected with 0.2μg D4ER, 0.4μg GGCX, 0.4μg VKORC1, 0.44μg STIM1-Myc and 0.22μg Orai1-HA and 0.33μg ERGP-3XFLAG or p3XFLAG-CMV-14 empty vector. The cells were equilibrated in HBSS for 45 minutes at 37°C before being imaged in Ca^2+^ free HBSS containing 1mM EGTA. Baseline measurements were recorded for 600 seconds before ER calcium stores were depleted by adding thapsigargin (final 1μM). The calcium ionophore ionomycin (final 3μM) and CaCl_2_ (final [Ca^2+^] 2mM) were added 1200 seconds after starting recording. Imaging of HEK293 cells was performed at 37°C.

Isolated mouse islets were semi-dispersed by digestion with 0.025% Trypsin-ETDA for 1 minute followed by up and down pipetting then transferred to culture media (RPMI, 10% FBS, penicillin/streptomycin) containing 22μM VK_1_. Semi-dispersed islets enclosed in 200 μL droplets (corresponding to approximately 100 islets) were plated on glass coverslips and allowed to attach for 30 minutes at 37°C, before adding 1mL of culture media for over-night recovery. Islet cells were then loaded with 5μM Fluo-4 AM and 2.5μM Fura Red AM in KRBH buffer (114mM NaCl, 1.2mM KH_2_PO_4_, 4.7mM KCl, 1.16mM MgSO_4_, 25.5mM NaHCO_3_, 2.5mM CaCl_2_, 5mM Glucose, 20mM HEPES, 0.2% fatty acid free bovine serum albumin [BSA]) for 30 minutes at room temperature. Baseline fluorescence ratio (F0) was measured for 90 seconds and response to 15mM glucose was recorded during 90-1700 seconds before KCl concentration was raised to 30mM. SOCE in islet cells was measured as described above for HEK293 but in KRBH buffer containing 200μM diazoxide (D9035; MilliporeSigma), an opener of the β-cell ATP sensitive K^+^ channel in beta cells, and 10μM verapamil (V4629; Millipore Sigma), a voltage gated Ca2+ channel blocker, for the duration of imaging. In other experiments, the effect of verapamil on cytosolic calcium levels of semi-dispersed islets was measured as described above, but in KRBH buffer containing 11mM glucose to mimic in vivo conditions. The semi-dispersed islets were first imaged for 600 seconds following vehicle (DMSO) treatment, then were imaged for an additional 600 seconds following treatment with 50μM verapamil. Imaging for islets was performed at 32°C and 5% CO_2_ enrichment.

Imaging was performed on a confocal rotary disk inverted microscope from Zeiss equipped with a Yokogawa CSU-1 module using a 20X objective. The microscope stage contained a conduction heater and was enclosed in an incubator to maintain cells at the desired temperature and CO_2_ percentage. Fluo-4 was excited with a 488nm laser and emission was recorded at 509nm (ZEN blue software). Laser power was set to 5%, exposure to 250ms, and EM gain to 500. One image was taken every 5 seconds. Fura Red was excited with a 488nm laser and emission was recorded at 660nm. Laser power was set to 20%, exposure to 500ms, and EM gain to 750. One image was taken every 5 seconds. D4ER reports ER calcium level via FRET between its CFP and Citrine domains, which come into closer proximity following calcium binding of its D4 and calreticulin domains. To record the CFP donor signal, D4ER was excited with a 405nm laser then emission recorded at 475nm. To record the FRET signal from Citrine, D4ER was excited with a 405nm laser then emission recorded at 524nm. Laser power was set to 20%, exposure to 500ms, and EM gain to 750 to record CFP. Laser power was set to 20%, exposure to 500ms, and EM gain to 500 to record Citrine FRET.

Quantification was performed using Fiji (Schindelin et al., 2012). Timelapses were 16-bit greyscale image stacks saved as Carl Zeiss Image data format files. Individual cells or islet clusters were selected as freehand selections to generate regions of interest (ROI). The mean gray value of each ROI was then measured for each image (1 per 5 seconds) for each channel (Fluo-4/Fura Red or Citrine/CFP). Cells that did not stay attached during the entire protocol were excluded from analysis. Fluo-4 fluorescence intensity was divided by Fura Red intensity to generate a ratiometric measurement of cytosolic calcium as previously described (Ferron et al., 2011). Similarly, the intensity of the FRET signal from Citrine was divided by the CFP intensity to generate a ratiometric measurement of ER calcium for the experiments using the D4ER probe as previously described (Greotti et al., 2016). The FRET-Citrine/CFP ratio was then normalized to the mean of the first 100 seconds recorded for the vehicle condition (i.e., without VK_1_ and without ERGP) and represented as % of vehicle. A total of n=54-130 *ASPH^-/-^* HEK 293 cells were averaged together for each experiment for cytosolic or ER calcium measurements, and n=9-14 independent experiment per condition were analyzed. In semi-dispersed islet imaging experiments, 82-240 islet cell clusters were averaged for each experiment, and n=7-8 independent experiment per condition were analyzed.

### STIM1/Orai1 puncta quantification by immunofluorescence

*ASPH^-/-^* HEK293 cells were transfected with GGCX, VKORC1, STIM1-myc, and Orai1-HA and ERGP-3XFLAG or p3XFLAG-CMV-14 empty vector, and plated on poly-L-lysine coated glass coverslip as detailed in the calcium live-imaging section. Two days later, cells were equilibrated in HBSS for 45 minutes and treated with 1μM thapsigargin for 15 minutes in Ca^2+^ free HBSS containing 1mM EGTA. Cells were then fixed in 4% paraformaldehyde in PBS for 15 minutes at room temperature and washed 3 times with PBS. Cells were stained using mouse anti-Myc (Cell Signalling; 2276) and rabbit anti-HA (Cell Signalling; 3724) antibodies as detailed in the immunofluorescence section and imaged with a 63x objective.

Quantification of puncta was performed using the Volocity 6.0 quantitation module. Puncta were determined by adjusting an intensity threshold and including only objects measuring between 0.1-and 1.0μm^2^. Saturated pixels were excluded from quantification. STIM1/Orai1 co-expressing puncta in *ASPH^-/-^* HEK293 cells were defined as areas of overlapping STIM1 and Orai1 expression in objects between 0.1-1.0μm^2^ in surface area. On average a cell contains 0-120 puncta in the absence of thapsigargin and 15-413 puncta following thapsigargin treatment. The number of STIM1/Orai1 colocalized puncta per cell was normalized to the cell area in μm^2^ (n=27-34 cells per condition, from 3 independent experiments). Islet clusters stained for STIM1 contained on average 0-57 puncta at steady state (KRBH buffer with 5mM glucose). The number of STIM1 puncta per islet cluster was normalized to the cluster area in μm^2^ (n=116-183 islets clusters per condition, from 2 independent experiments).

### STIM1/Orai1 FRET measurements

*ASPH^-/-^* HEK293 cells were transfected with 0.4μg GGCX, 0.4μg VKORC1, 0.44μg STIM1-CFP, 0.42μg YFP-Orai1 and 0.33μg ERGP-3XFLAG or p3XFLAG-CMV-14 empty vector and plated on poly-L-lysine coated glass coverslips the next day as detailed in the calcium live-cell imaging section. Two days later, cells were washed twice with HBSS and imaged for 120 seconds before 1μM thapsigargin and 3mM EGTA was added to deplete ER calcium stores and chelate extracellular calcium. Transfection were optimized to get <2-fold difference between STIM1-CFP and YFP-Orai1 expression levels. Images were recorded every three seconds with the 63X objective of a confocal rotary disk inverted microscope as described in the calcium live-cell imaging section.

STIM1-CFP was excited with a 405nm laser and emission was recorded at 475nm. Laser power was set to 20%, exposure to 750ms, and EM gain to 750. YFP-Orai1 was excited with a 488nm laser and emission was recorded at 524nm. Laser power was set to 10%, exposure to 250ms, and EM gain to 500. FRET signal was excited with the 405nm laser and emission was recorded at 524nm. Laser power was set to 20%, exposure to 500ms, and EM gain to 500.

Quantification was performed using Fiji (Schindelin et al., 2012) as previously described (Picard et al., 2009; Xia and Liu, 2001). Time lapse images were 16-bit greyscale image stacks saved as Carl Zeiss Image data format files. Each timepoint contained three images representing the CFP, YFP, and FRET channels. Individual cells were selected manually using the freehand ROI selection tool. Identical ROIs were used across timepoints and channels for each time lapse. An ROI containing no cells was also selected on each time lapse and used to subtract the background value of each channel. For each imaging experiment, cells transfected with STIM1-CFP or YFP-Orai1 alone were used to calculate bleed-through of the CFP or YFP signals into the FRET channel. The bleed-through coefficient for CFP (A) and YFP (B) were individually calculated, using linear regression of CFP or YFP intensity in their respective channels compared to the signal in the FRET channel.

Corrected FRET signal was defined as follows:

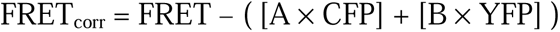

Where FRET is the fluorescent intensity at 524nm from the FRET channel, CFP is the fluorescent intensity at 475nm from the CFP channel, YFP is the fluorescent intensity at 524nm from the YFP channel, A is the CFP-FRET bleed-through coefficient determined by imaging CFP only cells, and B is the YFP-FRET bleed-through coefficient determined by imaging YFP only cells. Corrected FRET was divided by CFP intensity to control for transfection efficiency (FRET_corr_/CFP).

Area under the curve (AUC) of FRET_corr_/CFP traces were calculated with baseline AUC subtraction.

### RNA-sequencing

Total RNA from islets was extracted using the RNeasy Plus Mini Kit (74134; Qiagen) following manufacturer’s instructions (n=4 for each genotype). RNA integrity was evaluated using the 2100 Bioanalyzer system (Agilent) and samples with RIN >7.5 were used. From 1μg of total RNA, poly(A)+ transcripts were enriched using the NEBNext Poly(A) Magnetic isolation module (E7490; NEB) and libraries prepared using the KAPA stranded RNA-seq library preparation kit (KR0934; Roche Diagnostics) and the TruSeq DNA library Prep LT kit (Illumina) according to the manufacturer’s procedures. Clustering of the equimolar libraries in the flowcell was performed using the HiSeq PE cluster kit v4 cBot (PE-401-4001; Illumina) and the cBot cluster generation system (Illumina).

Sequencing was performed at the Génome Québec Innovation Center using the Illumina HiSeq 2500 system (average of 56 million paired end reads (PE50) per sample). The quality of the raw reads was assessed with FASTQC v0.11.8. After examining the quality of the raw reads, no trimming was deemed necessary. The reads were aligned to the GRCm38 genome with STAR v2.7.6a with more than 87% of reads uniquely mapped. Raw counts were computed using FeatureCounts v1.6.0 based on Ensembl mouse reference genome v101. Differential expression was performed using the DESeq2 R package and 319 differentially expressed genes (DEGs) were obtained using p-adjusted ≤ 0.05. Gene set enrichment and gene network analyses were performed using StringDB (https://string-db.org/) interrogating Biological Process (Gene Ontology), KEGG Pathways and Annotated Keywords (UniProt). Enrichment was considered significant if the false discovery rate was <0.05 and only networks of 800 or fewer genes were considered for the analysis. Previously published transcriptomic analysis of islets from pre-diabetic (5-weeks old C57BL/6 mice fed a HFD for 8 weeks) or diabetic (8-weeks old *Lepr^db/db^* and 7-weeks old *Ire1α^ff^; Ins2-Cre^ERT/+^*) mouse models (Lee et al., 2020; Motterle et al., 2017; Wang et al., 2012), were downloaded directly from the publications or from GEO (https://www.ncbi.nlm.nih.gov/geo/). Up-regulated and down-regulated genes for each model were selected using different p-adjusted values to limit the variability in the total number of genes included in each list: p<0.05 for *Lepr^db/db^*, p<0.01 for HFD and p<0.001 for *Ire1α^ff^; Ins2-Cre^ERT/+^*. Overlap between the various transcriptomes was next determined using jvenn (http://jvenn.toulouse.inra.fr/app/example.html). The statistical significance between each pair of comparisons was computed using an online tool (http://nemates.org/MA/progs/overlap_stats.html).

### Identification of carboxylated proteins by LC-MS/MS

Livers from 5-day old WT and *Vkorc1^-/-^* mice, or INS-1 832/3 cells cultured with VK_1_, 25mM glucose with or without 20nM thapsigargin, were homogenized in lysis buffer and carboxylated proteins immunoprecipitated as described above followed by three washes with 50mM ammonium bicarbonate. Immunoprecipitated proteins were then digested on-bead with trypsin at 37°C for 18 hours using 0.25ug of Sequencing grade trypsin (Promega). The samples were then reduced with 9 mM dithiothreitol at 37°C for 30 minutes and, after cooling for 10 minutes, alkylated with 17 mM iodoacetamide at room temperature for 20 minutes in the dark. The supernatants were acidified with trifluoroacetic acid and cleaned from residual detergents and reagents with MCX cartridges (Waters Oasis MCX 96-well Elution Plate) following the manufacturer’s instructions. After elution in 10% ammonium hydroxide /90% methanol (v/v), samples were dried with a Speed-vac, reconstituted under agitation for 15 min in 11 µL of 2%ACN-1%FA and 2.4% of each sample was loaded into a 75 μm i.d. × 150 mm Self-Pack C18 column installed in the Easy-nLC II system (Proxeon Biosystems). The buffers used for chromatography were 0.2% formic acid (buffer A) and 90% acetonitrile/0.2% formic acid (buffer B). Peptides were eluted with a two slopes gradient at a flowrate of 250 nL/min. Solvent B first increased from 2 to 44% in 100 min and then from 44 to 88% B in 20 min. The HPLC system was coupled to Orbitrap Fusion mass spectrometer (Thermo Scientific) through a Nanospray Flex Ion Source. Nanospray and S-lens voltages were set to 1.3-1.7 kV and 50 V, respectively. Capillary temperature was set to 225 °C. Full scan MS survey spectra (m/z 360-1560) in profile mode were acquired in the Orbitrap with a resolution of 120,000 with a target value at 3e5. The 25 most intense peptide ions were fragmented in the HCD cell and analyzed in the linear ion trap with a target value at 2e4 and a collision energy at 29. Target ions selected for fragmentation were dynamically excluded for 30 sec after 2 MS/MS events.

The peak list files were generated with Proteome Discoverer (version 2.3) using the following parameters: minimum mass set to 500 Da, maximum mass set to 6000 Da, no grouping of MS/MS spectra, precursor charge set to auto, and minimum number of fragment ions set to 5. Protein database searching was performed with Mascot 2.6 (Matrix Science) against the UniProt Mus Musculus protein database. The mass tolerances for precursor and fragment ions were set to 10 ppm and 0.6 Da, respectively. Trypsin was used as the enzyme allowing for up to 1 missed cleavage. Cysteine carbamidomethylation was specified as a fixed modification, and methionine oxidation, glutamic acid carboxylation and phosphorylation S/T/Y as variable modifications. Data interpretation was performed using Scaffold (version 4.8) using a peptide threshold of 80%, a protein threshold of 95% and one peptide minimum. We considered a protein as being carboxylated when the average exclusive spectrum count in WT samples was at least double of the *Vkorc1^-/-^*samples. To minimize the potential identification of proteins non-specifically binding the anti-Gla antibodies or the agarose-beads, we excluded proteins with more than 2 exclusive spectrum counts in the *Vkorc1^-/-^*samples or with a difference of less than 2 between the WT and *Vkorc1^-/-^*samples.

### Identification of carboxylated residues in ERGP by LC-MS/MS

HEK293 cells stably expressing mouse ERGP-3XFLAG were cultured in the presence of either VK_1_ (22μM) or warfarin (10μM) for at least 2 weeks, and carboxylated and uncarboxylated ERGP-3XFLAG were purified with anti-FLAG agarose beads (A2220; MilliporeSigma). On-bead proteins were first diluted in 2M Urea/50mM ammonium bicarbonate, and on-bead chymotrypsin digestion was performed overnight at 37°C. The supernatants were acidified with trifluoroacetic acid and cleaned from residual detergents and reagents with MCX cartridges (Waters Oasis MCX 96-well Elution Plate) following the manufacturer’s instructions. After elution in 10% ammonium hydroxide /90% methanol (v/v), samples were dried with a Speed-vac, reconstituted under agitation for 15 min in 11 µL of 2%ACN-1%FA and loaded into a 75 μm i.d. × 150 mm, Self-Pack C18 column, installed in the Easy-nLC II system (Proxeon Biosystems). Peptides were loaded on-column and eluted with a two-slope gradient at a flow rate of 250 nL/min. Solvent B first increased from 1 to 32% in 86 min and then from 32 to 82% B in 22 min. The HPLC system was coupled to Orbitrap Fusion mass spectrometer (Thermo Scientific) through a Nanospray Flex Ion Source. Nanospray and S-lens voltages were set to 1.3-1.8 kV and 50 V, respectively. Capillary temperature was set to 250 °C. Full scan MS survey spectra (m/z 320-1520) in profile mode were acquired in the Orbitrap with a resolution of 120,000 with a target value at 5e5. The most intense peptide ions were fragmented by ETD, CID and ETciD and analysed in the linear ion trap with a target value at 1e4. The peptide ion fragmentation parameters were as follow: a reaction time of 120 ms, a reagent target of 2.0e5 and a maximum reagent injection time of 200 ms for ETD, a normalized collision energy of 32% for CID, calibrated charge dependent ETD parameters and normalized supplemental activation at 18% for ETciD. The duty cycle was set to 4 seconds and target ions selected for fragmentation were dynamically excluded for 30 sec after 2 MS/MS scan events. Uncarboxylated bacterially produced His-tagged ERGP was digested in-solution with chymotrypsin in the aforementioned conditions.

The peak list files were generated with Proteome Discoverer (version 2.1 or 2.4) using the following parameters: minimum mass set to 500 Da, maximum mass set to 6000 Da, no grouping of MS/MS spectra, precursor charge set to auto, and the minimum number of fragment ions set to 5. Protein database searching was performed with Mascot 2.6 (Matrix Science) against a user-defined mouse ERGP database. The mass tolerances for precursor and fragment ions were set to 10 ppm and 0.6 Da, respectively. A semi-specific search was performed using chymotrypsin as the enzyme allowing for up to 1 missed cleavage. Methionine oxidation and carboxylation of glutamic acid were specified as variable modifications. Data interpretation was performed using Scaffold (version 4.8).

### Statistics

Statistical analyses were performed using GraphPad Prism software (version 9.3.1). Results are given as means ± SEM. For single measurement, unpaired, 2-tailed Student’s *t* test was used. Grouped analysis was performed using one-way ANOVA, followed by Bonferroni’s multiple comparisons test. For repeated measurements (metabolic tests), two-way ANOVA followed by Bonferroni’s post tests were used. Linear correlations were analyzed using Pearson’s correlation. In all figures, \**P* < 0.05; \*\**P* < 0.01; \*\*\**P* < 0.001. All experiments were repeated at least 3 times or performed on at least 3 independent animals.

## DATA AND CODE AVAILABILITY

RNA-seq data have been deposited at GEO and are publicly available as of the date of publication. Accession number is GSE199319.

The mass spectrometry proteomics dataset “Identification of vitamin K-dependent proteins in mouse liver by LC-MS/MS” has been deposited to the ProteomeXchange Consortium via the PRIDE partner repository (Perez-Riverol et al., 2022) with the dataset identifier PXD032920 and 10.6019/PXD032920.

The mass spectrometry proteomics dataset “Identification of gamma-carboxyglutamic acid residues in mouse ERGP by LC-MS/MS” have been deposited to the ProteomeXchange Consortium via the PRIDE partner repository (Perez-Riverol et al., 2022) with the dataset identifier PXD032955 and 10.6019/PXD032955.

## SUPPLEMENTAL FIGURE LEGENDS

**Figure S1, related to Figure 1:** *Ggcx* and *Vkorc1* are expressed in pancreatic endocrine cells. **(A-B)** *Ggcx* and *Vkorc1* gene expression was analyzed by quantitative PCR in various tissues from wild-type (WT) mice and normalized to *Actb*. **(C-D)** Violin plots representing single cell transcriptome data from mouse pancreatic tissues (https://tabula-muris.ds.czbiohub.org/). **(E-F)** *Ggcx* gene expression was analyzed by quantitative PCR in islets from **(E)** *Ggcx^ff^; Pdx1-Cre*, **(F)** *Ggcx^ff^; Ins1-Cre* and their respective *Ggcx^ff^* controls and normalized to *Actb* or *Gapdh* (n=4-5). Results represent the mean ± SEM; Unpaired, 2-tailed Student’s *t* test was used in (E-F). \*\*\**P* < 0.001; \*\**P* < 0.01.

**Figure S2, related to Figure 2:** Absence of *Ggcx* induces a diabetic signature in islets. **(A-C)** Gene expression in *Ggcx^ff^; Pdx1-Cre* and *Ggcx^ff^* islets were analyzed by bulk RNA-sequencing and gene set enrichment analysis was performed for genes significantly modulated by *Ggcx* (false discovery rate (FDR) ≤ 0.05) using the **(A)** Gene Ontology (GO) biological processes terms, **(B)** KEGG pathways and **(C)** Keywords in the Uniprot database. Data are represented as –log_10_(FDR) and the number of genes associated to each pathway is indicated on the bar graphs. **(D)** Schematic representing a protein-protein interaction network associated to endoplasmic reticulum protein processing and response to stress, within the differentially expressed gene set (analyzed using StringDB). **(E)** Venn diagram representing the overlap between the transcriptome of *Ggcx^ff^; Pdx1-Cre* islets and islets from pre-diabetic (HFD for 8 weeks) or diabetic (*Lepr^db/db^* and *Ire1α^β-/-^*) mouse models. The top panel represents the overlap between the significantly up-regulated genes in all models, while the bottom panel represents the down-regulated genes. Number of genes is shown for each overlap. **(F)** Heat map representing 198 genes dysregulated in *Ggcx*-deficient islets and in the islets of at least one of 3 mouse models of diabetes or pre-diabetes (HFD, *db/db* and *Ire1^β-/-^*). Genes were next clustered according to enrichment for UniProt keywords (Glycoproteins, Plasma membrane, Signal secreted), Gene Ontology biological process (Response to ER stress) or KEGG pathway (NFkappa B and TNF signaling pathway).

**Figure S3, related to Figure 2:** Characterization of the *Ggcx^ff^; Pdx1-Cre* mouse model. **(A-C)** Metabolic parameters of 32-weeks old *Ggcx^ff^; Pdx1-Cre* and *Ggcx^ff^*male mice (n=10). **(A)** Energy expenditure, O_2_ consumption, CO_2_ production, **(B)** physical activity (x, y and z axis) and **(C)** food intake during day and night was measured using a continuous metabolic system. **(D)** Pancreas weight from 24- to 28-weeks old male mice was determined in fed condition (n=9-13). **(E)** Body weight of *Ggcx^ff^; Pdx1-Cre* and *Ggcx^ff^* male mice fed a chow diet was measured weekly. **(F)** Genomic DNA from various tissues from *Ggcx^ff^; Pdx1-Cre* mice was extracted and used to amplify *Ggcx* by PCR to detect the floxed and excised allele (Δ). **(G)** Genomic DNA from various tissues from *Vkorc1^ff^; Vkorc1l1^ff^; Pdx1-Cre* mice was extracted and used to amplify *Vkorc1* and *Vkorc1l1* by PCR to detect the floxed and excised allele (Δ). **(H)** *Vkorc1* and *Vkorc1l1* gene expression was analyzed by quantitative PCR in islets from *Vkorc1^ff^; Vkorc1l1^ff^; Pdx1-Cre* (*c1^ff^; c1l1^ff^; Pdx1-Cre*) mice (n=4) and data were normalized to *Actb*. **(I)** Islets from *Vkorc1^ff^; Vkorc1l1^ff^; Pdx1-Cre* and *Vkorc1^ff^; Vkorc1l1^ff^* littermates were harvested and γ-carboxylation was analysed by western blot using anti-Gla antibodies. β-actin was used as a loading control. Arrows indicate carboxylated proteins, while asterisks indicate non-specific bands. **(J-K)** Histomorphometric analysis on pancreas section following insulin staining and hematoxylin counterstaining from **(J)** *Vkorc1^ff^; Vkorc1l1^ff^; Pdx1-Cre* (n=6-9) and **(K)** *Pdx1-Cre* (n=6) 24-weeks old male mice. **(L)** Pancreas from 24-weeks old *Pdx1-Cre* and WT male mice were homogenized, and insulin content measured by ELISA (n=10). Results represent the mean ± SEM; Two-way ANOVA with Bonferroni’s multiple comparisons test was used for (A-C) and (E); Unpaired, two-tailed Student’s *t* test was used in (D), (H) and (J-L); \*\*\**P* < 0.001; \**P* < 0.05.

**Figure S4, related to Figure 3:** 7 days of high fat feeding induces ER-stress in islets. **(A-F)** *Ggcx^ff^; Pdx1-Cre* and *Ggcx^ff^* control mice were fed either a control low fat, or a high fat diet for 7 days and gene expression in islets was analyzed by qPCR. Data were normalized to *Actb* (n=4-6; *P* values for 2-way ANOVA are indicated when <0.05). **(G-H)** *Ggcx^ff^; Pdx1-Cre* and *Ggcx^ff^*male mice were fed with a control low-fat diet or HFD for 7 days (n=4-13). **(G)** Body weight and **(H)** glucose tolerance (IPGTT) was measured. **(I-K)** Metabolic analysis of 12-weeks old *Ggcx^ff^; Ins1-Cre* and *Ins1-Cre* mice on a regular chow diet (n=6-12). **(I)** IPGTT, **(J)** fasting glucose and **(K)** fasting insulin were measured. **(L-N)** *Ggcx^ff^; Ins1-Cre* and *Ggcx^ff^* male mice were fed a HFD for 7 days (n=7-13). **(L)** Body weight, **(M)** glucose tolerance (IPGTT) and **(N)** oral glucose tolerance (OGTT) was measured. Results represent the mean ± SEM; 2-way ANOVA with Bonferroni’s post-test was used in (A-I) and (M-N); Unpaired, 2-tailed Student’s *t* test was used in (J-L); \*\*\**P* < 0.001; \*\**P* < 0.01; \**P* < 0.05.

**Figure S5, related to Figure 4:** Low dose of thapsigargin activates the unfolded protein response in β**-cells. (A-B)** INS-1 832/3 cells were cultured with VK_1_ (22μM) or vehicle for 48 hours before being cultured for 24 hours in media containing 25mM glucose and 20nM thapsigargin. **(A)** Western blot was performed to analyze the unfolded protein response using anti-phospho(S724)-IRE1 and anti-GRP78/BiP antibodies. β-actin was used as a loading control. **(B)** Quantification was performed using arbitrary densitometry units of phospho(S724)-IRE1 and GRP78/BiP signals over β-actin signals. Results represent the mean ± SEM.

**Figure S6, related to Figure 5:** ERGP is the predominant *Asph* isoform in pancreatic islets. **(A-C)** Gene expression analysis of mouse islets using RNA-sequencing. **(A)** Expression level of Gla proteins encoding genes in control mouse islets (expressed as read counts normalized to library size). **(B)** Schematic representation of the mouse *Asph* gene locus and of the major *Asph* isoforms with β-cell expression level of each exon from the locus based on previously published mouse RNAseq data (DiGruccio et al., 2016). Below is a schematic representation of the proteins encoded by the *Asph* isoforms. Ser rich: serine rich domain; TM: transmembrane domain; EF: EF hand domain; Glu rich domain: glutamic acid rich domain; Lys/Arg rich domain: lysine and arginine rich domain. **(C)** Median read counts in promoter 1 and 2 of the *Asph* locus and in the final exon encoding ASPH, ERGP, junctate, and junctin. **(D)** Schematic representation of full length ASPH and ERGP with their respective deletion mutants. **(E)** HEK293 cells transfected with the indicated constructs were cultured with VK_1_ (22μM) or warfarin (50μM) as specified. ASPH and ERGP were detected by western blot using anti-A/E-GRD antibodies and anti-FLAG was used as a loading control. **(F-G)** ASPH and ERGP γ-carboxylation was assessed in *Vkorc1^+/+^* and *Vkorc1^-/-^* 7-day-old mouse livers by **(F)** immunoprecipitation with anti-Gla antibody followed by western blot with anti-A/E-GRD antibodies and by **(G)** immunoprecipitation with anti-A/E-GRD antibodies followed by western blot with anti-Gla antibodies. **(H)** ASPH and ERGP γ-carboxylation was assessed in *Vkorc1^+/-^; APOE-Vkorc1l1* and *Vkorc1^-/-^; APOE-Vkorc1l1* mouse islets by immunoprecipitation with anti-Gla antibody followed by western blot with anti-A/H-GRD antibodies (longer exposure compared to Fig. 6A). **(I)** Representative LC-MS/MS spectrum of the LGVYDADGDGDFDVDDAK peptide contained in ERGP and identified following anti-Gla immunoprecipitation on cell extracts from INS-1 832/3 grown in presence of VK_1_ and treated for 24h with 25mM glucose and 20nM thapsigargin. **(J)** HEK293 cells transfected with the indicated constructs were cultured with VK_1_ (22μM) or warfarin (50μM) as specified. FLAG-tagged proteins were immunoprecipitated with anti-FLAG agarose beads followed by western blot with anti-Gla antibodies. Western blot with anti-FLAG antibodies was used as a loading control. **(K)** Sequence alignment of GRD from mammalian ERGP homologues. Sequences from mouse (*M. musculus*), rat (*R. norvegicus*), human (*H. sapiens*), cat (*F. catus*), horse (*H. caballus*), cattle (*B. taurus*), wild boar (*S. scrofa*) and blue whale (*B. musculus*) are shown. Glutamic acid residues are highlighted in green; single asterisk (*) indicates a fully conserved residue; a colon (:) indicates a strongly conserved residue; and a period (.) indicates moderate or weak conservation. **(L)** HEK293 cells transfected with the indicated constructs were cultured with VK_1_ (22μM) or warfarin (50μM) as specified. Carboxylated proteins were immunoprecipitated with anti-Gla antibodies followed by western blot with anti-FLAG antibodies. Results represent the mean ± SEM.

**Figure S7, related to Figure 6 and 7:** Gamma-carboxylation in β**-cells affects STIM1 puncta formation. (A-B)** Efficient knockout of ASPH and ERGP in parental (p) HEK293 cells was validated by **(A)** western blot and **(B)** qPCR analysis (n=4). Asterisks indicate non-specific binding. **(C)** *ASPH^-/-^* HEK293 cells were transfected with STIM1-Myc, Orai1-HA and ERGP-3XFLAG in presence or absence of 22μM VK_1_ as indicated. Expression and γ-carboxylation were monitored by western blot using anti-Myc, anti-HA, anti-FLAG and anti-Gla antibodies. **(D)** Quantification of the average D4ER F_524_/F_475_ ratio after the addition of 2mM Ca^2+^ and 3μM ionomycin for the last 300 seconds of recording and expressed as % of the baseline vehicle condition. Data are extracted from experiments shown in figure 6F (n=3 experimental replicates per condition, with data from 61-118 cells averaged for each replicate). **(E)** *ASPH^-/-^* HEK293 cells were transfected with STIM1-CFP, YFP-Orai1 and ERGP-3XFLAG in presence or absence of 22μM VK_1_ as indicated. Expression of STIM1-CFP and YFP-Orai1 was monitored by western blot using anti-GFP antibodies recognizing both CFP and YFP. β-actin was used as a loading control. Quantification was performed using arbitrary densitometry units of STIM1-CFP signals over YFP-Orai1 signals. **(F)** Average change in F_509_/F_660_ ratio from baseline (average T_0-120sec_) to peak after 10μM thapsigargin in *Ggcx^ff^; Ins1-Cre* and *Ins1-Cre* islets (n=8 experimental replicates per genotypes, with data from 109-182 islet cell clusters averaged for each replicate). **(G)** Quantification of STIM1 puncta in *Ggcx^ff^; Ins1-Cre* and *Ins1-Cre* islets at steady-state (5mM glucose). Data are represented as number of puncta per μm^2^ (n=116-183 cells). **(H)** Representative confocal immunofluorescence images of *Ggcx^ff^; Ins1-Cre* and *Ins1-Cre* islets fixed and stained with anti-STIM1 antibodies. DAPI was used to stain nuclei. Scale bar: 10μm. Results represent the mean ± SEM; Unpaired two-tailed Student’s *t* test was used in (B), (F) and (G); Ordinary one-way ANOVA with Bonferroni’s post-tests was used in (D); \*\*\**P* < 0.001; \**P* < 0.05.

## REFERENCES

Al Rifai, O., Chow, J., Lacombe, J., Julien, C., Faubert, D., Susan-Resiga, D., Essalmani, R., Creemers, J.W., Seidah, N.G., and Ferron, M. (2017). Proprotein convertase furin regulates osteocalcin and bone endocrine function. J Clin Invest 127, 4104–4117.

Alquier, T., and Poitout, V. (2018). Considerations and guidelines for mouse metabolic phenotyping in diabetes research. Diabetologia 61, 526–538.

Bandyopadhyay, P.K., Garrett, J.E., Shetty, R.P., Keate, T., Walker, C.S., and Olivera, B.M. (2002). gamma -Glutamyl carboxylation: An extracellular posttranslational modification that antedates the divergence of molluscs, arthropods, and chordates. Proc Natl Acad Sci U S A 99, 1264–1269.

Berkner, K.L., and Pudota, B.N. (1998). Vitamin K-dependent carboxylation of the carboxylase. Proc Natl Acad Sci U S A 95, 466–471.

Beulens, J.W., van der, A.D., Grobbee, D.E., Sluijs, I., Spijkerman, A.M., and van der Schouw, Y.T. (2010). Dietary phylloquinone and menaquinones intakes and risk of type 2 diabetes. Diabetes Care 33, 1699–1705.

Brouwers, B., de Faudeur, G., Osipovich, A.B., Goyvaerts, L., Lemaire, K., Boesmans, L., Cauwelier, E.J., Granvik, M., Pruniau, V.P., Van Lommel, L., et al. (2014). Impaired islet function in commonly used transgenic mouse lines due to human growth hormone minigene expression. Cell Metab 20, 979–990.

Chabosseau, P., and Rutter, G.A. (2016). Zinc and diabetes. Archives of biochemistry and biophysics 611, 79–85.

Cheung, C.L., Sing, C.W., Lau, W.C.Y., Li, G.H.Y., Lip, G.Y.H., Tan, K.C.B., Cheung, B.M.Y., Chan, E.W.Y., and Wong, I.C.K. (2021). Treatment with direct oral anticoagulants or warfarin and the risk for incident diabetes among patients with atrial fibrillation: a population-based cohort study. Cardiovascular diabetology 20, 71.

Chomczynski, P., and Sacchi, N. (2006). The single-step method of RNA isolation by acid guanidinium thiocyanate-phenol-chloroform extraction: twenty-something years on. Nature protocols 1, 581–585.

Consortium, T.M. (2018). Single-cell transcriptomics of 20 mouse organs creates a Tabula Muris. Nature 562, 367–372.

DiGruccio, M.R., Mawla, A.M., Donaldson, C.J., Noguchi, G.M., Vaughan, J., Cowing-Zitron, C., van der Meulen, T., and Huising, M.O. (2016). Comprehensive alpha, beta and delta cell transcriptomes reveal that ghrelin selectively activates delta cells and promotes somatostatin release from pancreatic islets. Mol Metab 5, 449–458.

Dihingia, A., Ozah, D., Ghosh, S., Sarkar, A., Baruah, P.K., Kalita, J., Sil, P.C., and Manna, P. (2018). Vitamin K1 inversely correlates with glycemia and insulin resistance in patients with type 2 diabetes (T2D) and positively regulates SIRT1/AMPK pathway of glucose metabolism in liver of T2D mice and hepatocytes cultured in high glucose. The Journal of nutritional biochemistry 52, 103–114.

Dinchuk, J.E., Henderson, N.L., Burn, T.C., Huber, R., Ho, S.P., Link, J., O’Neil, K.T., Focht, R.J., Scully, M.S., Hollis, J.M., et al. (2000). Aspartyl beta -hydroxylase (Asph) and an evolutionarily conserved isoform of Asph missing the catalytic domain share exons with junctin. J Biol Chem 275, 39543–39554.

Ewang-Emukowhate, M., Harrington, D.J., Botha, A., McGowan, B., and Wierzbicki, A.S. (2015). Vitamin K and other markers of micronutrient status in morbidly obese patients before bariatric surgery. Int J Clin Pract 69, 638–642.

Ferdaoussi, M., Dai, X., Jensen, M.V., Wang, R., Peterson, B.S., Huang, C., Ilkayeva, O., Smith, N., Miller, N., Hajmrle, C., et al. (2015). Isocitrate-to-SENP1 signaling amplifies insulin secretion and rescues dysfunctional beta cells. J Clin Invest 125, 3847–3860.

Feriotto, G., Finotti, A., Breveglieri, G., Treves, S., Zorzato, F., and Gambari, R. (2006). Multiple levels of control of the expression of the human A beta H-J-J locus encoding aspartyl-beta-hydroxylase, junctin, and junctate. Ann N Y Acad Sci 1091, 184–190.

Feriotto, G., Finotti, A., Volpe, P., Treves, S., Ferrari, S., Angelelli, C., Zorzato, F., and Gambari, R. (2005). Myocyte enhancer factor 2 activates promoter sequences of the human AbetaH-J-J locus, encoding aspartyl-beta-hydroxylase, junctin, and junctate. Mol Cell Biol 25, 3261–3275.

Ferron, M., Boudiffa, M., Arsenault, M., Rached, M., Pata, M., Giroux, S., Elfassihi, L., Kisseleva, M., Majerus, P.W., Rousseau, F., et al. (2011). Inositol polyphosphate 4-phosphatase B as a regulator of bone mass in mice and humans. Cell Metab 14, 466–477.

Ferron, M., Lacombe, J., Germain, A., Oury, F., and Karsenty, G. (2015). GGCX and VKORC1 inhibit osteocalcin endocrine functions. J Cell Biol 208, 761–776.

Furie, B., Bouchard, B.A., and Furie, B.C. (1999). Vitamin K-dependent biosynthesis of gamma-carboxyglutamic acid. Blood 93, 1798–1808.

Ghosh, S., Kraus, K., Biswas, A., Muller, J., Buhl, A.L., Forin, F., Singer, H., Honing, K., Hornung, V., Watzka, M., et al. (2021). GGCX mutations show different responses to vitamin K thereby determining the severity of the hemorrhagic phenotype in VKCFD1 patients. Journal of thrombosis and haemostasis : JTH 19, 1412–1424.

Greotti, E., Wong, A., Pozzan, T., Pendin, D., and Pizzo, P. (2016). Characterization of the ER-Targeted Low Affinity Ca(2+) Probe D4ER. Sensors (Basel) 16.

Hallgren, K.W., Zhang, D., Kinter, M., Willard, B., and Berkner, K.L. (2013). Methylation of gamma-carboxylated Glu (Gla) allows detection by liquid chromatography-mass spectrometry and the identification of Gla residues in the gamma-glutamyl carboxylase. Journal of proteome research 12, 2365–2374.

Haque, J.A., McDonald, M.G., Kulman, J.D., and Rettie, A.E. (2014). A cellular system for quantitation of vitamin K cycle activity: structure-activity effects on vitamin K antagonism by warfarin metabolites. Blood 123, 582–589.

Hingorani, S.R., Petricoin, E.F., Maitra, A., Rajapakse, V., King, C., Jacobetz, M.A., Ross, S., Conrads, T.P., Veenstra, T.D., Hitt, B.A., et al. (2003). Preinvasive and invasive ductal pancreatic cancer and its early detection in the mouse. Cancer cell 4, 437–450.

Hoffman, D.J., Powell, T.L., Barrett, E.S., and Hardy, D.B. (2021). Developmental origins of metabolic diseases. Physiol Rev 101, 739–795.

Hudish, L.I., Reusch, J.E., and Sussel, L. (2019). beta Cell dysfunction during progression of metabolic syndrome to type 2 diabetes. J Clin Invest 129, 4001–4008.

Ibarrola-Jurado, N., Salas-Salvado, J., Martinez-Gonzalez, M.A., and Bullo, M. (2012). Dietary phylloquinone intake and risk of type 2 diabetes in elderly subjects at high risk of cardiovascular disease. Am J Clin Nutr 96, 1113–1118.

Johnson, J.S., Kono, T., Tong, X., Yamamoto, W.R., Zarain-Herzberg, A., Merrins, M.J., Satin, L.S., Gilon, P., and Evans-Molina, C. (2014). Pancreatic and duodenal homeobox protein 1 (Pdx-1) maintains endoplasmic reticulum calcium levels through transcriptional regulation of sarco-endoplasmic reticulum calcium ATPase 2b (SERCA2b) in the islet beta cell. J Biol Chem 289, 32798–32810.

Kaidar-Person, O., Person, B., Szomstein, S., and Rosenthal, R.J. (2008). Nutritional deficiencies in morbidly obese patients: a new form of malnutrition? Part A: vitamins. Obesity surgery 18, 870–876.

Karamzad, N., Faraji, E., Adeli, S., Carson-Chahhoud, K., Azizi, S., and Pourghassem Gargari, B. (2020). Effects of MK-7 Supplementation on Glycemic Status, Anthropometric Indices and Lipid Profile in Patients with Type 2 Diabetes: A Randomized Controlled Trial. Diabetes Metab Syndr Obes 13, 2239–2249.

Kono, T., Tong, X., Taleb, S., Bone, R.N., Iida, H., Lee, C.C., Sohn, P., Gilon, P., Roe, M.W., and Evans-Molina, C. (2018). Impaired Store-Operated Calcium Entry and STIM1 Loss Lead to Reduced Insulin Secretion and Increased Endoplasmic Reticulum Stress in the Diabetic beta-Cell. Diabetes 67, 2293–2304.

Kwon, S.J., and Kim, D.H. (2009). Characterization of junctate-SERCA2a interaction in murine cardiomyocyte. Biochem Biophys Res Commun 390, 1389–1394.

Lacombe, J., Al Rifai, O., Loter, L., Moran, T., Turcotte, A.F., Grenier-Larouche, T., Tchernof, A., Biertho, L., Carpentier, A.C., Prud’homme, D., et al. (2020). Measurement of bioactive osteocalcin in humans using a novel immunoassay reveals association with glucose metabolism and beta-cell function. American journal of physiology Endocrinology and metabolism 318, E381–E391.

Lacombe, J., and Ferron, M. (2018). VKORC1L1, An Enzyme Mediating the Effect of Vitamin K in Liver and Extrahepatic Tissues. Nutrients 10, E970.

Lacombe, J., Rishavy, M.A., Berkner, K.L., and Ferron, M. (2018). VKOR paralog VKORC1L1 supports vitamin K-dependent protein carboxylation in vivo. JCI Insight 3, e96501.

Lee, H., Lee, Y.S., Harenda, Q., Pietrzak, S., Oktay, H.Z., Schreiber, S., Liao, Y., Sonthalia, S., Ciecko, A.E., Chen, Y.G., et al. (2020). Beta Cell Dedifferentiation Induced by IRE1alpha Deletion Prevents Type 1 Diabetes. Cell Metab 31, 822–836 e825

Lee, N.K., Sowa, H., Hinoi, E., Ferron, M., Ahn, J.D., Confavreux, C., Dacquin, R., Mee, P.J., McKee, M.D., Jung, D.Y., et al. (2007). Endocrine regulation of energy metabolism by the skeleton. Cell 130, 456–469.

Liang, K., Du, W., Lu, J., Li, F., Yang, L., Xue, Y., Hille, B., and Chen, L. (2014). Alterations of the Ca(2)(+) signaling pathway in pancreatic beta-cells isolated from db/db mice. Protein Cell 5, 783–794.

Liu, X., Feng, S., Chen, Z., Zhou, Y., Yin, K., Xue, Z., and Zhu, W. (2022). Is the Risk of Diabetes Lower in Patients With Atrial Fibrillation Treated With Direct Oral Anticoagulant Compared to Warfarin? Front Cardiovasc Med 9, 874795.

Lunz, V., Romanin, C., and Frischauf, I. (2019). STIM1 activation of Orai1. Cell calcium 77, 29–38.

Lyon, J., Manning Fox, J.E., Spigelman, A.F., Kim, R., Smith, N., O’Gorman, D., Kin, T., Shapiro, A.M., Rajotte, R.V., and MacDonald, P.E. (2016). Research-Focused Isolation of Human Islets From Donors With and Without Diabetes at the Alberta Diabetes Institute IsletCore. Endocrinology 157, 560–569.

Lyon, J., Spigelman, A.F., MacDonald, P.E., and Manning Fox, J.E. (2019). ADI IsletCore Protocols for the Isolation, Assessment and Cryopreservation of Human Pancreatic Islets of Langerhans for Research Purposes V.1. protocolio.

Mamenko, M., Dhande, I., Tomilin, V., Zaika, O., Boukelmoune, N., Zhu, Y., Gonzalez-Garay, M.L., Pochynyuk, O., and Doris, P.A. (2016). Defective Store-Operated Calcium Entry Causes Partial Nephrogenic Diabetes Insipidus. J Am Soc Nephrol 27, 2035–2048.

Mehran, A.E., Templeman, N.M., Brigidi, G.S., Lim, G.E., Chu, K.Y., Hu, X., Botezelli, J.D., Asadi, A., Hoffman, B.G., Kieffer, T.J., et al. (2012). Hyperinsulinemia drives diet-induced obesity independently of brain insulin production. Cell Metab 16, 723–737.

Mittendorfer, B., Patterson, B.W., Smith, G.I., Yoshino, M., and Klein, S. (2022). beta Cell function and plasma insulin clearance in people with obesity and different glycemic status. J Clin Invest 132.

Motterle, A., Gattesco, S., Peyot, M.L., Esguerra, J.L.S., Gomez-Ruiz, A., Laybutt, D.R., Gilon, P., Burdet, F., Ibberson, M., Eliasson, L., et al. (2017). Identification of islet-enriched long non-coding RNAs contributing to beta-cell failure in type 2 diabetes. Mol Metab 6, 1407–1418.

Murshed, M., Schinke, T., McKee, M.D., and Karsenty, G. (2004). Extracellular matrix mineralization is regulated locally; different roles of two gla-containing proteins. J Cell Biol 165, 625–630.

Oropeza, D., Jouvet, N., Budry, L., Campbell, J.E., Bouyakdan, K., Lacombe, J., Perron, G., Bergeron, V., Neuman, J.C., Brar, H.K., et al. (2015). Phenotypic Characterization of MIP-CreERT1Lphi Mice With Transgene-Driven Islet Expression of Human Growth Hormone. Diabetes 64, 3798–3807.

Palty, R., Raveh, A., Kaminsky, I., Meller, R., and Reuveny, E. (2012). SARAF inactivates the store operated calcium entry machinery to prevent excess calcium refilling. Cell 149, 425–438.

Pan, Y., and Jackson, R.T. (2009). Dietary phylloquinone intakes and metabolic syndrome in US young adults. J Am Coll Nutr 28, 369–379.

Park, C.Y., Hoover, P.J., Mullins, F.M., Bachhawat, P., Covington, E.D., Raunser, S., Walz, T., Garcia, K.C., Dolmetsch, R.E., and Lewis, R.S. (2009). STIM1 clusters and activates CRAC channels via direct binding of a cytosolic domain to Orai1. Cell 136, 876–890.

Perez-Riverol, Y., Bai, J., Bandla, C., Garcia-Seisdedos, D., Hewapathirana, S., Kamatchinathan, S., Kundu, D.J., Prakash, A., Frericks-Zipper, A., Eisenacher, M., et al. (2022). The PRIDE database resources in 2022: a hub for mass spectrometry-based proteomics evidences. Nucleic acids research 50, D543–D552.

Picard, M., Petrie, R.J., Antoine-Bertrand, J., Saint-Cyr-Proulx, E., Villemure, J.F., and Lamarche-Vane, N. (2009). Spatial and temporal activation of the small GTPases RhoA and Rac1 by the netrin-1 receptor UNC5a during neurite outgrowth. Cell Signal 21, 1961–1973.

Rahimi Sakak, F., Moslehi, N., Niroomand, M., and Mirmiran, P. (2021). Glycemic control improvement in individuals with type 2 diabetes with vitamin K2 supplementation: a randomized controlled trial. Eur J Nutr 60, 2495–2506.

Roe, M.W., Philipson, L.H., Frangakis, C.J., Kuznetsov, A., Mertz, R.J., Lancaster, M.E., Spencer, B., Worley, J.F., 3rd, and Dukes, I.D. (1994). Defective glucose-dependent endoplasmic reticulum Ca2+ sequestration in diabetic mouse islets of Langerhans. J Biol Chem 269, 18279–18282.

Ronnebaum, S.M., Jensen, M.V., Hohmeier, H.E., Burgess, S.C., Zhou, Y.P., Qian, S., MacNeil, D., Howard, A., Thornberry, N., Ilkayeva, O., et al. (2008). Silencing of cytosolic or mitochondrial isoforms of malic enzyme has no effect on glucose-stimulated insulin secretion from rodent islets. J Biol Chem 283, 28909–28917.

Sabatini, P.V., Speckmann, T., and Lynn, F.C. (2019). Friend and foe: beta-cell Ca(2+) signaling and the development of diabetes. Mol Metab 21, 1–12.

Sabourin, J., Le Gal, L., Saurwein, L., Haefliger, J.A., Raddatz, E., and Allagnat, F. (2015). Store-operated Ca2+ Entry Mediated by Orai1 and TRPC1 Participates to Insulin Secretion in Rat beta-Cells. J Biol Chem 290, 30530–30539.

Samtleben, S., Wachter, B., and Blum, R. (2015). Store-operated calcium entry compensates fast ER calcium loss in resting hippocampal neurons. Cell calcium 58, 147–159.

Schindelin, J., Arganda-Carreras, I., Frise, E., Kaynig, V., Longair, M., Pietzsch, T., Preibisch, S., Rueden, C., Saalfeld, S., Schmid, B., et al. (2012). Fiji: an open-source platform for biological-image analysis. Nat Methods 9, 676–682.

Schmidt, T., Samaras, P., Frejno, M., Gessulat, S., Barnert, M., Kienegger, H., Krcmar, H., Schlegl, J., Ehrlich, H.C., Aiche, S., et al. (2018). ProteomicsDB. Nucleic acids research 46, D1271–D1281.

Schulte, A., and Blum, R. (2022). Shaped by leaky ER: Homeostatic Ca(2+) fluxes. Front Physiol 13, 972104.

Sharma, R.B., Landa-Galvan, H.V., and Alonso, L.C. (2021). Living Dangerously: Protective and Harmful ER Stress Responses in Pancreatic beta-Cells. Diabetes 70, 2431–2443.

Sharma, R.B., O’Donnell, A.C., Stamateris, R.E., Ha, B., McCloskey, K.M., Reynolds, P.R., Arvan, P., and Alonso, L.C. (2015). Insulin demand regulates beta cell number via the unfolded protein response. J Clin Invest 125, 3831–3846.

Shen, G., Cui, W., Zhang, H., Zhou, F., Huang, W., Liu, Q., Yang, Y., Li, S., Bowman, G.R., Sadler, J.E.*, et al.* (2017). Warfarin traps human vitamin K epoxide reductase in an intermediate state during electron transfer. Nat Struct Mol Biol 24, 69–76.

Solis-Herrera, C., Triplitt, C., Cersosimo, E., and DeFronzo, R.A. (2000). Pathogenesis of Type 2 Diabetes Mellitus. In Endotext, K.R. Feingold, B. Anawalt, A. Boyce, G. Chrousos, W.W. de Herder, K. Dhatariya, K. Dungan, J.M. Hershman, J. Hofland, S. Kalra, et al., eds. (South Dartmouth (MA)).

Srikanth, S., Jew, M., Kim, K.D., Yee, M.K., Abramson, J., and Gwack, Y. (2012). Junctate is a Ca2+-sensing structural component of Orai1 and stromal interaction molecule 1 (STIM1). Proc Natl Acad Sci U S A 109, 8682–8687.

Srour, B., Fezeu, L.K., Kesse-Guyot, E., Alles, B., Debras, C., Druesne-Pecollo, N., Chazelas, E., Deschasaux, M., Hercberg, S., Galan, P.*, et al.* (2020). Ultraprocessed Food Consumption and Risk of Type 2 Diabetes Among Participants of the NutriNet-Sante Prospective Cohort. JAMA Intern Med 180, 283–291.

Stamateris, R.E., Sharma, R.B., Hollern, D.A., and Alonso, L.C. (2013). Adaptive beta-cell proliferation increases early in high-fat feeding in mice, concurrent with metabolic changes, with induction of islet cyclin D2 expression. American journal of physiology Endocrinology and metabolism 305, E149–159.

Stenflo, J., Fernlund, P., Egan, W., and Roepstorff, P. (1974). Vitamin K dependent modifications of glutamic acid residues in prothrombin. Proc Natl Acad Sci U S A 71, 2730–2733.

Thorens, B., Tarussio, D., Maestro, M.A., Rovira, M., Heikkila, E., and Ferrer, J. (2015). Ins1(Cre) knock-in mice for beta cell-specific gene recombination. Diabetologia 58, 558–565.

Tie, J.K., Jin, D.Y., Straight, D.L., and Stafford, D.W. (2011). Functional study of the vitamin K cycle in mammalian cells. Blood 117, 2967–2974.

Treves, S., Feriotto, G., Moccagatta, L., Gambari, R., and Zorzato, F. (2000). Molecular cloning, expression, functional characterization, chromosomal localization, and gene structure of junctate, a novel integral calcium binding protein of sarco(endo)plasmic reticulum membrane. J Biol Chem 275, 39555–39568.

Treves, S., Franzini-Armstrong, C., Moccagatta, L., Arnoult, C., Grasso, C., Schrum, A., Ducreux, S., Zhu, M.X., Mikoshiba, K., Girard, T.*, et al.* (2004). Junctate is a key element in calcium entry induced by activation of InsP3 receptors and/or calcium store depletion. J Cell Biol 166, 537–548.

Via, M. (2012). The malnutrition of obesity: micronutrient deficiencies that promote diabetes. ISRN endocrinology 2012, 103472.

Wang, I.M., Zhang, B., Yang, X., Zhu, J., Stepaniants, S., Zhang, C., Meng, Q., Peters, M., He, Y., Ni, C.*, et al.* (2012). Systems analysis of eleven rodent disease models reveals an inflammatome signature and key drivers. Mol Syst Biol 8, 594.

Weldemariam, M.M., Woo, J., and Zhang, Q. (2022). Pancreatic INS-1 beta-Cell Response to Thapsigargin and Rotenone: A Comparative Proteomics Analysis Uncovers Key Pathways of beta-Cell Dysfunction. Chemical research in toxicology 35, 1080–1094.

Xia, Z., and Liu, Y. (2001). Reliable and global measurement of fluorescence resonance energy transfer using fluorescence microscopes. Biophys J 81, 2395–2402.

Yong, J., Parekh, V.S., Reilly, S.M., Nayak, J., Chen, Z., Lebeaupin, C., Jang, I., Zhang, J., Prakash, T.P., Sun, H.*, et al.* (2021). Chop/Ddit3 depletion in beta cells alleviates ER stress and corrects hepatic steatosis in mice. Sci Transl Med 13.

Zhang, I.X., Ren, J., Vadrevu, S., Raghavan, M., and Satin, L.S. (2020). ER stress increases store-operated Ca(2+) entry (SOCE) and augments basal insulin secretion in pancreatic beta cells. J Biol Chem 295, 5685–5700.

Zwakenberg, S.R., Remmelzwaal, S., Beulens, J.W.J., Booth, S.L., Burgess, S., Dashti, H.S., Imamura, F., Feskens, E.J.M., van der Schouw, Y.T., and Sluijs, I. (2019). Circulating Phylloquinone Concentrations and Risk of Type 2 Diabetes: A Mendelian Randomization Study. Diabetes 68, 220–225.

