## Supplementary material for "Vitamin K-dependent γ-carboxylation regulates calcium flux and adaptation to metabolic stress in β-cells": Figures S1-S7; Tables S1, S3, S4 and S5

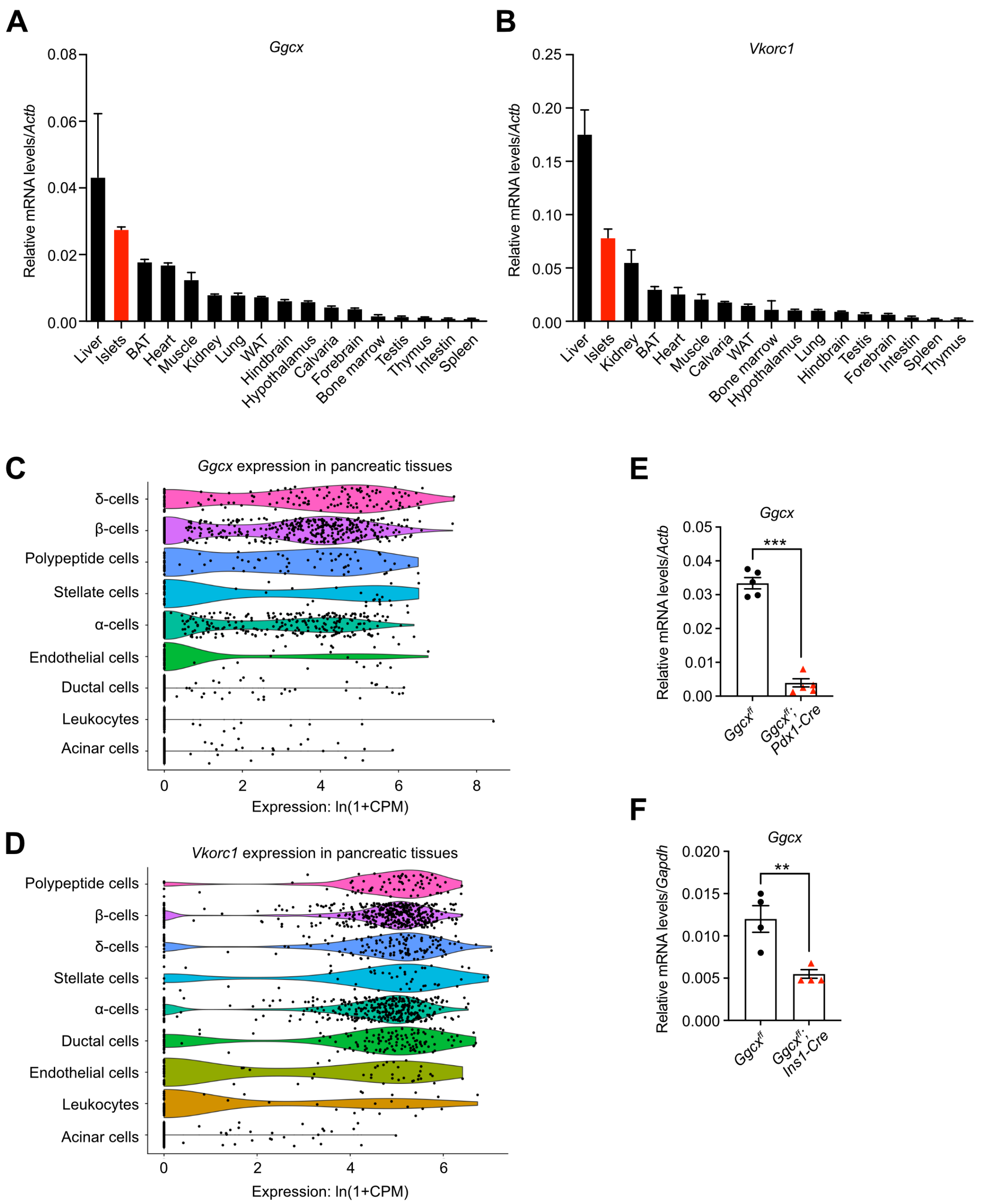


**Figure S1, related to Figure 1: *Ggcx* and *Vkorc1* are expressed in pancreatic endocrine cells. (A-B)** *Ggcx* and *Vkorc1* gene expression was analyzed by quantitative PCR in various tissues from wild-type (WT) mice and normalized to *Actb*. **(C-D)** Violin plots representing single cell transcriptome data from mouse pancreatic tissues (https://tabula-muris.ds.czbiohub.org/). **(E-F)** *Ggcx* gene expression was analyzed by quantitative PCR in islets from **(E)** *Ggcx$^{ff}$; Pdx1-Cre*, **(F)** *Ggcx$^{ff}$; Ins1-Cre* and their respective *Ggcx$^{ff}$* controls and normalized to *Actb* or *Gapdh* (n=4-5). Results represent the mean ± SEM; Unpaired, 2-tailed Student's *t* test was used in (E-F). ***$P < 0.001$; **$P < 0.01$.

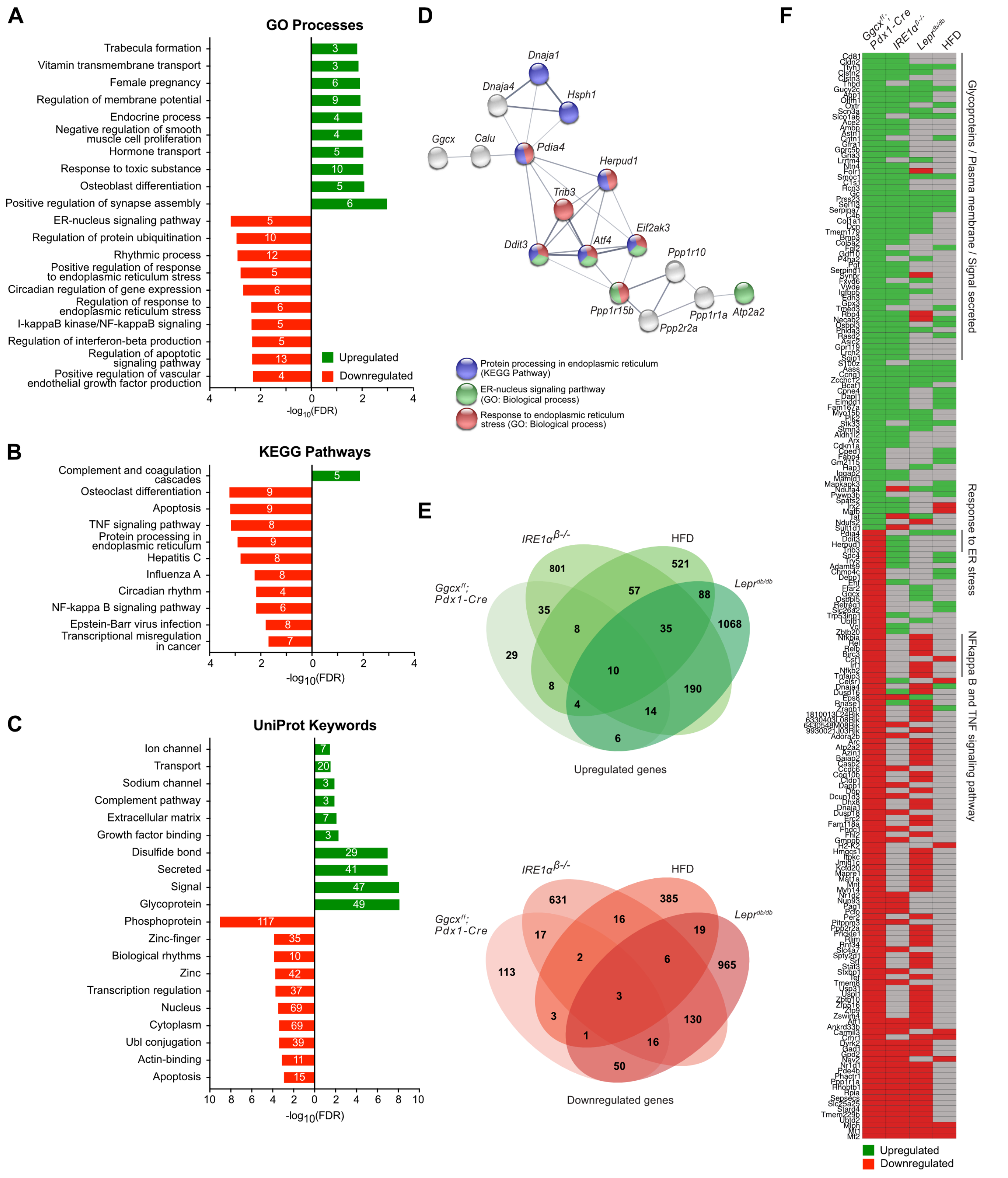


**Figure S2, related to Figure 2: Absence of *Ggcx* induces a diabetic signature in islets. (A-C)** Gene expression in *Ggcx$^{ff}$; Pdx1-Cre* and *Ggcx$^{ff}$* islets were analyzed by bulk RNA-sequencing and gene set enrichment analysis was performed for genes significantly modulated by *Ggcx* (false discovery rate (FDR) ≤ 0.05) using the **(A)** Gene Ontology (GO) biological processes terms, **(B)** KEGG pathways and

**(C)** Keywords in the Uniprot database. Data are represented as $-\log_{10}$(FDR) and the number of genes associated to each pathway is indicated on the bar graphs. **(D)** Schematic representing a protein-protein interaction network associated to endoplasmic reticulum protein processing and response to stress, within the differentially expressed gene set (analyzed using StringDB). **(E)** Venn diagram representing the overlap between the transcriptome of $Ggcx^{ff}$*; Pdx1-Cre* islets and islets from pre-diabetic (HFD for 8 weeks) or diabetic ($Lepr^{db/db}$ and $Ire1\alpha^{\beta-/-}$) mouse models. The top panel represents the overlap between the significantly up-regulated genes in all models, while the bottom panel represents the down-regulated genes. Number of genes is shown for each overlap. **(F)** Heat map representing 198 genes dysregulated in *Ggcx*-deficient islets and in the islets of at least one of 3 mouse models of diabetes or pre-diabetes (HFD, *db/db* and $Ire1^{\beta-/-}$). Genes were next clustered according to enrichment for UniProt keywords (Glycoproteins, Plasma membrane, Signal secreted), Gene Ontology biological process (Response to ER stress) or KEGG pathway (NFkappa B and TNF signaling pathway).

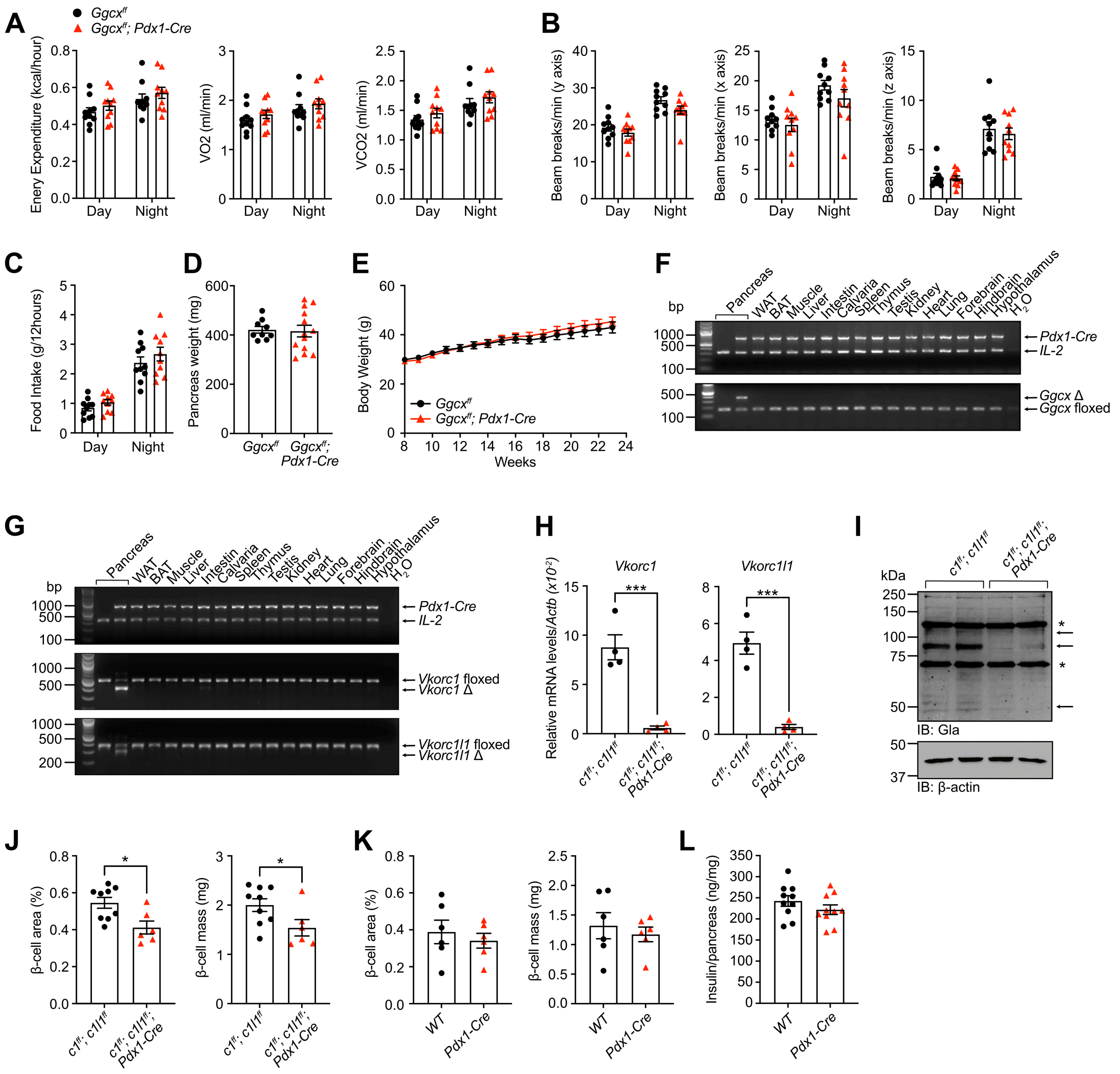


**Figure S3, related to Figure 2: Characterization of the *Ggcx*$^{ff}$*; Pdx1-Cre* mouse model. (A-C)** Metabolic parameters of 32-weeks old *Ggcx*$^{ff}$*; Pdx1-Cre* and *Ggcx*$^{ff}$ male mice (n=10). **(A)** Energy expenditure, $O_2$ consumption, $CO_2$ production, **(B)** physical activity (x, y and z axis) and **(C)** food intake during day and night was measured using a continuous metabolic system. **(D)** Pancreas weight from 24- to 28-weeks old male mice was determined in fed condition (n=9-13). **(E)** Body weight of *Ggcx*$^{ff}$*; Pdx1-Cre* and *Ggcx*$^{ff}$ male mice fed a chow diet was measured weekly. **(F)** Genomic DNA from various tissues from *Ggcx*$^{ff}$*; Pdx1-Cre* mice was extracted and used to amplify *Ggcx* by PCR to detect the floxed and excised allele (Δ). **(G)** Genomic DNA from various tissues from *Vkorc1*$^{ff}$*; Vkorc1l1*$^{ff}$*; Pdx1-Cre* mice was extracted and used to amplify *Vkorc1* and *Vkorc1l1* by PCR to detect the floxed and excised allele (Δ). **(H)** *Vkorc1* and *Vkorc1l1* gene expression was analyzed by quantitative PCR in islets from *Vkorc1$^{ff}$; Vkorc1l1$^{ff}$; Pdx1-Cre* (*c1$^{ff}$; c1l1$^{ff}$; Pdx1-Cre*) mice (n=4) and data were normalized to *Actb*. **(I)** Islets from *Vkorc1$^{ff}$; Vkorc1l1$^{ff}$; Pdx1-Cre* and *Vkorc1$^{ff}$; Vkorc1l1$^{ff}$* littermates were harvested and γ-carboxylation was analysed by western blot using anti-Gla antibodies. β-actin was used as a loading control. Arrows indicate carboxylated proteins, while asterisks indicate non-specific bands. **(J-K)** Histomorphometric analysis on pancreas section following insulin staining and hematoxylin counterstaining from **(J)** *Vkorc1$^{ff}$; Vkorc1l1$^{ff}$; Pdx1-Cre* (n=6-9) and **(K)** *Pdx1-Cre* (n=6) 24-weeks old male mice. **(L)** Pancreas from 24-weeks old *Pdx1-Cre* and WT male mice were homogenized, and insulin content measured by ELISA (n=10). Results represent the mean ± SEM; Two-way ANOVA with Bonferroni's multiple comparisons test was used for (A-C) and (E); Unpaired, two-tailed Student's *t* test was used in (D), (H) and (J-L); ***$P < 0.001$; *$P < 0.05$.

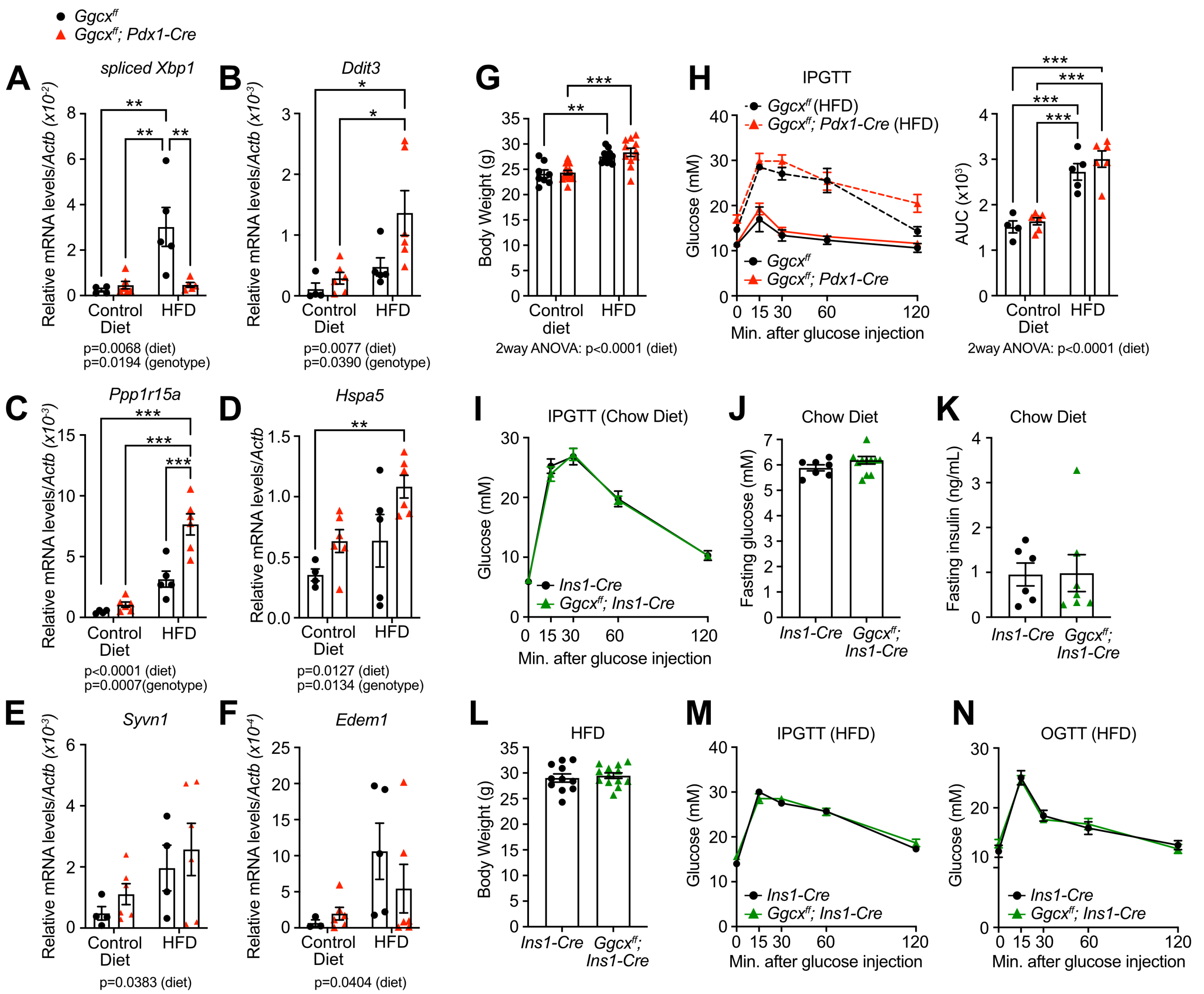


**Figure S4, related to Figure 3: 7 days of high fat feeding induces ER-stress in islets.** **(A-F)** *Ggcx$^{f/f}$; Pdx1-Cre* and *Ggcx$^{f/f}$* control mice were fed either a control low fat, or a high fat diet for 7 days and gene expression in islets was analyzed by qPCR. Data were normalized to *Actb* (n=4-6; *P* values for 2-way ANOVA are indicated when <0.05). **(G-H)** *Ggcx$^{f/f}$; Pdx1-Cre* and *Ggcx$^{f/f}$* male mice were fed with a control low-fat diet or HFD for 7 days (n=4-13). **(G)** Body weight and **(H)** glucose tolerance (IPGTT) was measured. **(I-K)** Metabolic analysis of 12-weeks old *Ggcx$^{f/f}$; Ins1-Cre* and *Ins1-Cre* mice on a regular chow diet (n=6-12). **(I)** IPGTT, **(J)** fasting glucose and **(K)** fasting insulin were measured. **(L-N)** *Ggcx$^{f/f}$; Ins1-Cre* and *Ggcx$^{f/f}$* male mice were fed a HFD for 7 days (n=7-13). **(L)** Body weight, **(M)** glucose tolerance (IPGTT) and **(N)** oral glucose tolerance (OGTT) was measured. Results represent the mean ± SEM; 2-way ANOVA with Bonferroni's post-test was used in (A-I) and (M-N); Unpaired, 2-tailed Student's *t* test was used in (J-L); ****P* < 0.001; ***P* < 0.01; **P* < 0.05.

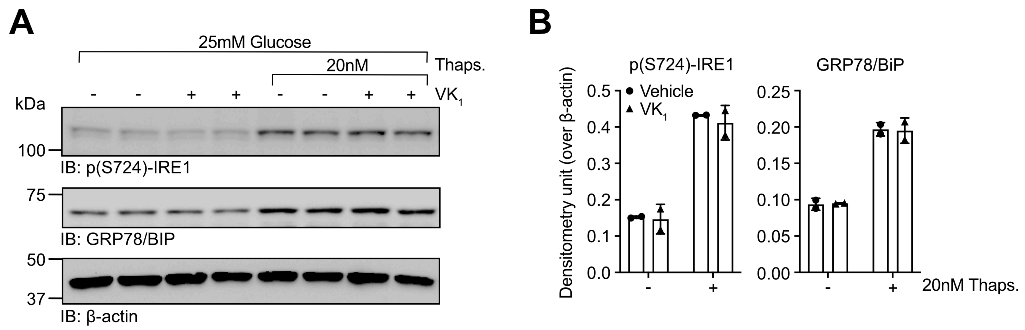


**Figure S5, related to Figure 4: Low dose of thapsigargin activates the unfolded protein response in β-cells. (A-B)** INS-1 832/3 cells were cultured with $VK_1$ (22µM) or vehicle for 48 hours before being cultured for 24 hours in media containing 25mM glucose and 20nM thapsigargin. **(A)** Western blot was performed to analyze the unfolded protein response using anti-phospho(S724)-IRE1 and anti-GRP78/BiP antibodies. β-actin was used as a loading control. **(B)** Quantification was performed using arbitrary densitometry units of phospho(S724)-IRE1 and GRP78/BiP signals over β-actin signals. Results represent the mean ± SEM.

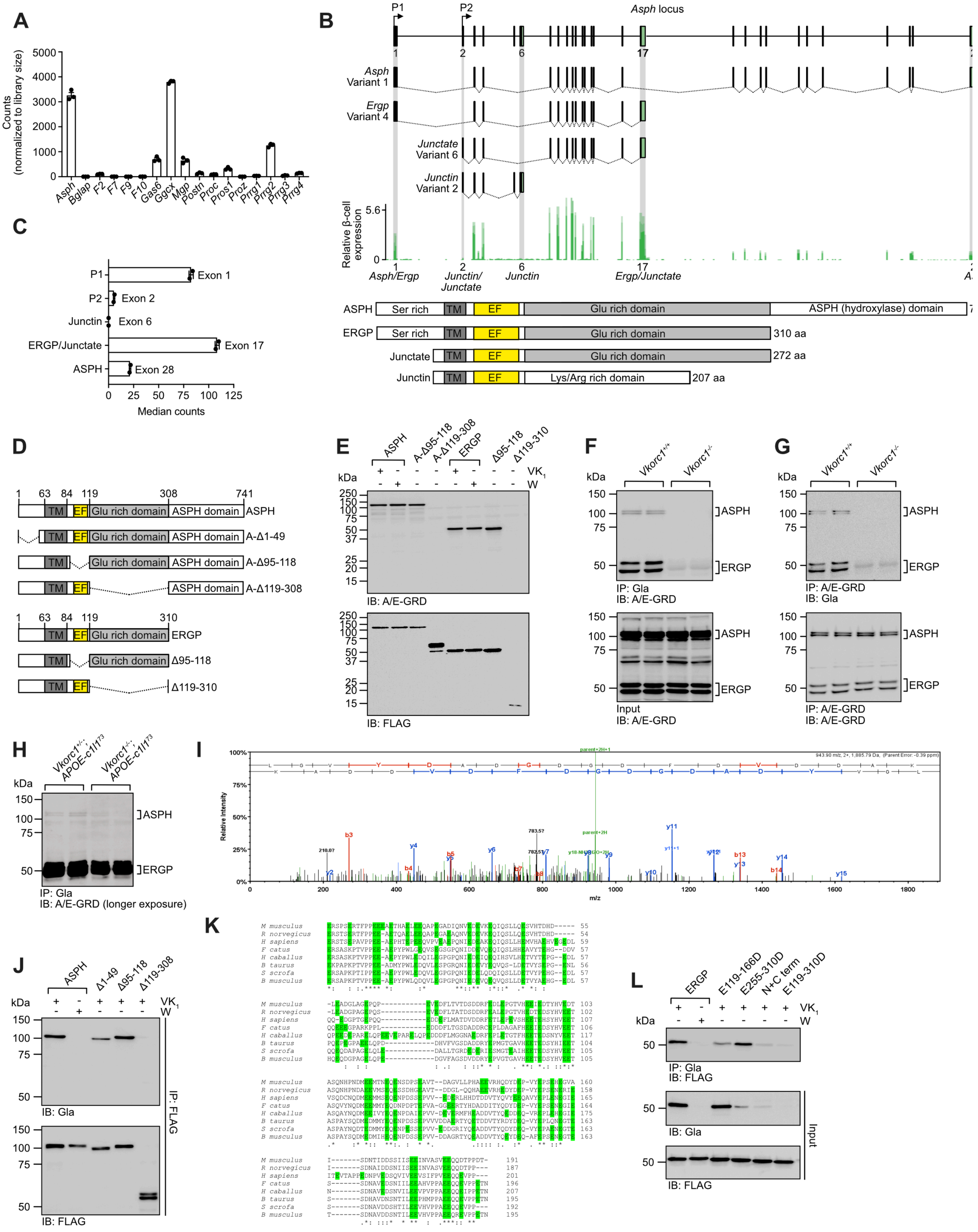


**Figure S6, related to Figure 5: ERGP is the predominant *Asph* isoform in pancreatic islets. (A-C)** Gene expression analysis of mouse islets using RNA-sequencing. **(A)** Expression level of Gla proteins encoding genes in control mouse islets (expressed as read counts normalized to library size). **(B)** Schematic representation of the mouse *Asph* gene locus and of the major *Asph* isoforms with β-cell expression level of each exon from the locus based on previously published mouse RNAseq data (DiGruccio et al., 2016). Below is a schematic representation of the proteins encoded by the *Asph* isoforms. Ser rich: serine rich domain; TM: transmembrane domain; EF: EF hand domain; Glu rich domain: glutamic acid rich domain; Lys/Arg rich domain: lysine and arginine rich domain. **(C)** Median read counts in promoter 1 and 2 of the *Asph* locus and in the final exon encoding ASPH, ERGP, junctate, and junctin. **(D)** Schematic representation of full length ASPH and ERGP with their respective deletion mutants. **(E)** HEK293 cells transfected with the indicated constructs were cultured with $VK_1$ (22μM) or warfarin (50μM) as specified. ASPH and ERGP were detected by western blot using anti-A/E-GRD antibodies and anti-FLAG was used as a loading control. **(F-G)** ASPH and ERGP γ-carboxylation was assessed in *Vkorc1*$^{+/+}$ and *Vkorc1*$^{-/-}$ 7-day-old mouse livers by **(F)** immunoprecipitation with anti-Gla antibody followed by western blot with anti-A/E-GRD antibodies and by **(G)** immunoprecipitation with anti-A/E-GRD antibodies followed by western blot with anti-Gla antibodies. **(H)** ASPH and ERGP γ-carboxylation was assessed in *Vkorc1*$^{+/-}$*; APOE-Vkorc1l1* and *Vkorc1*$^{-/-}$*; APOE-Vkorc1l1* mouse islets by immunoprecipitation with anti-Gla antibody followed by western blot with anti-A/H-GRD antibodies (longer exposure compared to Fig. 6A). **(I)** Representative LC-MS/MS spectrum of the LGVYDADGDGDFDVDDAK peptide contained in ERGP and identified following anti-Gla immunoprecipitation on cell extracts from INS-1 832/3 grown in presence of $VK_1$ and treated for 24h with 25mM glucose and 20nM thapsigargin. **(J)** HEK293 cells transfected with the indicated constructs were cultured with $VK_1$ (22μM) or warfarin (50μM) as specified. FLAG-tagged proteins were immunoprecipitated with anti-FLAG agarose beads followed by western blot with anti-Gla antibodies. Western blot with anti-FLAG antibodies was used as a loading control. **(K)** Sequence alignment of GRD from mammalian ERGP homologues. Sequences from mouse (*M. musculus*), rat (*R. norvegicus*), human (*H. sapiens*), cat (*F. catus*), horse (*H. caballus*), cattle (*B. taurus*), wild boar (*S. scrofa*) and blue whale (*B. musculus*) are shown. Glutamic acid residues are highlighted in green; single asterisk (*) indicates a fully conserved residue; a colon (:) indicates a strongly conserved residue; and a period (.) indicates moderate or weak conservation. **(L)** HEK293 cells transfected with the indicated constructs were cultured with $VK_1$ (22μM) or warfarin (50μM) as specified. Carboxylated proteins were immunoprecipitated with anti-Gla antibodies followed by western blot with anti-FLAG antibodies. Results represent the mean ± SEM.

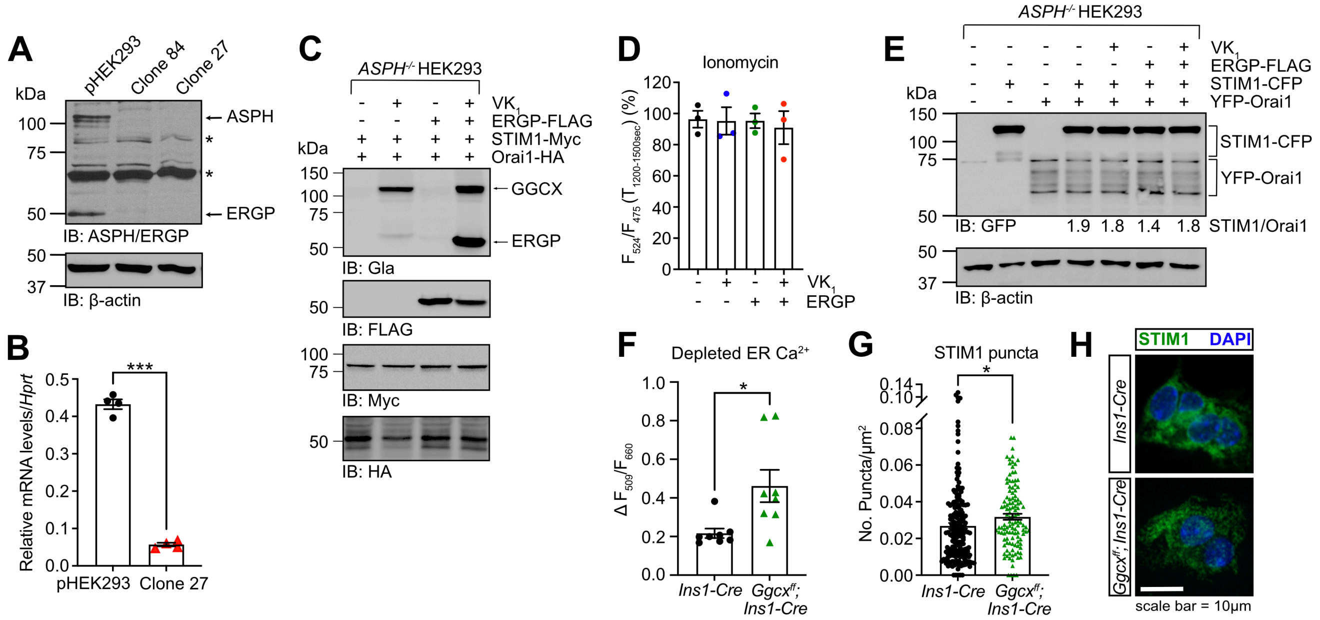


**Figure S7, related to Figure 6 and 7: Gamma-carboxylation in β-cells affects STIM1 puncta formation. (A-B)** Efficient knockout of ASPH and ERGP in parental (p) HEK293 cells was validated by **(A)** western blot and **(B)** qPCR analysis (n=4). Asterisks indicate non-specific binding. **(C)** *ASPH*$^{-/-}$ HEK293 cells were transfected with STIM1-Myc, Orai1-HA and ERGP-3XFLAG in presence or absence of 22µM $VK_1$ as indicated. Expression and γ-carboxylation were monitored by western blot using anti-Myc, anti-HA, anti-FLAG and anti-Gla antibodies. **(D)** Quantification of the average D4ER $F_{524}/F_{475}$ ratio after the addition of 2mM $Ca^{2+}$ and 3µM ionomycin for the last 300 seconds of recording and expressed as % of the baseline vehicle condition. Data are extracted from experiments shown in figure 6F (n=3 experimental replicates per condition, with data from 61-118 cells averaged for each replicate). **(E)** *ASPH*$^{-/-}$ HEK293 cells were transfected with STIM1-CFP, YFP-Orai1 and ERGP-3XFLAG in presence or absence of 22µM $VK_1$ as indicated. Expression of STIM1-CFP and YFP-Orai1 was monitored by western blot using anti-GFP antibodies recognizing both CFP and YFP. β-actin was used as a loading control. Quantification was performed using arbitrary densitometry units of STIM1-CFP signals over YFP-Orai1 signals. **(F)** Average change in $F_{509}/F_{660}$ ratio from baseline (average $T_{0\text{-}120sec}$) to peak after 10µM thapsigargin in *Ggcx*$^{ff}$*; Ins1-Cre* and *Ins1-Cre* islets (n=8 experimental replicates per genotypes, with data from 109-182 islet cell clusters averaged for each replicate). **(G)** Quantification of STIM1 puncta in *Ggcx*$^{ff}$*; Ins1-Cre* and *Ins1-Cre* islets at steady-state (5mM glucose). Data are represented as number of puncta per µm$^2$ (n=116-183 cells). **(H)** Representative confocal immunofluorescence images of *Ggcx*$^{ff}$*; Ins1-Cre* and *Ins1-Cre* islets fixed and stained with anti-STIM1 antibodies. DAPI was used to stain nuclei. Scale bar: 10µm. Results represent the mean ± SEM; Unpaired two-tailed Student's *t* test was used in (B), (F) and (G); Ordinary one-way ANOVA with Bonferroni's post-tests was used in (D); ***$P < 0.001$; *$P < 0.05$.

**Table S1: Donors informations**

| Donor ID | Sex | Age (year) | BMI | HbA1c (%) | Diabetes | Time from diagnosis (year) | Treatment | Figures | Source of Islets |
|---|---|---|---|---|---|---|---|---|---|
| **R266** | F | 74 | 29.2 | 6 | NO | n/a | | Fig. 1G-H | ADI IsletCore |
| **R276** | F | 54 | 24.4 | 7.2 | T2D | 10 | Insulin 1.5y | Fig. 4H-J | ADI IsletCore |
| **R286** | M | 41 | 20.4 | 5.2 | NO | n/a | | Fig. 4H-J | ADI IsletCore |
| **R288** | F | 69 | 27.7 | 5.7 | NO | n/a | | Fig. 4H-J, 5H-I | ADI IsletCore |
| **R292** | M | 47 | 27.6 | 5.6 | NO | n/a | | Fig. 4H-J | ADI IsletCore |
| **R307** | M | 43 | 25.4 | 6.5 | T2D | 2 | Metformin | Fig. 4H-J | ADI IsletCore |
| **R332** | M | 64 | 29.6 | 4.8 | NO | n/a | | Fig. 4H-J | ADI IsletCore |
| **R345** | M | 66 | 29.3 | 6.3 | T2D | 12 | Diet controlled | Fig. 4H-J | ADI IsletCore |
| **R358** | M | 58 | 32.5 | 5.4 | NO | n/a | | Fig. 4H-J | ADI IsletCore |
| **R369** | M | 66 | 25.6 | 4.9 | NO | n/a | | Fig. 4H-J | ADI IsletCore |
| **R382** | M | 45 | 29.7 | 5.5 | NO | n/a | | Fig. 4H-J | ADI IsletCore |
| **R389** | F | 65 | 24.4 | 5.5 | NO | n/a | | Fig. 4H-J | ADI IsletCore |
| **R391** | M | 67 | 24.5 | 4.9 | NO | n/a | | Fig. 4H-J | ADI IsletCore |
| **R399** | M | 47 | 40.3 | 5.5 | NO | n/a | | Fig. 4H-J | ADI IsletCore |
| **R401** | M | 51 | 37.4 | 6.4 | T2D | 10 | Metformin | Fig. 4H-J | ADI IsletCore |
| **R419** | M | 70 | 31.5 | 5.3 | NO | n/a | | Fig. 4H-J | ADI IsletCore |
| **R464** | M | 60 | 27.7 | 5.3 | NO | n/a | | Fig. 5I | ADI IsletCore |
| **HP-22265-01** | M | 67 | 34.4 | 5.3 | NO | n/a | | Fig. 1H | IIDP |
| **HU1252** | M | 52 | 37.5 | 5.3 | NO | n/a | | Fig. 1H, 5I | IIDP |

ADI IsletCore : Alberta Diabetes Institute IsletCore
IDDP : Integrated Islet Distribution Program

**Table S3: Gene expression profile overlap between *Ggcx$^{ff}$; Pdx1-Cre* islets and pre- or diabetic islets.**

| Gene set 1 (gene #) | Gene set 2 (gene #) | Overlap gene nb (%) | Enrichment (fold) | P value |
|---|---|---|---|---|
| ↑ in *Ggcx$^{ff}$; Pdx1-Cre* (79) | ↑ in *Lepr$^{db/db}$* (1278) | 29 (37) | 13.5 | $2.5E^{-25}$ |
| ↓ in *Ggcx$^{ff}$; Pdx1-Cre* (54) | ↓ in *Lepr$^{db/db}$* (1237) | 21 (39) | 14.4 | $1.3E^{-19}$ |
| ↑ in *Ggcx$^{ff}$; Pdx1-Cre* (79) | ↑ in *Ire α$^{β-/-}$* (2285) | 50 (63) | 13 | $1.0E^{-45}$ |
| ↓ in *Ggcx$^{ff}$; Pdx1-Cre* (54) | ↓ in *Ire α$^{β-/-}$* (1852) | 21 (39) | 9.9 | $4.3E^{-16}$ |
| ↑ in *Ggcx$^{ff}$; Pdx1-Cre* (79) | ↑ in HFD (971) | 20 (25) | 12.3 | $1.4E^{-16}$ |
| ↓ in *Ggcx$^{ff}$; Pdx1-Cre* (54) | ↓ in HFD (395) | 6 (11) | 13.2 | $6.2E^{-6}$ |

**Table S4 : LC-MS/MS analyses of WT and *Vkorc1$^{-/-}$* P5 liver extracts following a-Gla IP**

| | Exclusive spectrum count | | | | |
|---|---|---|---|---|---|
| **Identified proteins (gene name)** | **WT #1** | **WT #2** | ***Vkorc1$^{-/-}$* #1** | ***Vkorc1$^{-/-}$* #2** | **Cellular localization** |
| ***Asph*** | 27 | 24 | 0 | 0 | Endoplasmic reticulum |
| ***Ggcx*** | 23 | 23 | 1 | 0 | Endoplasmic reticulum |
| ***F2*** | 4 | 2 | 0 | 0 | Extracellular space |
| ***Pgam5*** | 3 | 3 | 0 | 0 | Mitochondria |
| ***Bdh1*** | 2 | 4 | 0 | 0 | Mitochondria |
| ***Ass1*** | 4 | 1 | 0 | 0 | Cytosol |
| ***Atp5a1*** | 6 | 3 | 1 | 0 | Mitochondria |

**Table S5 : Identification of the LGVYDADGDGDFDVDDAK peptide of ERGP by LC-MS/MS analyses following a-Gla IP in INS-1 832/3 cells.**

| | Identification of LGVYDADGDGDFDVDDAK | | | |
|---|---|---|---|---|
| **Culture conditions** | **a-Gla IP #1** | **a-Gla IP #2** | **a-Gla IP #3** | **Absolute Intensity (average ± SD)** |
| $VK_1$ | yes | yes | yes | $(10 \pm 6.3)$ x$10^4$ |
| $VK_1$ + Thapsigargin | yes | yes | yes | $(5.5 \pm 2.1)$ x$10^4$ |
| Thapsigargin | no | no | no | |
